# Synthesis of geometrically realistic and watertight neuronal ultrastructure manifolds for *in silico* modeling

**DOI:** 10.1101/2024.03.11.584388

**Authors:** Marwan Abdellah, Alessandro Foni, Juan José García Cantero, Nadir Román Guerrero, Elvis Boci, Adrien Fleury, Jay S. Coggan, Daniel Keller, Judit Planas, Jean-Denis Courcol, Georges Khazen

**Affiliations:** Blue Brain Project, École Polytecnique Fédérale de Lausanne (EPFL), Geneva, Switzerland

**Keywords:** Ultrasturcture, mesh reconstruction, surface & solid voxelization, watertight, *in silico*, molecular simulations, reaction-diffusion simulations

## Abstract

Understanding the intracellular dynamics of brain cells entails performing three-dimensional molecular simulations incorporating ultrastructural models that can capture cellular membrane geometries at nanometer scales. While there is an abundance of neuronal morphologies available online, e.g. from NeuroMorpho.Org, converting those fairly abstract point-and-diameter representations into geometrically realistic and simulation-ready, i.e. watertight, manifolds is challenging. Many neuronal mesh reconstruction methods have been proposed, however, their resulting meshes are either biologically unplausible or non-watertight. We present an effective and unconditionally robust method capable of generating geometrically realistic and watertight surface manifolds of spiny cortical neurons from their morphological descriptions. The robustness of our method is assessed based on a mixed dataset of cortical neurons with a wide variety of morphological classes. The implementation is seamlessly extended and applied to synthetic astrocytic morphologies that are also plausibly biological in detail. Resulting meshes are ultimately used to create volumetric meshes with tetrahedral domains to perform scalable *in silico* reaction-diffusion simulations for revealing cellular structure–function relationships.

**Availability and implementation:** Our method is implemented in NeuroMorphoVis, a neuroscience-specific open source Blender add-on, making it freely accessible for neuroscience researchers.

**Key points:**

- A plethora of neuronal morphologies is available in a point-and-diameter format, but there are no robust techniques capable of converting these morphologies into geometrically realistic models that can be used to conduct subcellular simulations.
- We present a scalable method capable of synthesizing high fidelity watertight ultrastructural manifolds of complete neuronal models from their one-dimensional descriptions using the synaptic data obtained from the digitally reconstructed neuronal circuits of the Blue Brain Project.
- Resulting manifold models comprise geometrically realistic somata and spine geometries, enabling accurate *in silico* experiments that can probe intricate structure-function relationships.
- Our method is extensible and can be seamlessly applied to other cellular structures such as astroglial morphologies and even large networks of cerebral vasculature.

## 1 Introduction

Revealing the underlying mechanisms governing brain function requires an in-depth understanding of cellular and network dynamics at multiple levels of detail, with significant biological computations also taking place in very small volumes within critically defined ultrastructures^1,2^. Although wet lab experiments, such as *in vivo* and *in vitro* studies, remain essential in neuroscience, the use of computer simulation-based, or *in silico*, experiments complements the research cycle^3^, allowing certain observations that are unattainable through traditional methods. Particularly at the cellular and subcellular levels, this complementary approach entails employing high performance simulations that utilize biophysically plausible and highly detailed structural models of brain cells, such as neurons and glial cells^4^, in which cellular kinetics can be simulated to characterize structure– function relationships. Simulations have been performed previously using reduced (one-dimensional) compartmental models employing the Neuron simulator^5^ to compute electrophysiological responses on a cellular level. Those simulations range from single cells up to large networks of digitally reconstructed circuits^6^. Simulation scalability was attainable relying on an optimized version of the Neuron simulator called CoreNeuron^7^. Nevertheless, these simulations neglected the precise physical structure of the neuron and therefore could not be used to capture the dynamics of subcellular scales. Reaction-diffusion models were introduced to address this challenge^8^, by incorporating detailed cellular morphologies with geometrically realistic structures into three-dimensional (3D) simulations with which we can realize complex signaling pathways in the brain in space and time^9–11^. While there has been extensive ongoing research focused on improving the performance of reaction-diffusion simulators to run efficiently on large scale supercomputing architectures, for example with STEPS 4.0^12^, there is still a large unfilled gap in the generation of those ultrastructural data models needed to conduct the simulations. Our method is introduced to fill in this gap, enabling computational neuroscientists to automate the generation of brain tissue models with realistic cellular boundaries to conduct high performance molecular simulations.

### 1.1 Neuronal models and the watertightness challenge

Reaction-diffusion simulations are principally applied to 3D volumetric meshes that are subdivided into tetrahedral subdomains where molecular interactions can be contained^12^. Such tetrahedral meshes are ordinarily created from corresponding surface counterparts, for example using TetGen^13^. However, this process requires the input surface mesh to be a watertight manifold, i.e. two-manifold (with zero non-manifold edges and vertices) with no self-intersecting facets (refer to Supplementary Section 1 and Supplementary Fig. S1). While TetWild^14^ is developed specifically to handle non-watertight meshes, it has impractical performance and cannot process thin structures. The watertightness requirement is relatively challenging to attain for brain cells, particularly for neuronal morphologies that are characterized by complex arborizations with thin fibers, irregular branching geometries, large spatial extents and low volume occupancy.

Neuronal cells are commonly available from open neuroscience databases, such as NeuroMorpho.Org, in a hierarchical structure with point-and-diameter representations^15^, which further complicates the generation of biologically realistic models that can be plugged directly into the simulation. Morphological models of biological neurons are typically segmented from optical, tomography or electron microscopy stacks^16,17^. Segmentation can be manual, semi-automated or even fully automated^18^. Moreover, biologically inspired neuronal models are becoming more available thanks to recent studies that utilize biophysical simulations to synthesize neurons relying on advanced protocols derived from the analysis of realistic counterparts^19^. After their segmentation or synthesis, morphologies are restructured into a conventional hierarchical representation based on a connected set of digitized samples. This representation uses directed acyclic graphs (DAGs) to store and build the connectivity between those samples where the root node represents the cell body (or soma) and the leaf nodes represent the terminal samples. Neuronal skeletons are composed of a set of morphological samples; each pair of adjacent samples defines a segment and a set of connected segments between two branching samples defines a section. The arbors, or neurites, are composed of a set of connected sections in a tree structure (Figure 1). Converting this structure into a continuous watertight manifold in a reliable manner is missing.

**Figure 1.**
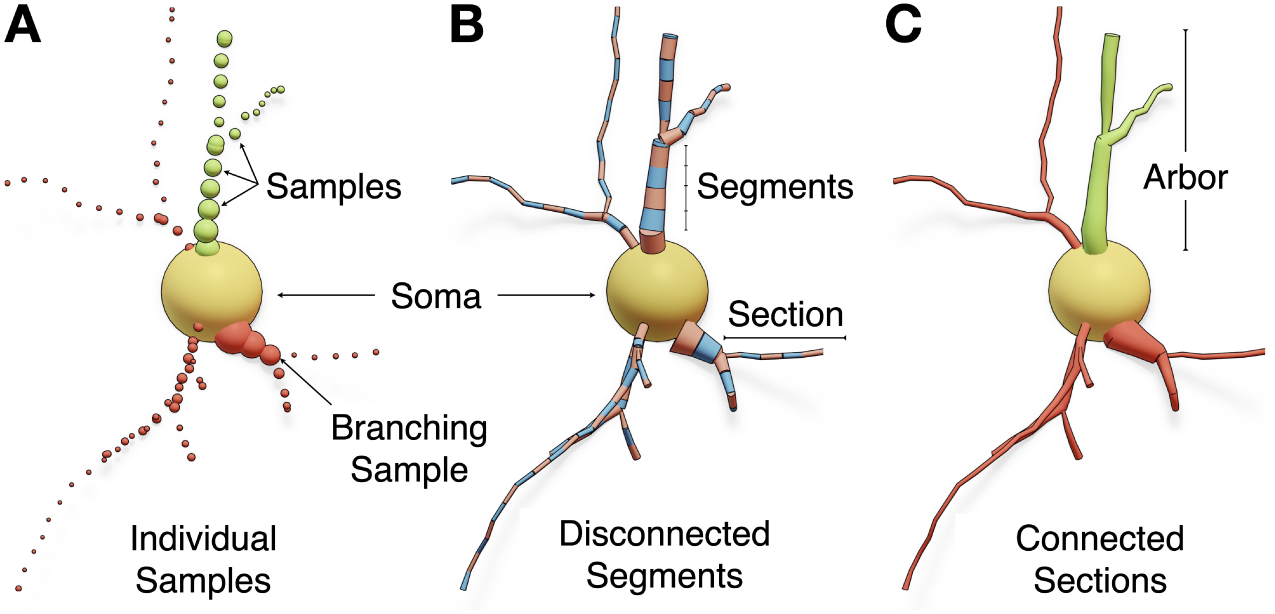
The structure of a neuronal morphology in three formats: (A) *individual samples*, where each sample has a unique identifier, position and diameter, (B) *segments*, in which each pair of connected samples form conical segments and (C) *sections*, where adjacent segments between two branching points construct an individual section. The arbors are composed of a set of connected sections and stored in acyclic graphs. Dendritic spines are not shown.

### 1.2 Related Work

Reconstructing surface meshes of neurons from fairly abstract graph representation has been investigated in several interdisciplinary studies either in computer graphics and visual computing or in bioinformatics. The majority of those studies focused on designing meshing algorithms capable of creating visually appealing — and more importantly lightweight — mesh models that could be applied in domain-specific visual analytics applications, e.g., to visualize the electrophysiological activity^20–22^ of digitally reconstructed neuronal networks^6^ using the compartmental reports generated with Neuron^5^. Those algorithms were primarily concerned with the tessellation of the resulting models, i.e., creating appearance-preserving meshes with minimal polygon counts, allowing the visualization of compartmental simulations of large-scale circuits containing several hundreds of thousands or millions of neurons^23,24^. Some approaches presented advanced solutions to improve the realism of the resulting models with different objectives: (i) to construct faithful 3D somatic profiles using physics simulations, e.g., Neuronize^25^, NeuroTessMesh^26^ and NeuroMorphoVis^27,28^, as opposed to earlier applications that used simplified primitives such as spheres^29^ and cylinders^18^ to model the soma; (ii) to create smooth and accurate branching geometries along the neuronal arbors using skin modifiers^22^ and Laplacian smoothing with Boolean unions^30^; and (iii) to integrate geometrically realistic spine models (extracted from electron microscopy) along the dendrites of the neurons^31^. However, watertightness was lacking and the usability of the resulting neuronal meshes was consequently limited either to visual analytics or content creations purposes.

On the contrary, and until recently, the watertightness aspect in neuronal mesh generation was only accomplished in two principal studies^32,33^. McDougal et al. developed the constructive tessellated neuronal geometry (CTNG) algorithm to create a continuous intermediate representation of the cellular plasma membrane, with which an extended version of the marching cubes algorithm (called constructive marching cubes) is applied to convert this intermediate surface into a watertight manifold consistent with the neuronal morphology^32^. This CTNG algorithm uses a complex set of geometric subroutines to fill the gaps, remove the overlaps and extrusions between the consecutive segments of the arbors and along the connections between the soma and first-order sections. The CTNG algorithm was coded in a combination of C, Python and Cython, and the implementation was open sourced on ModelDB, but the code is deprecated and can no longer be used with the recent versions of Python. Therefore we were unable to assess either its performance or the quality of its resulting meshes.

Mörschel et al. developed AnaMorph, another domain-specific solution tailored to create watertight neuronal meshes from SWC morphologies using non-linear piecewise analytical modeling and union operators^33^. AnaMorph is publicly available under LGPL license, the code is entirely implemented in modern C++ and can be easily compiled on Unix-based operating systems. Nonetheless, this solution was limited in the following aspects: (i) it did not account for a realistic somatic profile since somata were approximated by spheres; (ii) the implementation fails if local self-intersections exist; (iii) the volume of the resulting mesh is significantly lower than the actual volume of the morphology; and finally (iv) the algorithm was incapable of incorporating spine models with realistic shapes into the final meshes but rather approximated them with cylinders.

To address this challenge, we present an efficient, unique and accessible solution that can combine the aspects of realism and watertightness in a straightforward way, without requiring any complex geometric operations. This approach enables the creation of a continuous and smooth surface mesh reflecting the plasma membrane of a neuron from its graph.

## 2 Contributions

1. Intuitive and unconditionally robust algorithm for synthesizing geometrically realistic, watertight and optimized manifolds of spiny neurons from their morphological traces, making it possible to improve the biological accuracy of reaction-diffusion simulations.
2. Efficient implementation of the algorithm using the modeling toolsets of Blenderand its multi-threaded Voxel remesher.
3. Integrating the implementation into the Meshing Toolbox of the NeuroMorphoVis add-on^28^ and exposing the functionality to users from the graphical user interface (GUI) of Blender and the command line interface (CLI) of the add-on.
4. Applying the implementation to thousands of neurons with different morphological classes^6^ and evaluating the quantitative and qualitative aspects of the resulting meshes.
5. Extending the implementation to create watertight manifolds of synthetic astroglial morphologies with realistic endfeet data^34^.

## 3 Methods

We present an effective method capable of synthesizing a smooth watertight surface manifold that can model the plasma membrane of a spiny neuron from its abstract skeletal representation. Our algorithm consists of three principal stages. The first builds individual, overlapping, and non-watertight proxy meshes of the different components of the neuron, which are nonetheless **geometrically realistic**. These include arbors, somata, and spines. The second stage assembles those proxies into a joint mesh object that is rasterized and polygonized to yield an intermediate coarse watertight mesh of the neuron. This stage is implemented exclusively relying on the Voxel remeshing modifier that has been incorporated into the recent versions of Blender. The last stage optimizes the intermediate mesh to generate a **volume-preserved, watertight** and continuous surface manifold of the neuronal membrane. To complete the pipeline, the resulting mesh is then used by TetGen^13^ to create a compartmental volume grid with tetrahedral subdomains for performing stochastic reaction-diffusion simulations, mainly in STEPS^12^. The pipeline is graphically illustrated, end-to-end, in Figure 2. The specific details of the tetrahedralization and simulation protocols are beyond the scope of this work.

**Figure 2.**
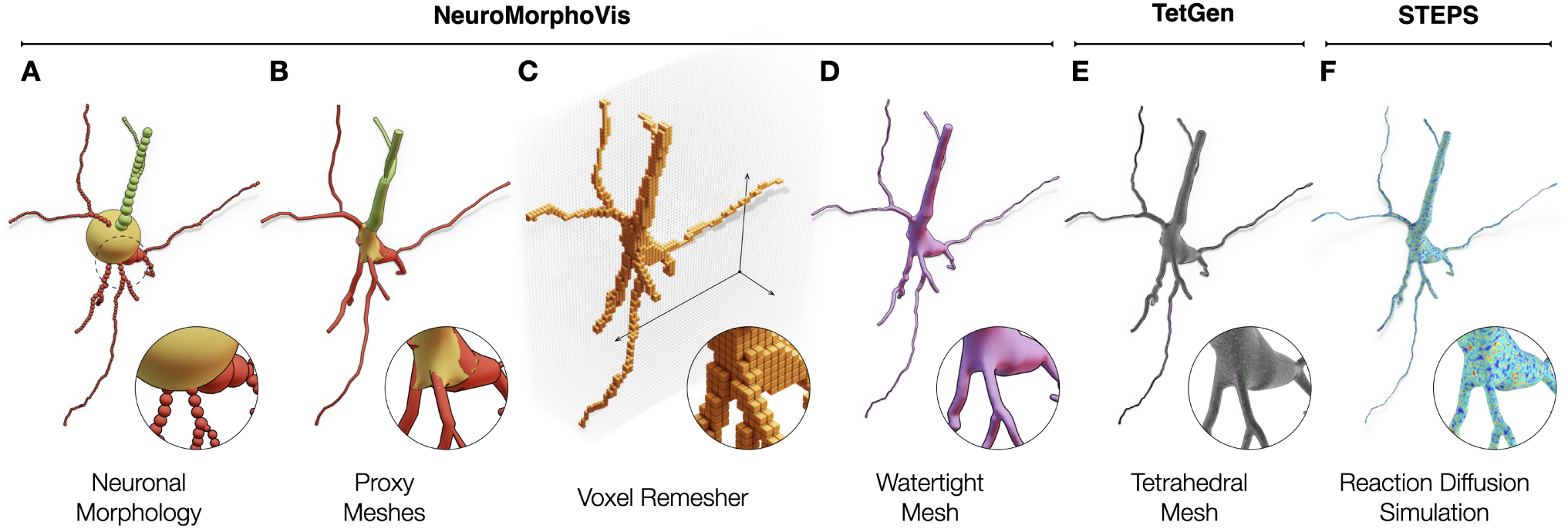
The neuronal morphology (A) is initially used to create a set of corresponding proxy meshes of every individual component of the morphology, which are then combined into a single mesh object with overlapping geometries using a joint operation (B). The Voxel remesher is applied to this mesh object to create a volumetric representation of the membrane (C) with which all the overlapping structures are eliminated. This remesher creates a watertight manifold with a continuous and smooth surface (D), which can be used to synthesize a volumetric mesh (E), for example using TetGen, to perform a stochastic reaction-diffusion simulation in STEPS (F). Dendritic spines are not shown.

### 3.1 Morphology preprocessing

Biological neuronal morphologies extracted from tissue samples are typically traced and digitized in a manual or semi-automated manner. This reconstruction process can be accompanied by various patterns of artifacts either due to the staining procedure itself or due to other manual errors introduced by the operator. A set of predefined processing operations are therefore applied to the input morphology to assert the elimination of any skeletal artifacts that might lead to subsequent geometric deficits with potential impact on the simulation results. However, these operations do not change the structure of the morphology, such as the connectivity between the branches. This process includes: (i) verification of the connectivity of the emanating arbors from the soma; (ii) removal of the morphological samples that are located within the somatic spatial extent; and (iii) adaptive resampling of the morphological sections to eliminate the high frequency perturbations along the surface of the resulting mesh.

### 3.2 Generation of proxy arbors

Proxies of neuronal arbors are generated using one of the two following algorithms: (i) node-to-leaf path construction or (ii) articulated sections. The first algorithm uses depth-first traversal to construct, per neurite, a set of paths starting from the root node of the morphology graph (first-order sections) to its leaf nodes (terminal sections). Before the construction of the paths, each section in the morphology is labeled as either primary or secondary, for the purpose of deciding at the respective branching points which child section forms the most natural continuation along the path. This labeling scheme is adopted in previous methods^21,30^ to form piecewise linear paths on a per-section-basis, using cubic splines to form the path.

The articulated sections algorithm uses geodesic polyhedra – or icospheres – to connect between the different sections of the morphological hierarchy. The diameter of each icosphere is calculated based on the largest sample at each respective branching point; this guarantees a seamless continuation from parent to child sections. Unlike the first method, the samples of each section are used to build an independent path that has no connectivity with parent or child sections. This connection is maintained by the articulation icospheres. The difference between the two methods is illustrated in Figure 3. In both cases, and after the construction of the linear paths, a circular cross-sectional ring is used to interpolate those paths and construct a set of tubular proxy geometries, which can be rasterized by the Voxel remesher.

**Figure 3.**
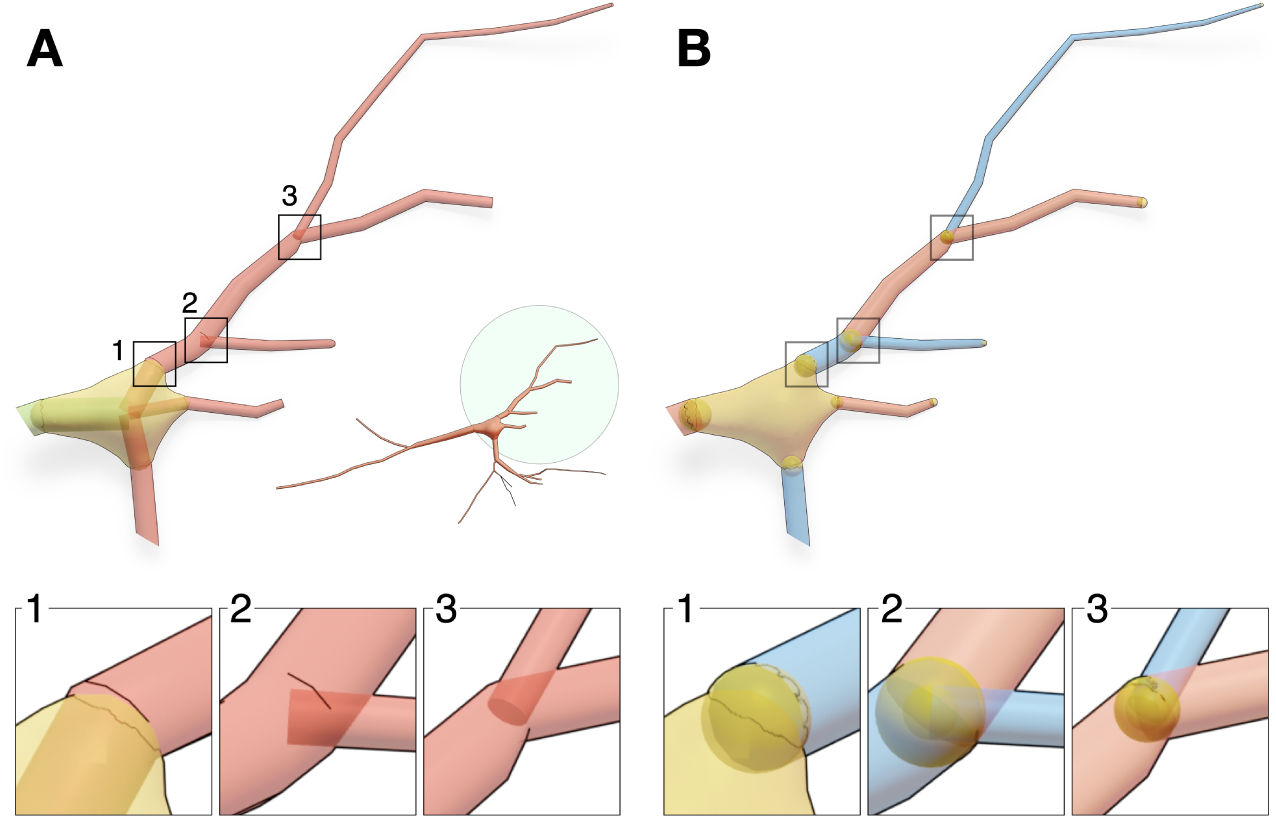
Two approaches are used to construct the proxies of the arbors: node-to-leaf path constructions (A) and articulated sections, where an icosphere (or geodesic polyhedra) is added at every branching point along the arbor (B).

### 3.3 Reconstruction of realistic somatic profiles

Due to certain limitations in the acquisition process, the soma is identified in the majority of biological reconstructions with a sphere, whose radius approximates the distance between the local centroid of the soma and the initial segments of all the emanating branches. More advanced reconstructions integrate a two-dimensional somatic profile reflecting the projection of the soma along the optical axis of the microscope. Modeling somata with primitive spheres remains a limitation, particularly when the resulting model is employed within the context of a geometrically realistic simulation, where structure impacts the function. Implicit surfaces have been used to reconstruct approximate somatic profiles (Figure 4A) better than spheres^30,35^. However, connecting the arbors to the reconstructed shapes requires a brute force approach such as a boolean operation or a complex geometric algorithm in which arbors can be bridged to emanate smoothly from the soma. A recent Blender-based implementation^27^ incorporated mass-spring models and Hooke’s law to simulate the somatic growth in a physically plausible manner. This implementation can reconstruct faithful somatic profiles (Figure 4B) starting from an icosphere that is expanding towards the initial segments of all the arbors that are verified in the pre-processing stage to be directly connected to the soma. The final shape of the soma depends on three parameters: (i) the initial radius of the sphere used in the simulation; (ii) its stiffness; and (iii) the number of time steps used in the simulation as depicted in (Figure 4C). As elaborated earlier and shown in Figure 3, the continuity between the soma and the arbors at their respective initial segments is guaranteed either by adding auxiliary segments that connect the initial sample of each arbor to the center of the soma (Figure 3A1) or by adding auxiliary spheres (Figure 3B1). The principal advantage of this technique is that somata are, intuitively, integrated in the mesh — there is no need to use boolean or bridging operators to weld the somatic mesh with the arbors. Contrary to the Union operators approach^30^, the Voxel remesher can seamlessly handle this problem and guarantee the resulting manifold to be continuous and watertight.

**Figure 4.**
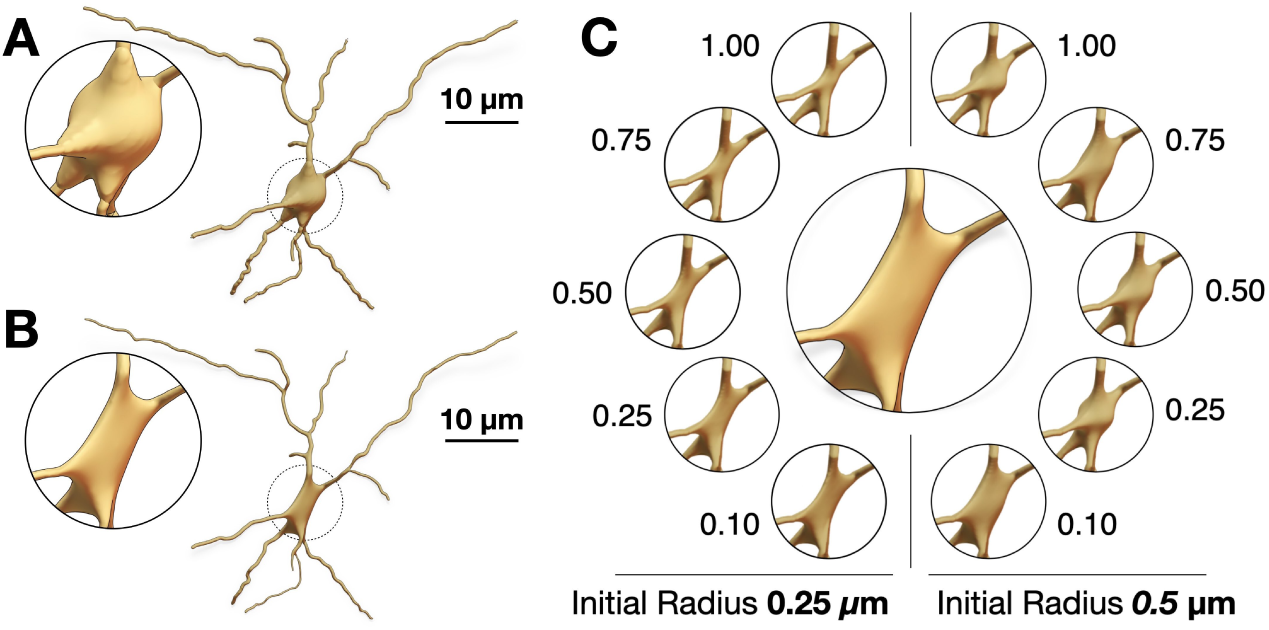
Creation of somatic profiles using implicit surfaces (A) and soft body simulations (B). The realism of the resulting profile using the soft body approach requires tuning the stiffness of the soft body object – indicated on the side of every simulation – and the initial radius of the ico-sphere used to build the mesh (C).

### 3.4 Integrating geometrically realistic spines

Spines are those tiny bulbous protruding structures (1-3 µm in length) distributed along neuronal dendrites, whose function is to receive excitatory synaptic inputs from pre-synaptic neurons to generate a compartmentalized post-synaptic response^36^. Traces of individual neuronal morphologies, for example those that are available from NeuroMorpho.Org, do not comprise any relevant information on the spines; the traces are typically acquired using optical microscopy with limited lateral resolution, making it difficult to reconstruct spine morphologies with enough detail^37^. Electron microscopy (EM) is used in relevant studies^4^ to visualize the neuron ultrastructure, allowing the reconstruction of detailed morphologies of dendritic spines with realistic geometries at nanometer resolutions. Those EM neuronal reconstructions are used to identify and segment a set of dendritic spine geometries that can be used to improve the realism of the resulting neuronal models as opposed to using cylinders to represent the spines^38^.

Our implementation integrates those spine models (Supplementary Fig. S6) only along the membranes of neuronal morphologies that are part of a digitally reconstructed cortical circuit^6^, where we can identify spine types, dimensions, locations and correct orientations. In a similar study, spine models were also integrated along the dendritic shafts of neurons using union operators^30^, but the resulting meshes were not watertight. Our proposed method takes advantage of having accurate and smooth connection between the dendritic membrane and the spine without performing any geometric operations that are error-prone and might fail if the topology of the mesh is not good as shown in Figure 5.

**Figure 5.**
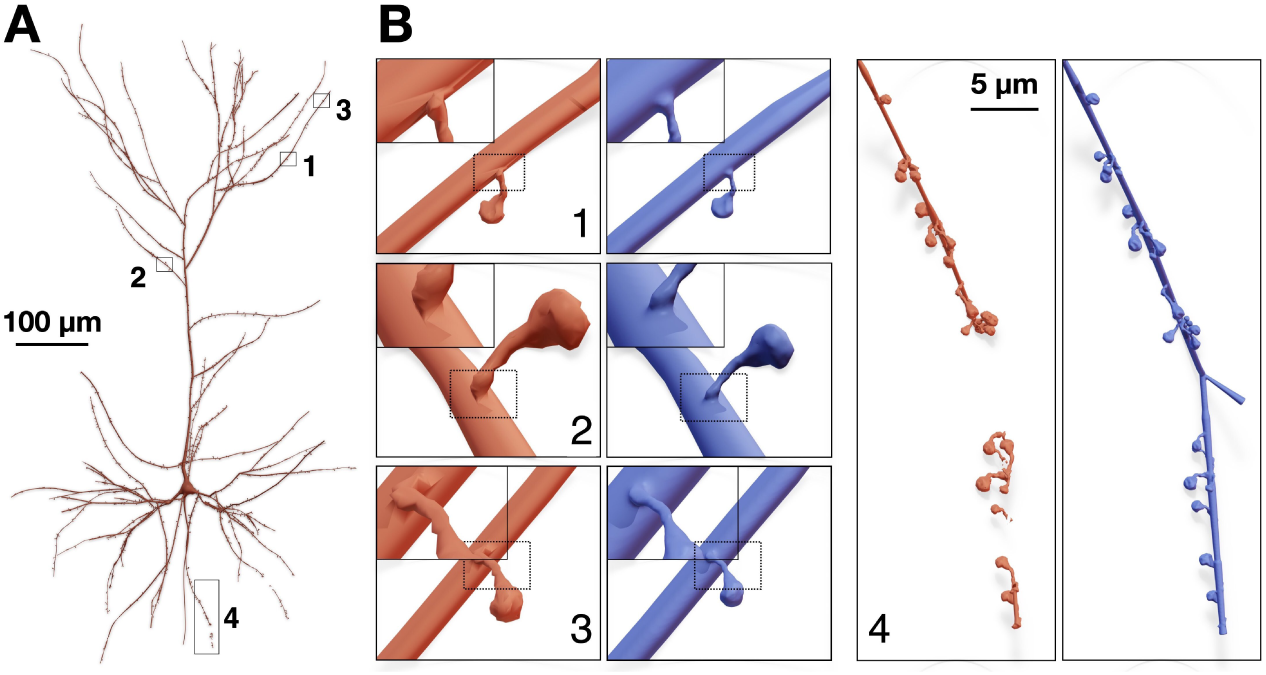
Integration of spines models with realistic geometries along the dendrites of a pyramidal neuron (A). The mesh in light red is created using union operators as described in^30^, while that in blue is created with our proposed method. The closeups in (B1–3) demonstrate the smooth connectivity between the spines and the dendrites. Wireframe visualizations are shown in Fig. 8. (B4) The union operator fails to weld the spine meshes with the dendritic mesh resulting in a fragmented mesh.

### 3.5 The Voxel remesher

After the generation of the proxies corresponding to soma, arbors and spines, a joint operator is applied to create a single proxy mesh. This mesh is guaranteed to have no geometric gaps, but it indeed has self-intersecting facets, particularly at the branching points, at the junctions between the soma and the emanating arbors, and between the dendritic membranes and the spines, as illustrated by the closeup in Figure 2B. The Voxel remesher is then applied to the joint proxy mesh to create an intermediate uniformly sampled volumetric representation of the neuron as showin in Figure 2C. Logically, and while it preserves the structure of the membrane, this conversion into a voxel-based model removes all the self-intersections of the proxy mesh. This volume is then used to create a watertight surface manifold (Figure 2D) using an advanced implementation of the marching cubes algorithm. To capture all the geometric details of the neuron, the voxelization resolution is defined based on the size of the smallest morphological sample in the morphology and the thinnest cross-section of the smallest spine. Using a greater value might lead to creating a fragmented mesh as shown in Figure 6. Therefore a post-processing operation is applied to verify whether the final mesh has a single continuous manifold — that is essential for the simulation — or is composed of multiple disconnected partitions. One main advantage of the Voxel remesher is its performance; it has a multi-threaded implementation, contrary to the metaball polygonization modifier.

**Figure 6.**
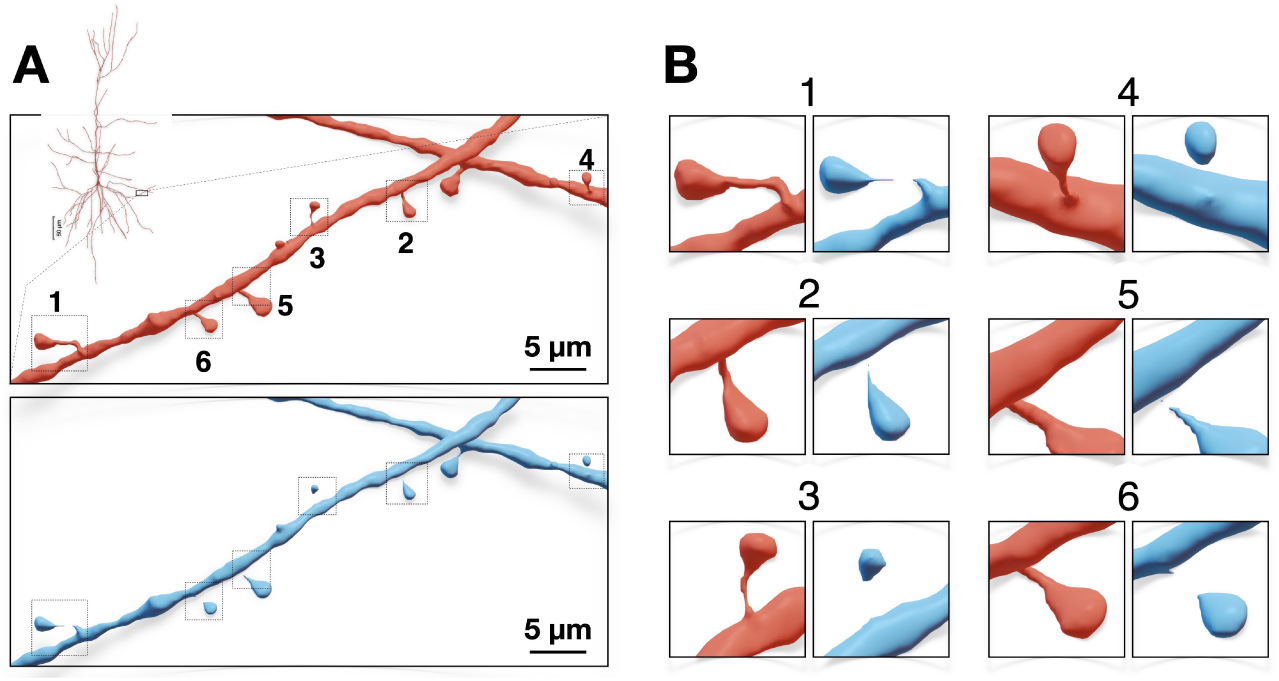
(A) A closeup on a dendritic segment of a spiny neuron showing the resulting meshes with different voxelization resolutions: 0.07 and 0.1 *µ*m for the red and blue meshes respectively. Using lower resolution without taking into consideration the dimensions of the spine meshes results in fragmented mesh partitions as demonstrated by the magnifications in (B).

### 3.6 Mesh optimization & watertightness verification

As shown in Figure 6, and in order to capture the finest structure of the neuron – mainly the extrusions of the spines from the apical dendrites, the Voxel remesher is applied with a decent resolution that is sufficient to reconstruct the ultrastructural geometric details of the smallest spine in the final mesh. This leads to creating over-tessellated surface manifolds with several millions or even tens of millions of facets, which limits the potential usability of the resulting meshes in some simulation applications. To address this challenge, we use adaptive geometry-preserving mesh optimization to decimate the surface of the intermediate mesh, as much as possible, while preserving the geometric aspects of the morphology (Figs. 7 & 8). The mesh optimization procedure includes surface coarsening, iterative face and normal smoothing. We implemented an extended and efficient library called OMesh (or OptimizationMesh) with dedicated Python bindings that can be seamlessly integrated into Blender to optimize the meshes produced by the Voxel remesher within the same context.

**Figure 7.**
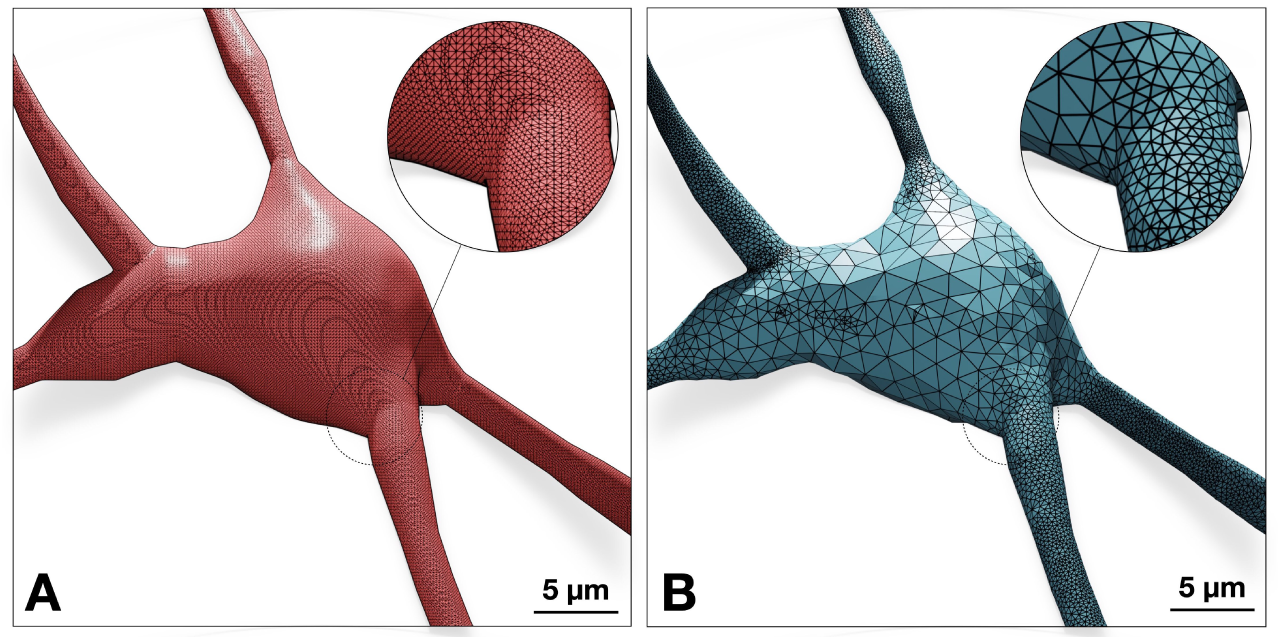
The neuronal mesh generated from the Voxel remesher (left) is typically highly tessellated (*∼*100k triangles). This mesh is re-tessellated using coarsening to create an adaptively optimized clone (right) – with *∼*68k triangles, where local regions with high frequency contain more facets than flat regions. Complete analysis of both meshes is illustrated Fig. 9.

**Figure 8.**
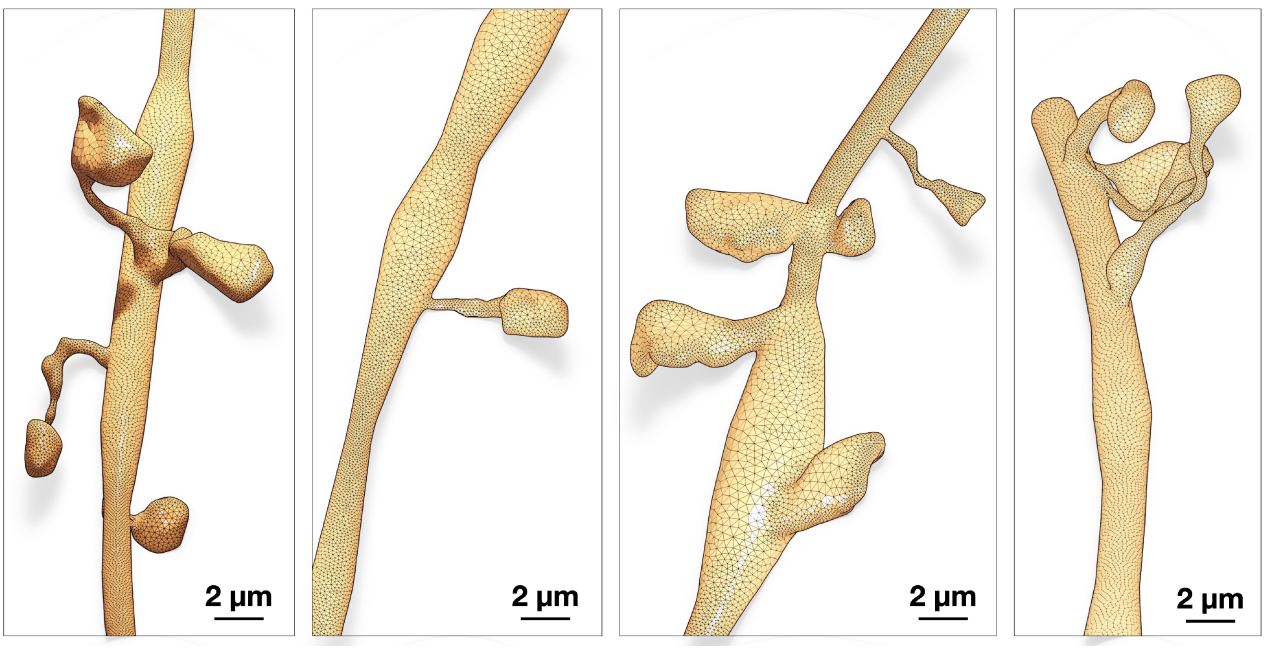
While adaptive optimization eliminates unnecessary vertices of flat regions of the manifold – mainly across the somatic region as shown in Fig. 7, the topology of the mesh around spines still have sufficient number of vertices to capture their geometric details.

Mesh coarsening impacts the watertightness of the mesh by introducing self-intersecting facets along its surface^39^. While iterative smoothing reduces the number of self-intersecting facets, it cannot guarantee the complete elimination of all the self-intersections produced in the coarsening stage (as shown in Supplementary Fig. S5), which makes our algorithm unrobust. We resolved this issue by implementing an effective iterative watertightness verification scheme that detects any elements (non-manifold edges, non-manifold vertices, thin-faces, zero-faces, or self-intersections) that might affect the watertightness of the mesh. The vertices corresponding to those elements are marked for elimination, and a re-triangulation operation is then applied to close the holes introduced across the surface of the mesh. This procedure is detailed in Supplementary Section 4.

### 3.7 Implementation

Our algorithm is implemented in Blender 3.5 using its embedded Python interpreter and API modules. The implementation is integrated to the Meshing Toolbox of the NeuroMorphoVis^28^ add-on. The functionality is made accessible to users, primarily computational neurobiologists and neuroanatomists, from the GUI of Blender (for single cell analysis and mesh generation) and also from the CLI of the add-on (for executing batch jobs of neuron groups). The implementation takes advantage of the capabilities of NeuroMorphoVis to generate the meshes in parallel on a computing cluster using the Slurm scheduler.

## 4 Results and discussion

Reliability and performance of our technique are assessed by applying the meshing implementation to a set of 60 classes of various types of cortical morphologies, each class has 100 neurons. Those exemplary neurons are randomly sampled from a digitally simulated cortical circuit^6,40^ containing 4.2 million biophysically detailed compartmental neurons and 13.4 billion synapses covering 8 cortical subregions. Figure 10 shows a collective collage assembling a set of 60 meshes, each representing a distinct morphological type (Supplementary Table S1). Resulting meshes are exported into STL file format to be compatible with TetGen. All the meshes are verified to have continuous watertight manifolds and high quality qualitative distributions (refer to fact sheets in the Supplementary Figures S7–S126).

### 4.1 Meshing qualitative and quantitative analysis

While watertightness and adaptive mesh refinement are principal objectives of the presented work, high quality geometric measures are still necessary to accomplish — they significantly impact the accuracy of the simulation results. The Verdict library^41^ provides a set of metrics with which geometric qualities of a triangulated surface mesh can be evaluated. This set includes radius ratio, edge ratio, radius to edge ratio, minimum and maximum dihedral angles. To make a complete analysis of the resulting meshes, a collective fact sheet that combines qualitative and the quantitative measures is created. This fact sheet provides comparative results of the intermediate mesh generated with the Voxel remesher and the optimized one that is used for the simulation. Figure 9 demonstrates a comparative analysis fact sheet of the meshes created from a neuron with L6_SBC type. The analysis of the meshes of the other morphological types is provides in Supplementary Figures S7–S126. This analysis includes wireframe visualizations of each mesh to highlight the difference in tessellation between the intermediate and final meshes.

**Figure 9.**
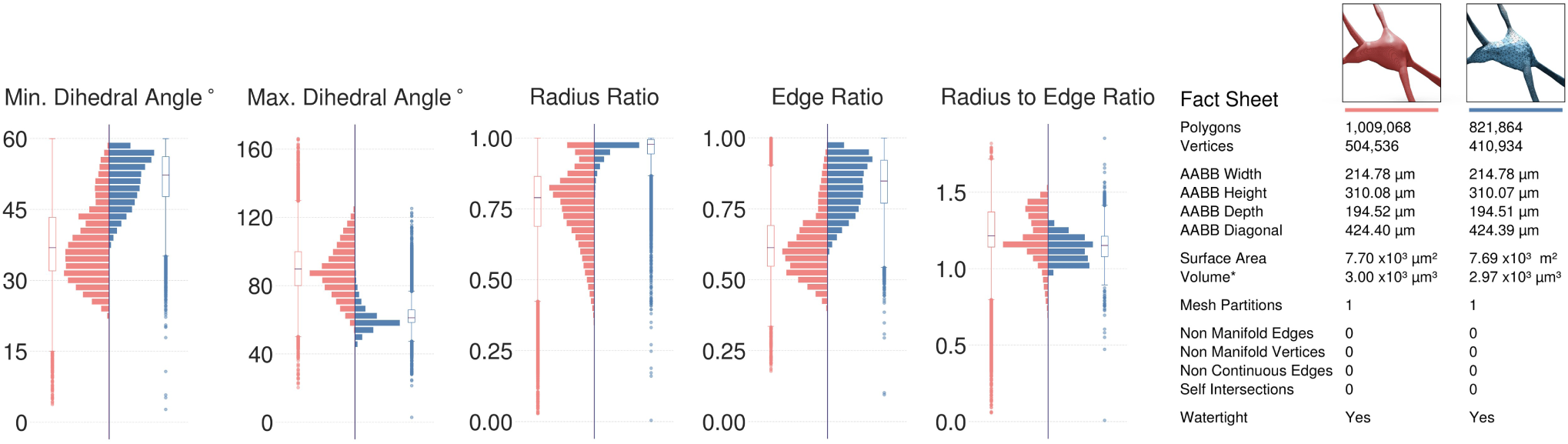
Comparative quantitative and qualitative analyses of the meshes generated from the Voxel remesher (in light red) and the mesh optimizer (in light blue) shown in Fig. 7. Note that the volume of the mesh is preserved.

### 4.2 Performance analysis

The theoretical performance of our algorithm depends on the following factors: (i) the arborization complexity of the morphology, in terms of its maximum branching order and the total number of segments per section; (ii) the total count of dendritic spines in the morphology; (iii) the spatial extent, or the bounding box of the morphology; (iv) the size of the finest structure in the morphology; and (v) the complexity (number of triangles) of the intermediate mesh generated from the Voxel remesher. The arborization complexity determines the tessellation (or number of polygons) of the proxy arbors. While the generation time of the proxies is relatively fast, the more those proxies are tessellated, the longer it takes to rasterize the polygons during the application of the Voxel remesher on the joint proxy. Moreover, and since we use geometrically realistic morphologies for the dendritic spines, having more spines along the dendrites of the neuron will reduce the performance of the joint operation that merges all the proxies into a single mesh object and will similarly impact the performance of the Voxel remesher. The resolution of the volume grid used to voxelize the joint proxy is determined based on the smallest structure in the morphology; either the diameter of the smallest segment (typically 0.10 *µ*m) in the morphology, if the neuron is aspiny (without spines), or the size of the finest spine (*∼*0.06 - 0.80 *µ*m). The optimization procedure is iterative. Depending on the presence of any self-intersections (due to sharp edges or extremely thin faces), the mesh will be subject to a new watertightness verification loop. Therefore, the timing of the optimization procedure cannot be estimated (Section 4 in the Supplementary Document).

We assessed the overall performance of our implementation using 60 exemplar neurons, each one represents a distinct morphological type. The benchmarks, shown in Figure 10, highlight the performance variations for the three stages of the workflow: proxies generation, Voxel remeshing and mesh optimization. The pre-processing stage is relatively negligible, proxies generation and Voxel remeshing (with multi-threaded implementation) take on average *∼*10-12 seconds. The optimization procedure is executed on a per-vertex basis to evaluate if a specific vertex can be deleted from the mesh. This local operation requires querying the neighboring vertices, which make this process unparallizable. Therefore, it takes a few hundreds of seconds to complete.

**Figure 10.**
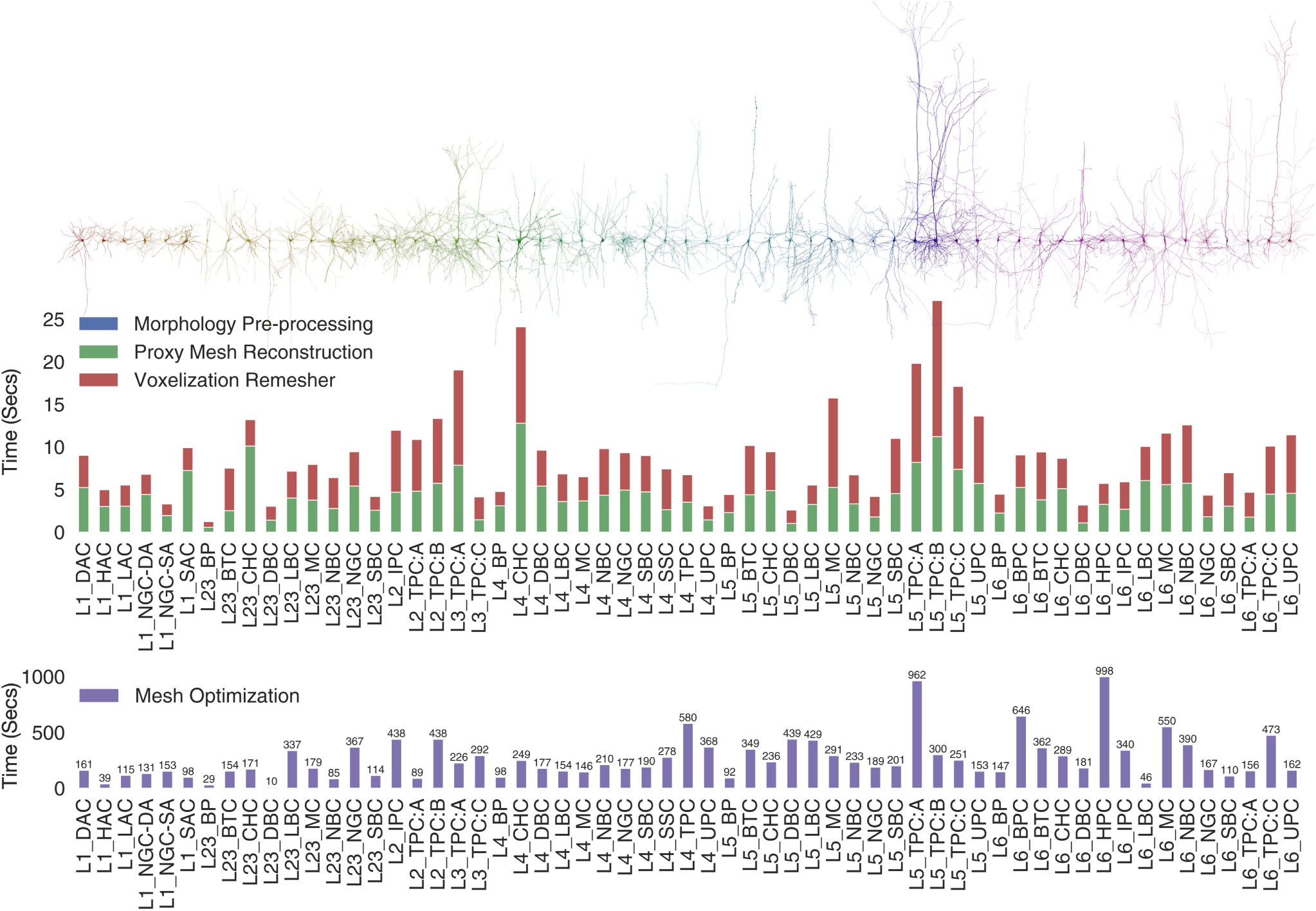
Performance benchmarks (in seconds) for our implemenation based on a data set consisting of 60 various morphological types of cortical neurons^6,40^. The timing of the Proxy Mesh Reconstruction stage comprises the soma simulation time, arbors reconstruction and the integration of the spines along the dendritic arbors. The mesh optimization time comprises the mesh coarsening and smoothing. Axons of the shown neurons are limited to second-order branching only.

These benchmarks are measured on a commodity compute node shipped with 32 GBytes of memory and an Intel core i7-8700 CPU running at 3.2 GHz. These specifications are sufficient to process and mesh large pyramidal neurons that have tens of thousands of dendritic spines. A comparative performance analysis between our Voxel-based remeshing implementation and other meshing algorithms that are exclusively implemented in Blender is thoroughly discussed in the Supplementary document (Section 7).

### 4.3 Application to astrocytic morphologies

Astrocytic morphologies have similar branching structure to neurons, but they contain further endfeet processes wrapped around the vessels of cerebral vasculature to transmit energy to the neurons in the NGV tripartite^4^. A recent implementation extended NeuroMorphoVis to load and visualize astrocytic morphologies^34^ to create corresponding polygonal mesh models using implicit surfaces polygonization^35^. Nonetheless, resulting meshes were not watertight and further post-processing was necessary to ensure watertightness. In comparison, our implementation is fit-for-purpose extended and applied to astrocytic morphologies to create watertight meshes in a single step. Figure 11 illustrates a synthetic astrocyte morphology and its corresponding watertight mesh created with our technique.

**Figure 11.**
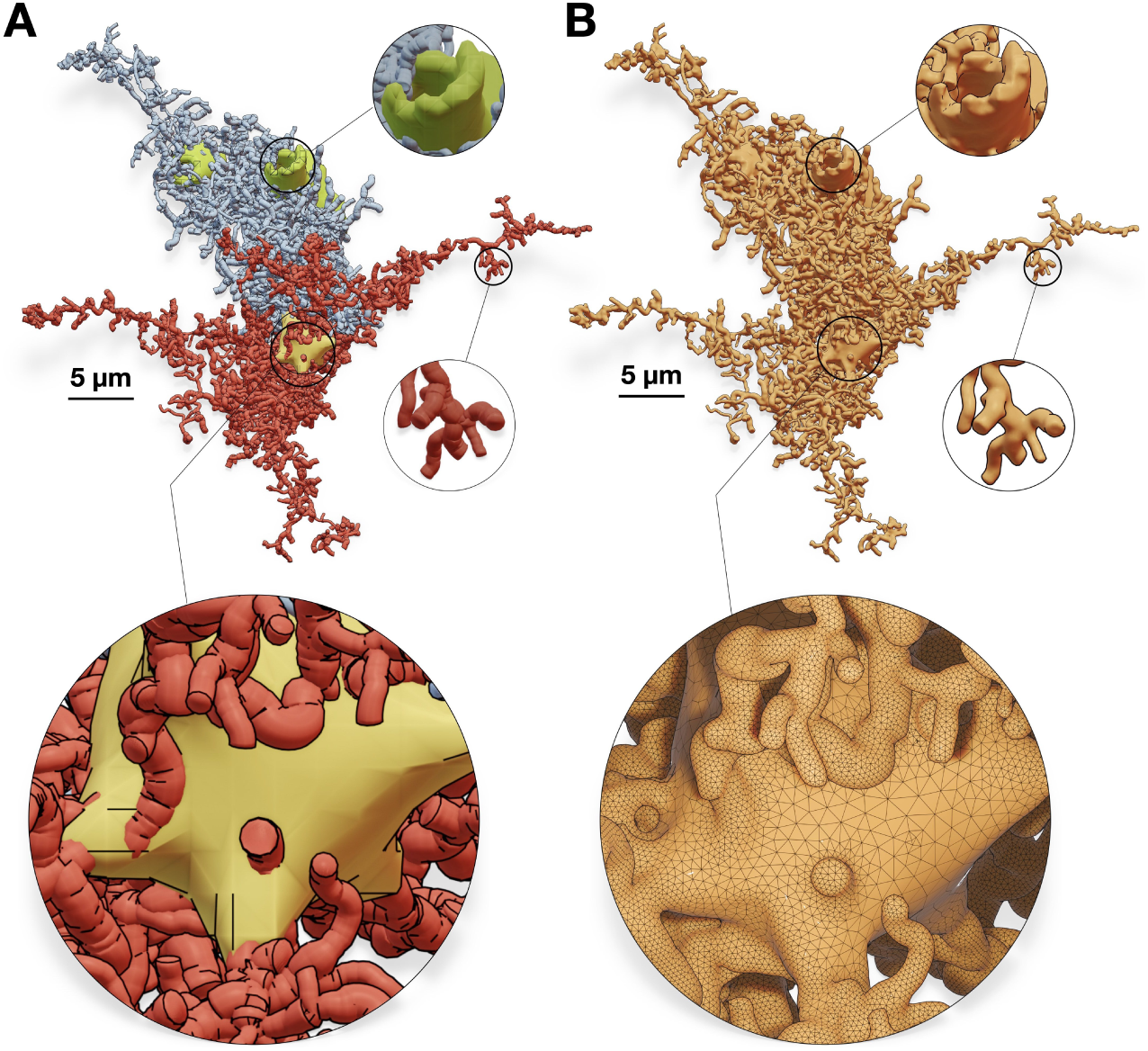
Our algorithm is applied to a synthetic astroglial cell^34^ (A) to create a corresponding watertight mesh with a single manifold (B). Perisynaptic processes, perivascular processes, and endfeet are colored in red, blue and green respectively. The wireframe closeup highlights the topology of the mesh around the astrocytic soma.

### 4.4 Application: reaction-diffusion simulation

While resulting meshes can still be used for other analytics applications that necessitate watertightness such as diffusion MRI simulations^42^, the principal objective of our work is to automate the reaction-diffusion simulation pipelines as demonstrated earlier in Figure 2. From an abstract point-and-diameter description of the neuron, a faithful and geometrically realistic surface mesh model is created with which we can synthesize a tetrahedral counterpart using TetGen to ultimately run an accurate STEPS simulation, for example Ca^+2^ signaling. Figure 12 shows a visualization of randomly generated simulation reports with a geometrically realistic pyramidal neuron model. The mechanisms of the simulation as well as the interpretation of the simulation results are beyond the scope of our work.

**Figure 12.**
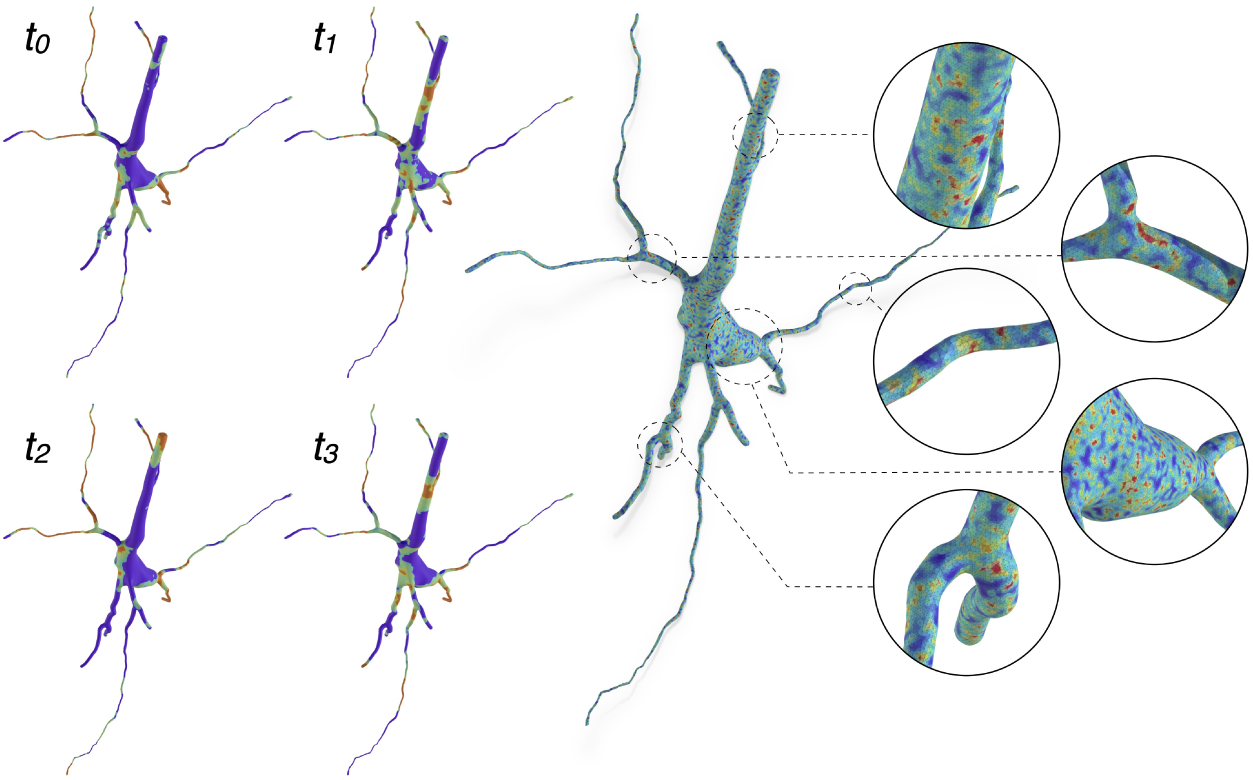
A tetrahedral mesh of a pyramidal neuron visualizing random simulation reports at multiple time steps, mimicking the variations of the Ca^+2^ signals across the cellular membrane. The watertight mesh created with our implementation is used to synthesize the tetrahedral counterpart relying on TetGen.

### 4.5 Comparative analysis with existing methods

There are several methods that can synthesize neuronal mesh representations from morphological trees^21,22,25,30^, but there are only two implementations capable of generating watertight manifolds: the CTNG technique^32^ and AnaMorph^33^. The validity of the resulting meshes from the CTNG implementation could not be verified because the code is deprecated and cannot be executed on recent hardware. However, we were able to make some comparative analysis demonstrating the resulting manifolds synthesized with our implementation and those generated with AnaMorph. This analysis is illustrated in Figure 13. Our implementation outperforms AnaMorph in two principal aspects: (i) AnaMorph can only approximate somatic profiles using primitive sphere, while our implementation generates a faithful and realistic 3D somatic profile using physics simulation, and (ii) AnaMorph has no support to load or integrate geometrically realistic spines along the dendritic branches of the neuronal membrane.

**Figure 13.**
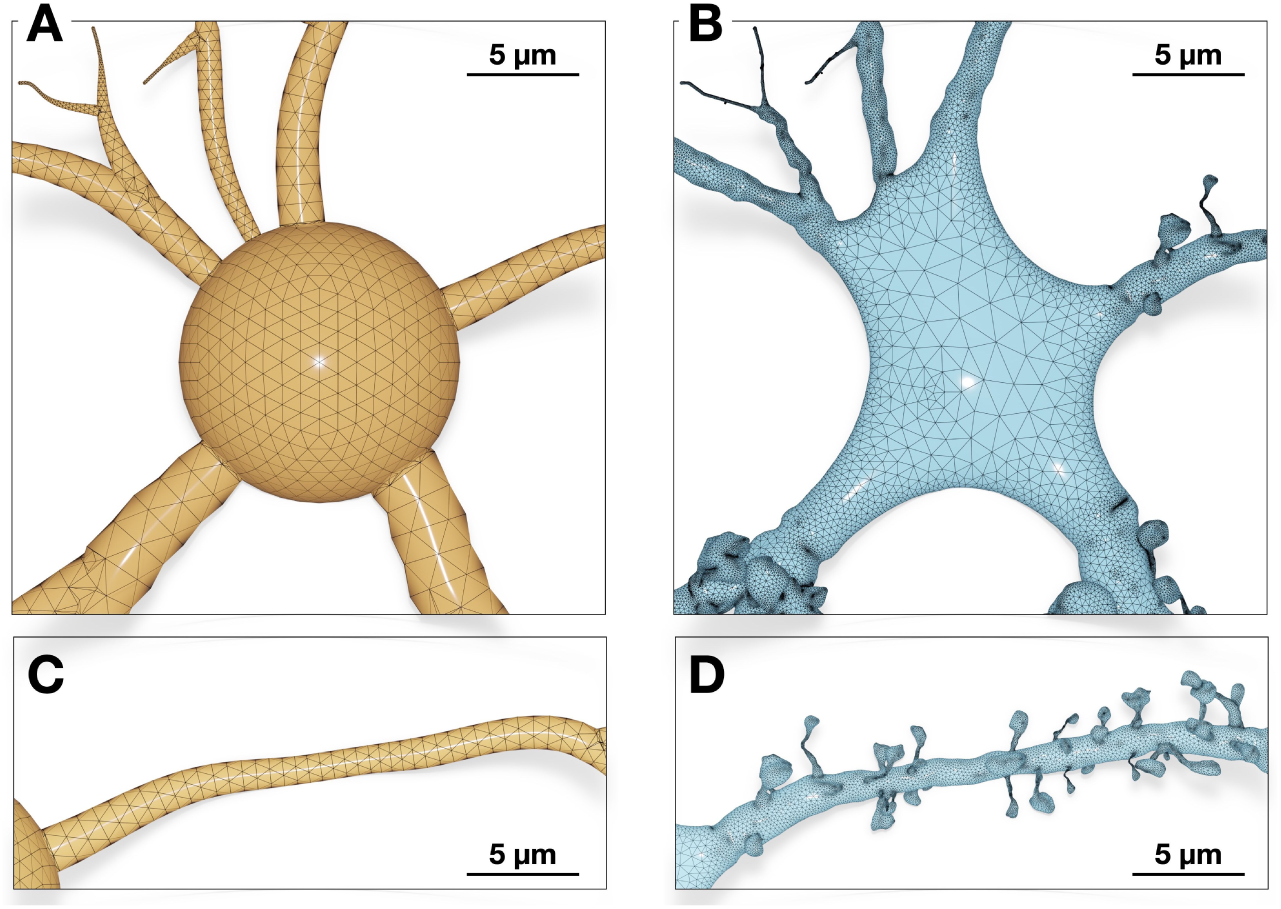
A side-by-side comparison between a neuronal mesh reconstructed with AnaMorph^33^ (in orange) and our method (in light blue). While AnaMorph approximates the soma with a symbolic sphere (A), our method is capable of reconstructing a faithful 3D profile of the soma based on mass-spring modeling and soft body dynamics (B). The meshing algorithm of AnaMorph cannot integrate any realistic spines along the dendritic branches of the neuron (C). Our implementation can seamlessly integrate detailed and highly realistic spine geometries that emanate smoothly from the neuronal membrane (D).

## 5 Conclusion

With the recent advances of computational modeling in neuroscience, detailed and geometrically realistic spatial models of neurons are becoming essential to better understand the impact of cellular morphology and subcellular structures on the underlying functions. While there is major ongoing research and increasing demand to build more efficient and scalable molecular simulators, the models with which we can perform accurate simulations are lacking. Biologically speaking, neurons have interweaving arborizations characterized by thin branches, complex branching geometries and large spatial extents. For these reasons, creating high quality and simulation-ready models that can accurately reflect their plasma membranes is challenging. In this use case, we present an intuitive, efficient and robust method capable of creating watertight mesh models of spiny neurons from their corresponding morphological descriptions. Our method uses the modeling toolsets in Blender to initially create a set of accurate but non-watertight and overlapping proxies and then uses the Voxel remesher to create a continuous watertight manifold that can accurately represent the surface membrane of the neuron. The performance and scalability of our implementation is assessed with a mixed dataset of neurons representing 60 various types of cortical morphologies. To make it publicly available and easily accessible, the implementation is integrated into the domain-specific NeuroMorphoVis framework. This integration expands the usability of the framework as a meshing tool, making it possible to generate lightweight meshes for visual analytics purposes on the one hand and to generate geometrically realistic watertight manifolds on the other hand. Our method can be seamlessly applied to other brain cells such as astrocytes and microglia, and even cerebral vasculature networks which are represented by cyclic graphs.

## Data sources

Neuronal morphologies shown in **Figure 10** and **Supplementary Figures S7 - S125** are available from the reconstructions made by Henry Markram^6^. Similar neuronal morphologies are publicly available from Neuro-Morpho.Org^15^. Astrocytic morphologies (**Figure 11**) are provided by Eleftherios Zisis^34^. Supplementary data including the resulting meshes of the 60 morphologies described in Figure 10 and their analysis factsheets are available on Zenodo (https://doi.org/10.5281/zenodo.10558475).

## Software availability

The voxelization-based remeshing algorithm is implemented in Blender^43^ based on its Python API. The technique is integrated within the Meshing Toolbox of the NeuroMorphoVis^28^ add-on. The code is released to public as an open-source software (OSS) in accordance with the regulations of the Blue Brain Project, École polytechnique fédérale de Lausanne (EPFL) for open sourcing under the GNU GPL3 license.

## Acknowledgments

We thank Grigori Chevtchenko for the impactful discussions on watertight meshing and Pawel Podhajski on technical assistance to deploy the software on Blue Brain 5. We also thank Karin Holm for her valuable comments on the manuscript.

## Authors’ contributions

M.A. conceived the study. M.A., A.Fo. and JJ.G.C. co-led the study. M.A. designed and implemented the algorithm in NeuroMorphoVis, reconstructed and analyzed the resulting models and wrote the manuscript with input and critique from all authors. A.Fo. implemented the morphology and mesh analysis code. JJ.G.C. assisted in the implementation of the soma generation algorithm and contributed to the manuscript. N.R.G implemented the surface smoothing filters and contributed to the manuscript. E.B contributed to the design of the figures. A.Fl. improved the performance of the optimization code and assisted in the integration of the code via pybind. JS.C. contributed to the simulation discussions and the manuscript. D.K. contributed to the discussions on the simulation requirements and the watertight meshing performance. J.P., JD.C. and G.K managed the project. All authors reviewed and approved the manuscript.

## Authors’ Biography

**Marwan Abdellah** is a senior visualization research engineer in the Visualization section of the Computing division at the Blue Brain Project of the École polytechnique fédérale de Lausanne (EPFL). He received his Ph.D. in neuroscience from EPFL in 2017.

**Alessandro Foni** is a system specialist at the Blue Brain Project of the École polytechnique fédérale de Lausanne (EPFL). He received his Ph.D. Information Systems from the faculty of Economics and Social Sciences of the University of Geneva in 2013.

**Juan José García Cantero** is a software engineer and visualization researcher at the Blue Brain Project of the École polytechnique fédérale de Lausanne (EPFL). He holds a Ph.D. in Computer Graphics from the Rey Juan Carlos University in Spain.

**Nadir Román Guerrero** is a visualization software engineer at the Blue Brain Project of the École polytechnique fédérale de Lausanne (EPFL).

**Elvis Boci** is a system specialist and technical artist in the Visualization section of the Computing division at the Blue Brain Project of the École polytechnique fédérale de Lausanne (EPFL).

**Adrien Fleury** is a software engineer in the Visualization section of the Computing division at the Blue Brain Project of the École polytechnique fédérale de Lausanne (EPFL).

**Jay S. Coggan** is a senior scientist in the Molecular Systems group in the Simulation Neuroscience division at the Blue Brain Project of the École polytechnique fédérale de Lausanne (EPFL).

**Daniel Keller** is the group leader of the Molecular Systems group at the Blue Brain Project of the École polytechnique fédérale de Lausanne (EPFL). His PhD in Neuroscience was achieved with the Computational Neurobiology Laboratory at the Salk Institute, UCSD.

**Judit Planas** is the manager of the Visualization section at Blue Brain Project of the École polytechnique fédérale de Lausanne (EPFL). She obtained her PhD in High Performance Computing from the Barcelona Supercomputing Center.

**Jean-Denis Courcol** is the section manager of the Neuroscientific Software Engineering team and deputy head of the Computing division at the Blue Brain Project of the École polytechnique fédérale de Lausanne (EPFL).

**Georges Khazen** is the associate director of Scientific Coordination section at the Blue Brain Project of the École polytechnique fédérale de Lausanne (EPFL). He received his PhD in Neuroscience from EPFL in 2011.

## Additional information

**Supplementary information** is available in the attached **Supplementary Document**.

**Correspondence and requests for materials** should be addressed to M.A.

## References

1. Keller, D. X., Franks, K. M., Bartol Jr, T. M. & Sejnowski, T. J. Calmodulin activation by calcium transients in the postsynaptic density of dendritic spines. PloS one 3, e2045. doi: 10.1371/journal.pone.0002045 (2008).

2. D’Angelo, E. & Jirsa, V. The quest for multiscale brain modeling. Trends in neurosciences. doi: 10.1016/j.tins.2022.06.007 (2022).

3. Di Ventura, B., Lemerle, C., Michalodimitrakis, K. & Serrano, L. From in vivo to in silico biology and back. Nature 443, 527–533. doi: 10.1038/nature05127 (2006).

4. Coggan, J. S. et al. A process for digitizing and simulating biologically realistic oligocellular networks demonstrated for the neuro-glio-vascular ensemble. Frontiers in neuroscience 12. doi: 10.3389/fnins.2018.00664 (2018).

5. Hines, M. L. & Carnevale, N. T. The NEURON simulation environment. Neural computation 9, 1179–1209. doi: 10.1162/neco.1997.9.6.1179 (1997).

6. Markram, H., Muller, E., Ramaswamy, S., Reimann, M. W., Abdellah, M., et al. Reconstruction and simulation of neocortical microcircuitry. Cell 163, 456–492. doi: 10.1016/j.cell.2015.09.029 (2015).

7. Kumbhar, P. et al. CoreNEURON: an optimized compute engine for the NEURON simulator. Frontiers in neuroinformatics 13, 63. doi: 10.3389/fninf.2019.00063 (2019).

8. Hepburn, I., Chen, W., Wils, S. & De Schutter, E. STEPS: efficient simulation of stochastic reaction–diffusion models in realistic morphologies. BMC systems biology 6, 1–19. doi: 10.1186/1752-0509-6-36 (2012).

9. Coggan, J. S. et al. Evidence for ectopic neurotransmission at a neuronal synapse. Science 309, 446–451. doi: 10.1126/science.1108239 (2005).

10. Grein, S., Stepniewski, M., Reiter, S., Knodel, M. M. & Queisser, G. 1D-3D hybrid modeling—from multi-compartment models to full resolution models in space and time. Frontiers in neuroinformatics 8, 68. doi: 10.3389/fninf.2014.00068 (2014).

11. Bartol, T. M. et al. Computational reconstitution of spine calcium transients from individual proteins. Frontiers in synaptic neuroscience 7, 17. doi: 10.3389/fnsyn.2015.00017 (2015).

12. Chen, W., et al. STEPS 4.0: Fast and memory-efficient molecular simulations of neurons at the nanoscale. *bioRxiv*, 2022–03. doi: 10.3389/fninf.2022.883742 (2022).

13. Hang, S. TetGen, a Delaunay-based quality tetrahedral mesh generator. ACM Trans. Math. Softw 41, 11. doi: 10.1145/2629697 (2015).

14. Hu, Y. et al. Tetrahedral meshing in the wild. ACM Trans. Graph. 37, 60–1. doi: 10.1145/3197517.3201353 (2018).

15. Ascoli, G. A., Donohue, D. E. & Halavi, M. NeuroMorpho.Org: a central resource for neuronal morphologies. Journal of Neuroscience 27, 9247–9251. doi: 10.1523/JNEUROSCI.2055-07.2007 (2007).

16. Perkins, G. et al. Electron tomography of large, multicomponent biological structures. Journal of structural biology 120, 219–227. doi: 10.1006/jsbi.1997.3920 (1997).

17. Donohue, D. E. & Ascoli, G. A. Automated reconstruction of neuronal morphology: an overview. Brain research reviews 67, 94–102. doi: 10.1016/j.brainresrev.2010.11.003 (2011).

18. Gleeson, P., Steuber, V. & Silver, R. A. neuroConstruct: a tool for modeling networks of neurons in 3D space. Neuron 54, 219– 235. doi: 10.1016/j.neuron.2007.03.025 (2007).

19. Kanari, L. et al. Objective Morphological Classification of Neocortical Pyramidal Cells. Cerebral Cortex 29, 1719–1735. doi: 10.1093/cercor/bhy339 (2019).

20. Eilemann, S., et al. Parallel rendering on hybrid multi-gpu clusters in Eurographics Symposium on Parallel Graphics and Visualization (2012), 109–117. doi: 10.2312/EGPGV/EGPGV12/109-117.

21. Lasserre, S. et al. A neuron membrane mesh representation for visualization of electrophysiological simulations. IEEE Transactions on Visualization and Computer Graphics 18, 214–227. doi: 10.1109/TVCG.2011.55 (2012).

22. Abdellah, M., Favreau, C., Hernando, J., Lapere, S. & Schürmann, F. Generating high fidelity surface meshes of neocortical neurons using skin modifiers in Computer Graphics and Visual Computing (CGVC) (eds Vidal, F. P., Tam, G. K. L. & Roberts, J. C.) (The Eurographics Association, 2019). isbn: 978-3-03868-096-3. doi: 10.2312/cgvc.20191257.

23. Hernando, J. B., Biddiscombe, J., Bohara, B., Eilemann, S. & Schürmann, F. Practical Parallel Rendering of Detailed Neuron Simulations in Eurographics Symposium on Parallel Graphics and Visualization (eds Marton, F. & Moreland, K.) (The Eurographics Association, 2013). doi: 10.2312/EGPGV/EGPGV13/049-056.

24. Eilemann, S., Makhinya, M. & Pajarola, R. Equalizer: A scalable parallel rendering framework. IEEE transactions on visualization and computer graphics 15, 436–452. doi: 10.1109/TVCG.2008.104 (2009).

25. Brito, J. P. et al. Neuronize: a tool for building realistic neuronal cell morphologies. Frontiers in neuroanatomy 7. doi: 10.3389/fnana.2013.00015 (2013).

26. Garcia-Cantero, J. J., Brito, J. P., Mata, S., Bayona, S. & Pastor, L. NeurotessMesh: A tool for the Generation and Visualization of Neuron Meshes and Adaptive on-the-Fly Refinement. Frontiers in neuroinformatics 11, 38. doi: 10.3389/fninf.2017.00038 (2017).

27. Abdellah, M. et al. Reconstruction and visualization of large-scale volumetric models of neocortical circuits for physically-plausible in silico optical studies. BMC bioinformatics 18, 402. doi: 10.1186/s12859-017-1788-4 (2017).

28. Abdellah, M. et al. NeuroMorphoVis: a collaborative framework for analysis and visualization of neuronal morphology skeletons reconstructed from microscopy stacks. Bioinformatics 34, i574–i582. doi: 10.1093/bioinformatics/bty231. https://github.com/BlueBrain/NeuroMorphoVis (2018).

29. Glaser, J. R. & Glaser, E. M. Neuron imaging with Neurolucida—a PC-based system for image combining microscopy. Computerized Medical Imaging and Graphics 14, 307–317. doi: 10.1016/0895-6111(90)90105-K (1990).

30. Abdellah, M., et al. Meshing of spiny neuronal morphologies using union operators in Computer Graphics and Visual Computing (CGVC) (eds Vangorp, P. & Turner, M. J.) (The Eurographics Association, 2022). isbn: 978-3-03868-188-5. doi: 10.2312/cgvc.20221168.

31. Velasco, I. et al. Neuronize v2: bridging the gap between existing proprietary tools to optimize neuroscientific workflows. Frontiers in Neuroanatomy 14, 585793. doi: 10.3389/fnana.2020.585793 (2020).

32. McDougal, R. A., Hines, M. L. & Lytton, W. W. Water-tight membranes from neuronal morphology files. Journal of neuroscience methods 220, 167–178. doi: 10.1016/j.jneumeth.2013.09.011 (2013).

33. Mörschel, K., Breit, M. & Queisser, G. Generating neuron geometries for detailed three-dimensional simulations using anamorph. Neuroinformatics 15, 247–269. doi: 10.1007/s12021-017-9329-x (2017).

34. Zisis, E. et al. Digital reconstruction of the neuro-glia-vascular architecture. Cerebral Cortex 31, 5686–5703. doi: 10.1093/cercor/bhab254 (2021).

35. Abdellah, M. et al. Metaball skinning of synthetic astroglial morphologies into realistic mesh models for in silico simulations and visual analytics. Bioinformatics 37, i426–i433. doi: 10.1093/bioinformatics/btab280 (2021).

36. Gipson, C. D. & Olive, M. F. Structural and functional plasticity of dendritic spines–root or result of behavior? *Genes*, Brain and Behavior 16, 101–117. doi: 10.1111/gbb.12324 (2017).

37. Son, J., Song, S., Lee, S., Chang, S. & Kim, M. Morphological change tracking of dendritic spines based on structural features. Journal of microscopy 241, 261–272. doi: 10.1111/j.1365-2818.2010.03427.x (2011).

38. Chen, W. & De Schutter, E. Time to bring single neuron modeling into 3D. Neuroinformatics 15, 1–3. doi: 10.1007/s12021-016-9321-x (2017).

39. Attene, M. A lightweight approach to repairing digitized polygon meshes. The visual computer 26, 1393–1406. doi: 10.1007/s00371-010-0416-3 (2010).

40. Ramaswamy, S. et al. The neocortical microcircuit collaboration portal: a resource for rat somatosensory cortex. Frontiers in neural circuits 9. doi: 10.3389/fncir.2015.00044 (2015).

41. Pébay, P. P., et al. New applications of the verdict library for standardized mesh verification pre, post, and end-to-end processing in Proceedings of the 16th international meshing roundtable (2008), 535–552. doi: 10.1007/978-3-540-75103-8_30.

42. Fang, C., Nguyen, V.-D., Wassermann, D. & Li, J.-R. Diffusion MRI simulation of realistic neurons with SpinDoctor and the Neuron Module. NeuroImage 222, 117198. doi: 10.1016/j.neuroimage.2020.117198 (2020).

43. Blender. An open source 3D modelling and rendering package. The Blender Foundation (Blender Institute, Amsterdam, 2024).

