## Supplementary Document for "Synthesis of geometrically realistic and watertight neuronal ultrastructure manifolds for *in silico* modeling"

#### Supplementary Material

Marwan Abdellah,* Alessandro Foni, Juan José García Cantero, Nadir Román Guerrero, Elvis Boci, Adrien Fleury, Jay S. Coggan, Daniel Keller, Judit Planas, Jean-Denis Courcol, and Georges Khazen

Blue Brain Project
École Polytecnique Fédérale de Lausanne (EPFL)
Geneva, Switzerland

78 Pages, 127 Figures

July 2024

### Contents

**1 Watertight manifolds** 4

**2 Digitally reconstructed cortical circuits and neuronal morphological types** 5

**3 Reconstruction of watertight manifolds of geometrically realistic neurons** 8

**4 Surface mesh optimization** 10

**5 Integration of dendritic spine models with realistic geometries** 15

**6 Quantitative and qualitative measures** 16

**7 Comparative performance analysis** 76

**8 Software** 77

**9 Supplementary data** 77

### List of Tables

S1 Summary of the selected neurons from a recent digitally reconstructed cortical circuit[2,3] and their morphological types and cell identifiers (GIDs) in the circuit. The analysis of the resulting meshes of each neuronal morphology is shown in each corresponding figure in Section 6. . . 5

### 1 Watertight manifolds

A watertight surface mesh is a manifold that consists of one closed surface, i.e. it does not contain any gaps or holes and have a clearly defined boundary and inside. By definition, a surface mesh is watertight if the following conditions are met: (i) the mesh has no self-intersecting faces, (ii) the mesh is two-manifold, i.e. it does not contain any non-manifold edges or non-manifold vertices, and (iii) the mesh has no boundary edges. A self-intersection is an intersection of two facets belonging to the same mesh. A non-manifold edge is an edge that has more or less than two incident faces. If the edge is connected to only one facet, it is a non-manifold boundary edge.

To understand what a non-manifold vertex is, we define the *star of a vertex* to be the union of all its incident faces. A non-manifold vertex is a vertex where the corresponding star is not any further connected after the removal of the vertex. A two-manifold mesh is a mesh that has zero non-manifold edges and non-manifold vertices. A watertight manifold is then a two-manifold mesh that has no self-intersecting faces and zero boundary edges[1]. Figure S1 illustrates the differences between manifold and non-manifold vertices and edges.

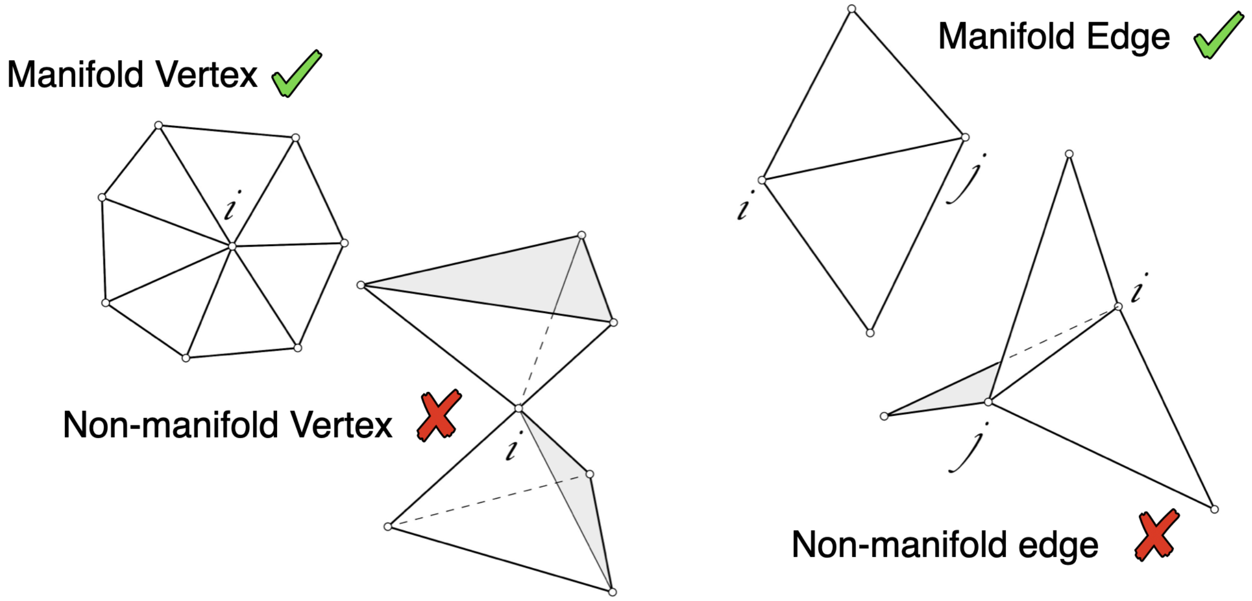

Figure S1: A comparative illustration showing the configurations of manifold and non-manifold vertices (left) and manifold and non-manifold edges (right). A watertight surface manifold must have zero non-manifold edges and zero non-manifold vertices. The vertex is labeled $i$, while the edge is labeled $ij$.

#### 2 Digitally reconstructed cortical circuits and neuronal morphological types

In 2015, a first large-scale model of the microcircuitry of somatosensory cortex of a two-weeks old rat is presented[2]. Using detailed anatomical and physiological models gathered from experimental data, a biologically plausible digital reconstruction of the cortical circuit is achieved. Recent circuits contains 60 different types of neuronal morphologies[3]. The robustness of the presented meshing pipeline (refer to Figure S2) is evaluated by applying the pipeline to a diverse set of neurons that are sampled from a recent digitally reconstructed circuit. We selected 60 cellular exemplars, where each cell represent a single morphological type. Table S1 lists those exemplars, their morphological types and cellular identifiers (or GIDs) in the circuit. Quantitative and qualitative analysis of the resulting meshes is discussed in Section 6 (Figs. S7 - S126).

Table S1: Summary of the selected neurons from a recent digitally reconstructed cortical circuit[2,3] and their morphological types and cell identifiers (GIDs) in the circuit. The analysis of the resulting meshes of each neuronal morphology is shown in each corresponding figure in Section 6.

| m-type | Morphological type[2] | GID | Mesh Analysis |
|---|---|---|---|
| L1_DAC | Layer 1 Descending Axon Cell | 6343 | Figs. S7 & S8 |
| L1_HAC | Layer 1 Horizontal Axon Cell | 662845 | Figs. S9 & S10 |
| L1_LAC | Layer 1 Large Axon Cell | 674955 | Figs. S11 & S12 |
| L1_NGC-DA | Layer 1 Neurogliaform Cell with Dense Axonal Arborization | 653769 | Figs. S13 & S14 |
| L1_NGC-SA | Layer 1 Neurogliaform Cell with Slender Axonal Arborization | 688678 | Figs. S15 & S16 |
| L1_SAC | Layer 1 Small Axon Cell | 681256 | Figs. S17 & S18 |
| L23_BP | Layer 2-3 Bipolar Cell | 3022157 | Figs. S19 & S20 |
| L23_BTC | Layer 2-3 Bitufted Cell | 2983868 | Figs. S21 & S22 |
| L23_CHC | Layer 2-3 Chandelier Cell | 3047737 | Figs. S23 &. S24 |
| L23_DBC | Layer 2-3 Double Bouquet Cell | 3417463 | Figs. S25 & S26 |
| L23_LBC | Layer 2-3 Large Basket Cell | 3019557 | Figs. S27 & S28 |
| L23_MC | Layer 2-3 Martinotti Cell | 508578 | Figs. S29 & S30 |
| L23_NBC | Layer 2-3 Nest Basket Cell | 3039549 | Figs. S31 & S32 |
| L23_NGC | Layer 2-3 Neurogliaform Cell | 2980862 | Figs. S33 & S34 |
| L23_SBC | Layer 2-3 Small Basket Cell | 502166 | Figs. S33 & S34 |
| L2_IPC | Layer 2 Inverted Pyramidal Cell | 2944367 | Figs. S37 & S38 |

| | | | |
|---|---|---|---|
| L2_TPC:A | Layer 2 Tufted Pyramidal Cell A | 2925968 | Figs. S39 & S40 |
| L2_TPC:B | Layer 2 Tufted Pyramidal Cell B | 3328718 | Figs. S41 & S42 |
| L3_TPC:A | Layer 3 Tufted Pyramidal Cell A | 452629 | Figs. S43 & S44 |
| L3_TPC:C | Layer 3 Tufted Pyramidal Cell C | 532420 | Figs. S45 & S46 |
| L4_BP | Layer 4 Bipolar Cell | 2206966 | Figs. S47 & S48 |
| L4_BTC | Layer 4 Bitufted Cell | 2859275 | Figs. S49 & S50 |
| L4_CHC | Layer 4 Chandelier Cell | 2208302 | Figs. S51 & S52 |
| L4_DBC | Layer 4 Double Bouquet Cell | 2380929 | Figs. S53 & S54 |
| L4_LBC | Layer 4 Large Basket Cell | 2875360 | Figs. S55 & S56 |
| L4_MC | Layer 4 Martinotti Cell | 2872311 | Figs. S57 & S58 |
| L4_NBC | Layer 4 Nest Basket Cell | 2797995 | Figs. S59 & S60 |
| L4_NGC | Layer 4 Neurogliaform Cell | 2378362 | Figs. S61 & S62 |
| L4_SBC | Layer 4 Small Basket Cell | 2381531 | Figs. S63 & S64 |
| L4_SSC | Layer 4 Spiny Stellate Cell | 2819361 | Figs. S65 & S66 |
| L4_TPC | Layer 4 Tufted Pyramidal Cell | 2776911 | Figs. S67 & S68 |
| L4_UPC | Layer 4 Untufted Pyramidal Cell | 2252026 | Figs. S69 & S70 |
| L5_BP | Layer 5 Bipolar Cell | 4234789 | Figs. S71 & S72 |
| L5_BTC | Layer 5 Bitufted Cell | 3597773 | Figs. S73 & S74 |
| L5_CHC | Layer 5 Chandelier Cell | 3422989 | Figs. S75 & S76 |
| L5_DBC | Layer 5 Double Bouquet Cell | 3608613 | Figs. S77 & S78 |
| L5_LBC | Layer 5 Large Basket Cell | 3489410 | Figs. S79 & S80 |
| L5_MC | Layer 5 Martinotti Cell | 4230916 | Figs. S81 & S82 |
| L5_NBC | Layer 5 Nest Basket Cell | 3569992 | Figs. S83 & S84 |
| L5_NGC | Layer 5 Neurogliaform Cell | 4212531 | Figs. S85 & S86 |
| L5_SBC | Layer 5 Small Basket Cell | 3512410 | Figs. S87 & S88 |
| L5_TPC:A | Layer 5 Thick-tufted Pyramidal Cell A | 4163878 | Figs. S89 & S90 |
| L5_TPC:B | Layer 5 Thick-tufted Pyramidal Cell B | 3794149 | Figs. S91 & S92 |

| | | | |
|---|---|---|---|
| L5_TPC:C | Layer 5 Thick-tufted Pyramidal Cell C | 3466005 | Figs. S93 & S94 |
| L5_UPC | Layer 5 Untufted Pyramidal Cell | 3547415 | Figs. S95 & S96 |
| L6_BP | Layer 6 Bipolar Cell | 944429 | Figs. S97 & S98 |
| L6_BPC | Layer 6 Pyramidal Cell with Bipolar Apical-like Dendrites | 744886 | Figs. S99 & S100 |
| L6_BTC | Layer 6 Bitufted Cell | 950455 | Figs. S101 & S102 |
| L6_CHC | Layer 6 Chandelier Cell | 1723993 | Figs. S103 & S104 |
| L6_DBC | Layer 6 Double Bouquet Cell | 1994509 | Figs. S105 & S106 |
| L6_HPC | Layer 6 Horizontal Pyramidal Cell | 1240273 | Figs. S107 & S108 |
| L6_IPC | Layer 6 Pyramidal Cell with Inverted Apical-like Dendrites | 1561862 | Figs. S109 & S110 |
| L6_LBC | Layer 6 Large Basket Cell | 1374612 | Figs. S111 & S112 |
| L6_MC | Layer 6 Martinotti Cell | 1122106 | Figs. S113 & S114 |
| L6_NBC | Layer 6 Nest Basket Cell | 2204257 | Figs. S115 & S116 |
| L6_NGC | Layer 6 Neurogliaform Cell | 962348 | Figs. S117. S118 |
| L6_SBC | Layer 6 Small Basket Cell | 1408681 | Figs. S119 & S120 |
| L6_TPC:A | Layer 6 Tufted Pyramidal Cell with Dendritic Tuft A | 1895896 | Figs. S121 & S122 |
| L6_TPC:C | Layer 6 Tufted Pyramidal Cell with Dendritic Tuft C | 2147655 | Figs. S123 & S124 |
| L6_UPC | Layer 6 Untufted Pyramidal Cell | 1063319 | Figs. S125 & S126 |

#### 3 Reconstruction of watertight manifolds of geometrically realistic neurons

Figure S2 shows a high level overview of our pipeline including watertight surface mesh generation, tetrahedralization and reaction-diffusion simulation. The principal focus of this work is the automated generation of optimized and watertight surface meshes that can be directly plugged into the simulation. Tetrahedralization[4] and reaction-diffusion simulations[5] are complementary steps that are beyond the scope of this work.

The input morphology is used to construct a list of proxy meshes, where each proxy corresponds to an individual object in the morphology (soma, branches, or spines). Proxies are grouped into a single mesh object, with which the Voxel remesher can be applied. The dimensions of the smallest structure in the proxy meshes are evaluated and the resolution (or Voxel Size) of the Voxel remesher is adjusted accordingly. This remesher uses an efficient variant of the marching cubes algorithm to construct a single manifold that represent the cellular membrane of the neuronal morphology. Typically, this manifold has highly tessellated surface with huge number of facets. Therefore, mesh optimization is applied to create a corresponding watertight manifold with a fewer number of facets that is convenient to run a simulation. The resulting surface mesh is adapted to create a corresponding tetrahedral volumetric mesh, for example using TetGen[4,6], and is plugged into a reaction-diffusion simulation in STEPs simulator[5,7,8].

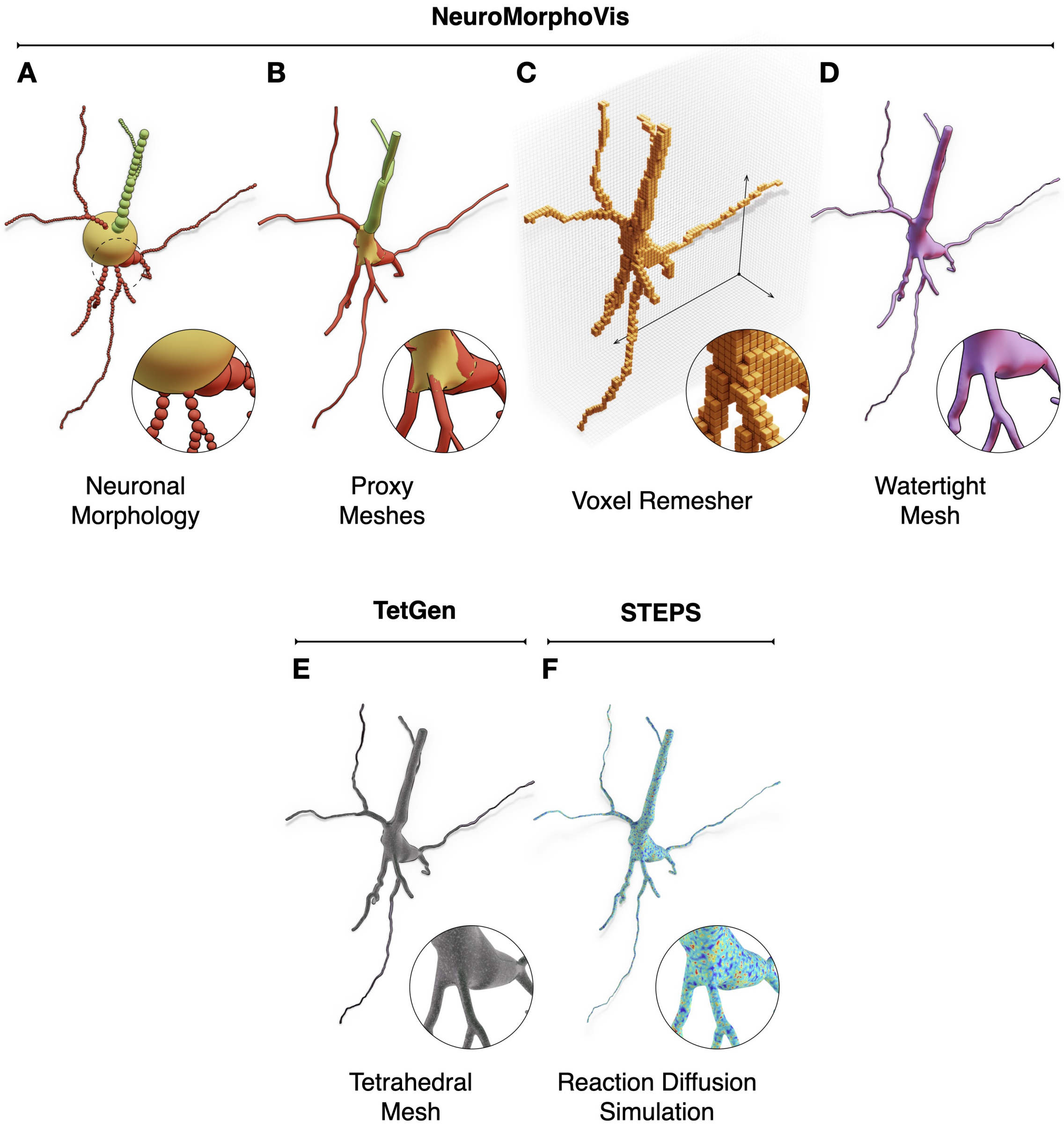

Figure S2: **Mesh generation & simulation pipeline**. The neuronal morphology (A) is initially used to create a set of corresponding proxy meshes of every individual component of the morphology, which are then combined into a single mesh object with overlapping geometries using a joint operation (B). The Voxel remesher is applied to this mesh object to create a volumetric representation of the membrane (C) with which all the overlapping structures are eliminated. This remesher creates a watertight manifold with a continuous and smooth surface (D), which is then optimized to synthesize a volumetric mesh (E), for example using TetGen, to perform a stochastic reaction-diffusion simulation in Steps (F). Spines are not shown.

### 4 Surface mesh optimization

#### 4.1 Re-tessellation via coarsening

The resulting mesh from the Voxel remesher in Blender[9] is reconstructed with an extension of the popular marching cubes algorithm. Based on the spatial extent of the mesh and the size of its smallest structure, the voxelization resolution is set, which often leads to reconstruct a mesh with gigantic number of facets that are uniformly distributed along the surface of the mesh. This mesh is mathematically guaranteed to be watertight, but it has two principal limitations when used in reaction-diffusion simulations. First, and due to its high tessellation, it is accompanied with high computational costs. Second, it has low geometric quality because the edges of its triangles are much different in length; i.e. the aspect ratio is less than one. Therefore, it will have poor numerical accuracy that is reflected on the results of the simulation.

To resolve these issues, we have adapted and extended the GAmer – or Geometry-preserving Adaptive MeshER – library[10]; and provided an optimized extension called OMesh (or OptimizationMesh). As the optimization procedure is applied per vertex, implementing the code in Python is obviously inefficient. Therefore, OMesh is developed in C++, but it has Python bindings, which makes it compatible with the Python API of Blender. Moreover, OMesh uses OpenMP to parallelize the embarrassingly parallel sections of the code. Further details about the code, its implementation aspects and installation are provided in Section 8.

Adaptive surface coarsening reduces the number of facets in local regions with low frequency features and preserves a decent amount of vertices to capture high frequency features as shown in Figure S3. The local regions across the mesh surface are quantified using a local structure tensor, where we can evaluate the number of vertices that can be safely eliminated without changing the structure. This evaluation is based on several factors including the local sparseness and curvature of the surface mesh at each vertex. Once a vertex is removed, the patch of the incident neighbors is re-triangulated to close the manifold.

#### 4.2 Self-intersections

While the surface coarsening process is significant to eliminate unnecessary vertices from the mesh and to reduce its computational complexity, re-triangulation of the holes caused by the deleted vertices introduces self-intersecting facets, leading to a non-watertight mesh as explained earlier in Section 1. These self-intersections can be reduced and possibly removed by applying triangular smoothing across the surface of the mesh in an iterative fashion. Figure S4 shows wireframe visualizations of the mesh shown in Figure S3 after every surface smoothing iteration for a total of 10 iterations. Surface smoothing tends to stretch the entire surface of the mesh trying to eliminate self-intersections, nonetheless, removing self-intersecting factes completely is not guaranteed. Typically, after 15 - 30 smoothing iterations, a relatively few self-intersecting facets – with respect to the mesh size – might still exist as shown in Figure S5, where 14 meshes (14 out of 60) still have self-intersecting faces even after 50 iterations of surface smoothing.

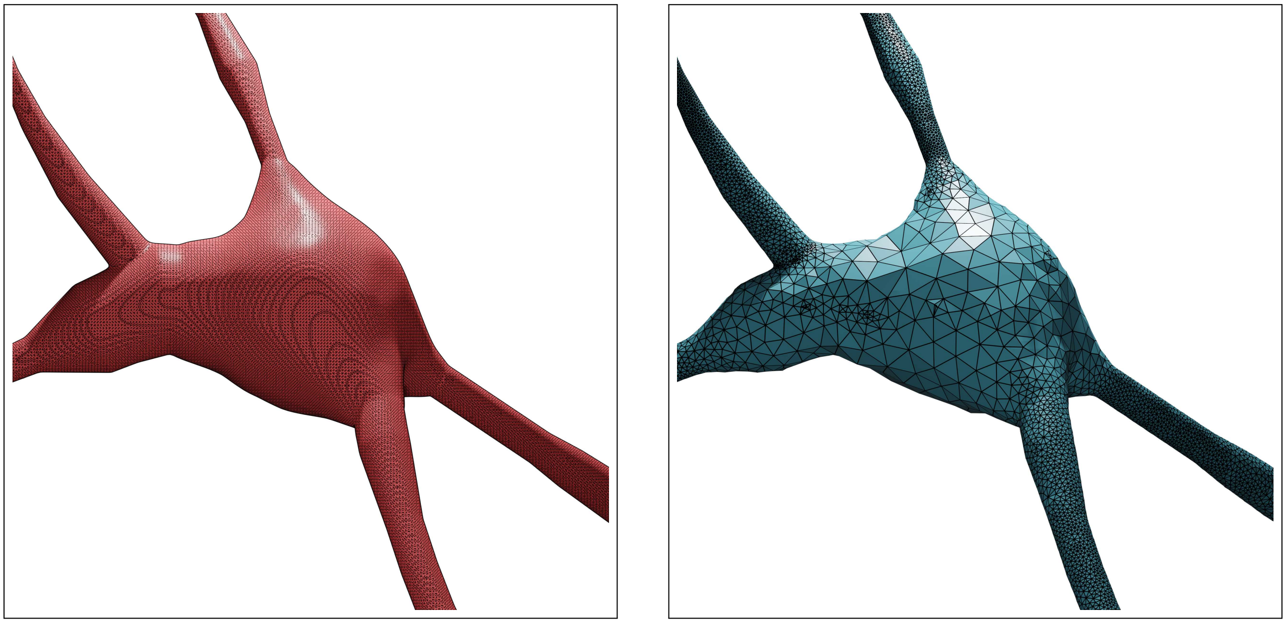

Figure S3: **Surface mesh coarsening**. The neuronal mesh generated from the Voxel remesher (left) is typically highly tessellated (∼100k triangles). This mesh is re-tessellated using coarsening to create an adaptively optimized clone (right) – with ∼68k triangles, where local regions with high frequency contain more faces than flat regions.

##### 4.3 Watertightness verification

To guarantee the robustness of our solution, we use the modeling tools in Blender, including the internal mesh editing API (called BMesh), to implement an iterative watertightness verification procedure to ensure that the optimized mesh (the blue mesh in Figure S3) is watertight. This procedure initially identifies if the mesh has non-manifold edges, non-manifold vertices or self-intersecting faces or not. If self-intersecting facets are detected, the corresponding vertices of those facets are identified and marked for deletion. The elimination of these vertices is accompanied with the generation of four artifacts: (i) non-manifold edges, (ii) possible non-manifold vertices, (iii) possible floating vertices and (iv) possible tiny floating partitions.

These artifacts are handled in the following order. Initially, if the mesh has any floating vertices, i.e. vertices that are not connected to any edges, we mark those floating vertices and eliminate them from the mesh all at once. Afterwards, we count the number of partitions in the mesh. In case the mesh has more than one partition, we select the largest partition, or the partition that has the largest number of vertices, and consider it the principal partition in the mesh. This partition is preserved, while the other secondary partitions (with significantly less number of vertices) are marked for removal. The vertices of the secondary mesh partitions are selected and eliminated from the mesh. At this stage, the principal partition has no self-intersections and zero non-manifold vertices, but it contains non-manifold edges that form multiple holes across the surface of the mesh. We then apply an efficient hold-filling strategy that takes a list of edges corresponding to the present non-manifold edges in the mesh to create a list of triangle facets leading to the repair of all the non-manifold edges. In the majority of the cases, filling the holes using this approach resolves the non-watertightness problem. But in a few cases, the newly created facets might intersect with other facets of the mesh. This case particularly happens with meshes containing sharp edges. If this scenario occurs, a new watertightness verification iteration is applied, where the self-intersecting facets are eliminated until a the mesh is confirmed to have no self-intersections and zero non-manifold edges and vertices.

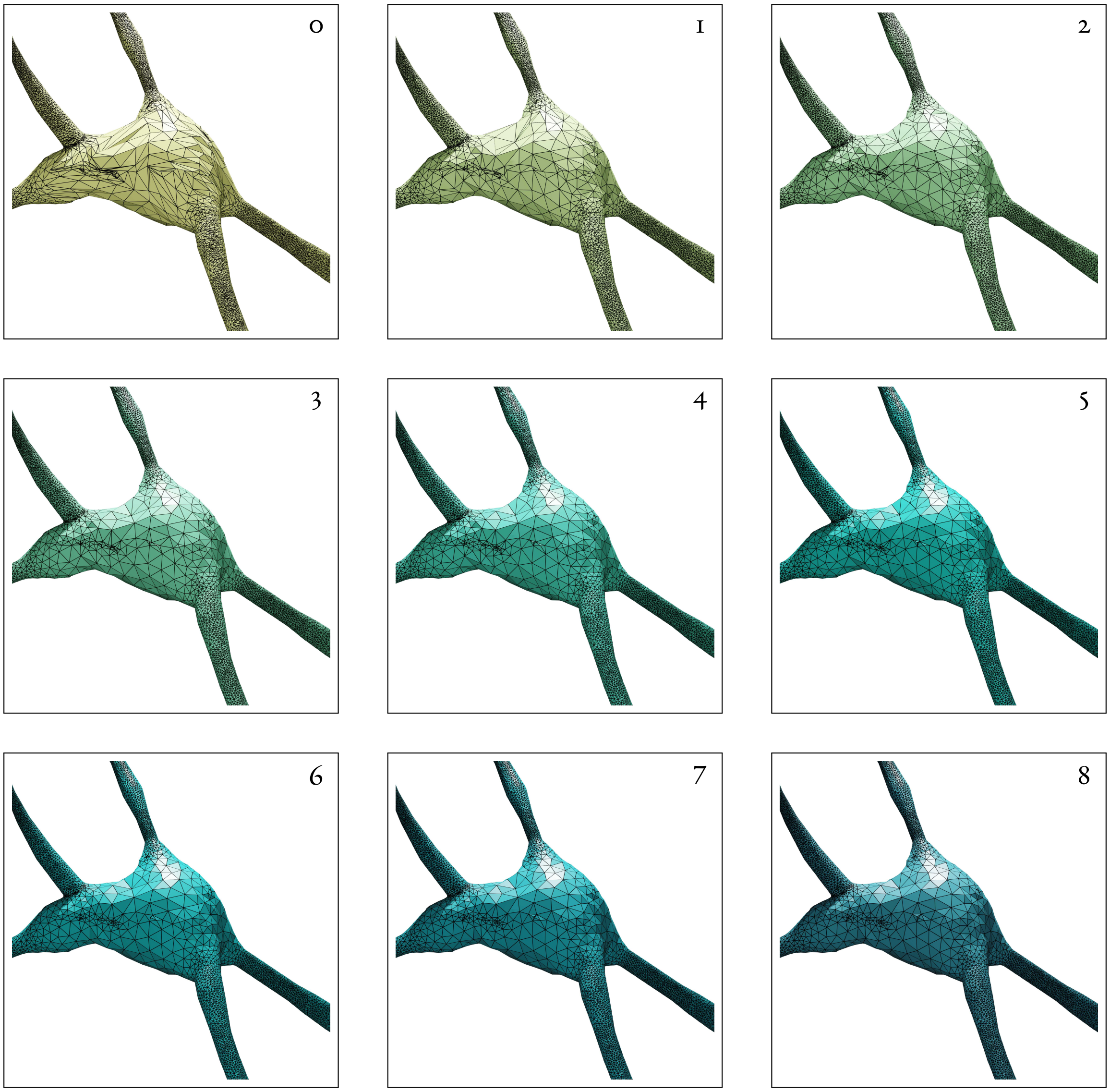

Figure S4: Iterative smoothing of a decimated surface mesh of a neuronal morphology. The decimation procedure – or mesh coarsening – introduces self-intersecting facets. In every smoothing iteration, the surface of the mesh is stretched and the number of self-intersecting facets is reduced. Nonetheless, and in certain complex geometric scenarios, it is not guaranteed to eventually remove all the self-intersections even after large number of iterations. The number of smoothing iterations is indicated on the top right of every rendering. Related to Fig. S5.

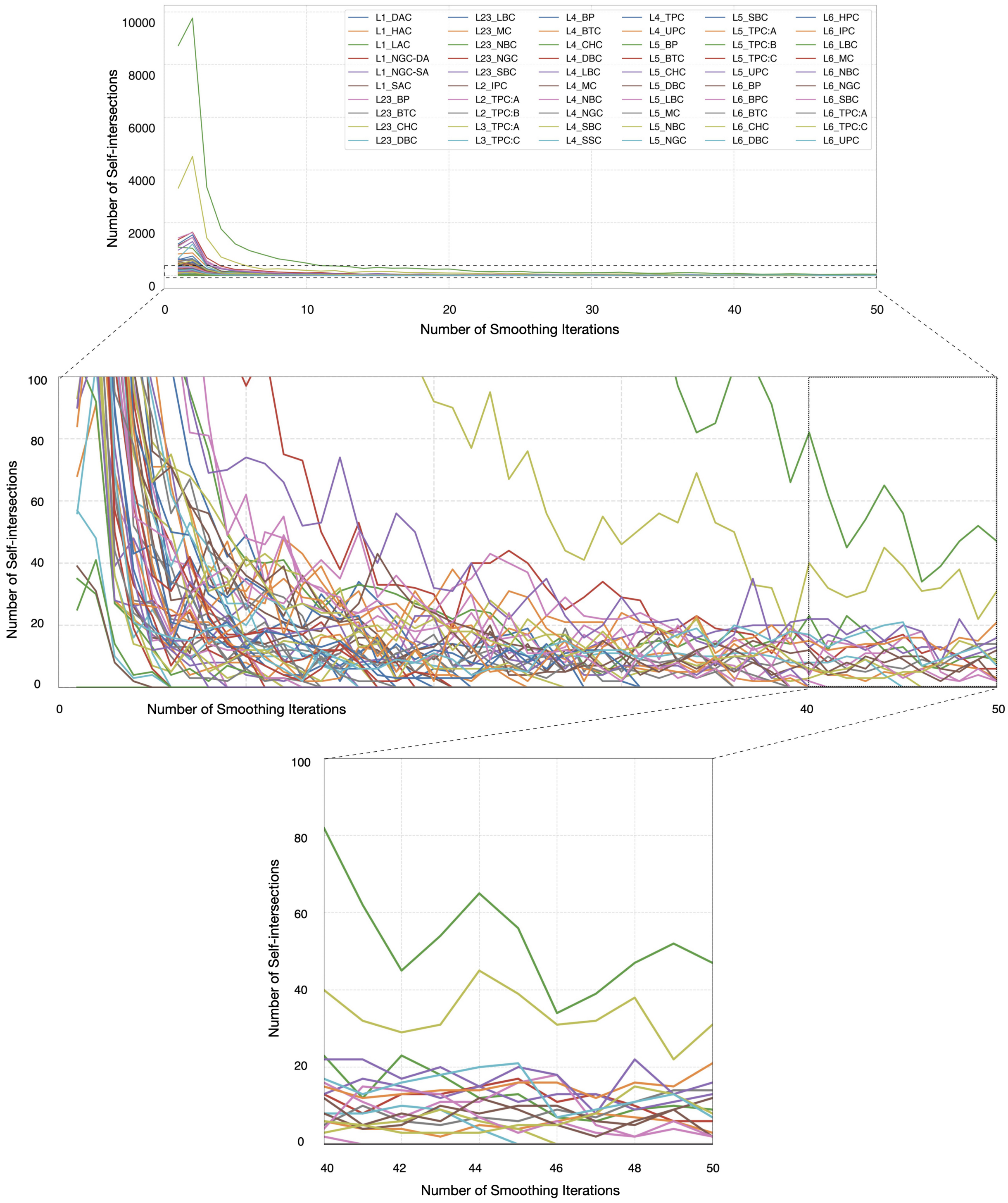

Figure S5: Number of self-intersecting facets with respect to number of smoothing iterations for the meshes created from their corresponding neuronal morphologies. Related to Fig. S4.

### 5 Integration of dendritic spine models with realistic geometries

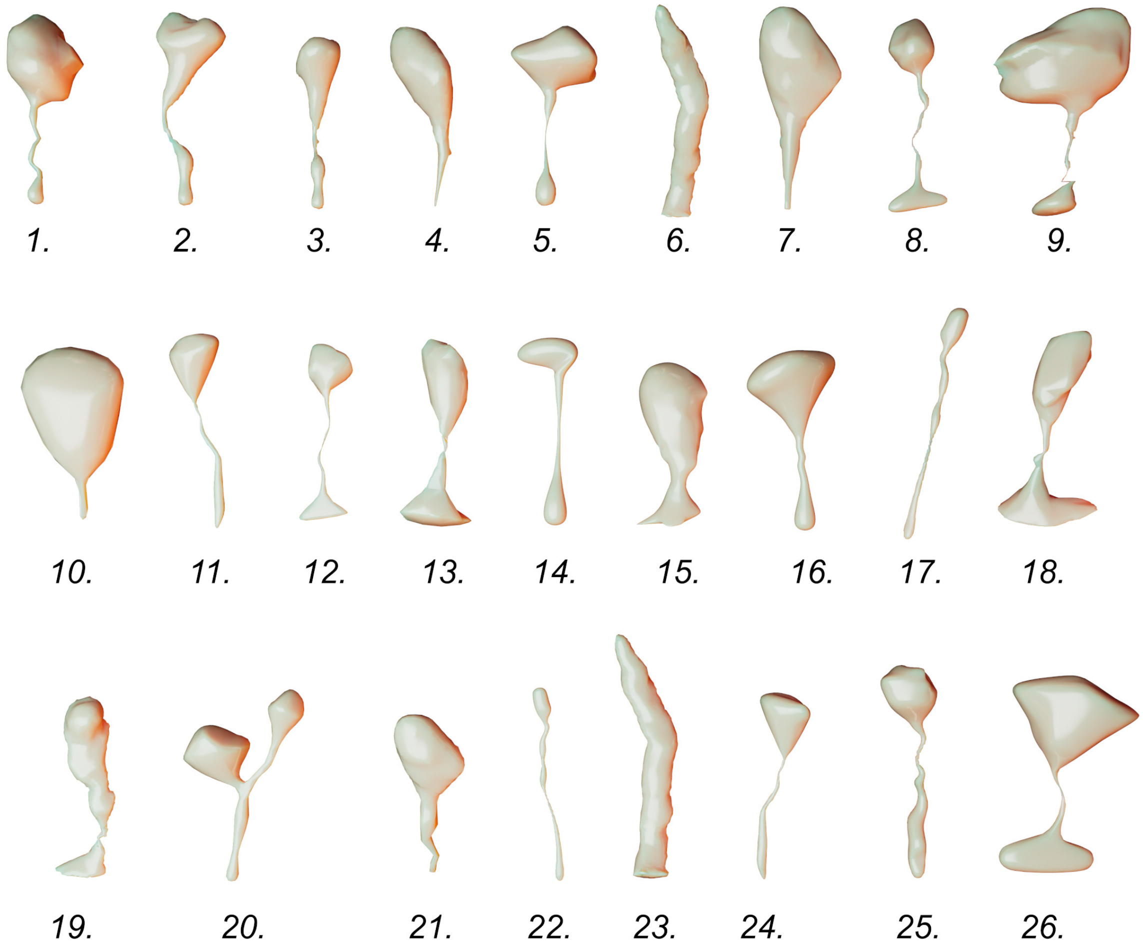

Figure S6: Spine mesh models with realistic geometries segmented from a cortical electron microscopy volume of a two-weeks old rat[11].

### 6 Quantitative and qualitative measures

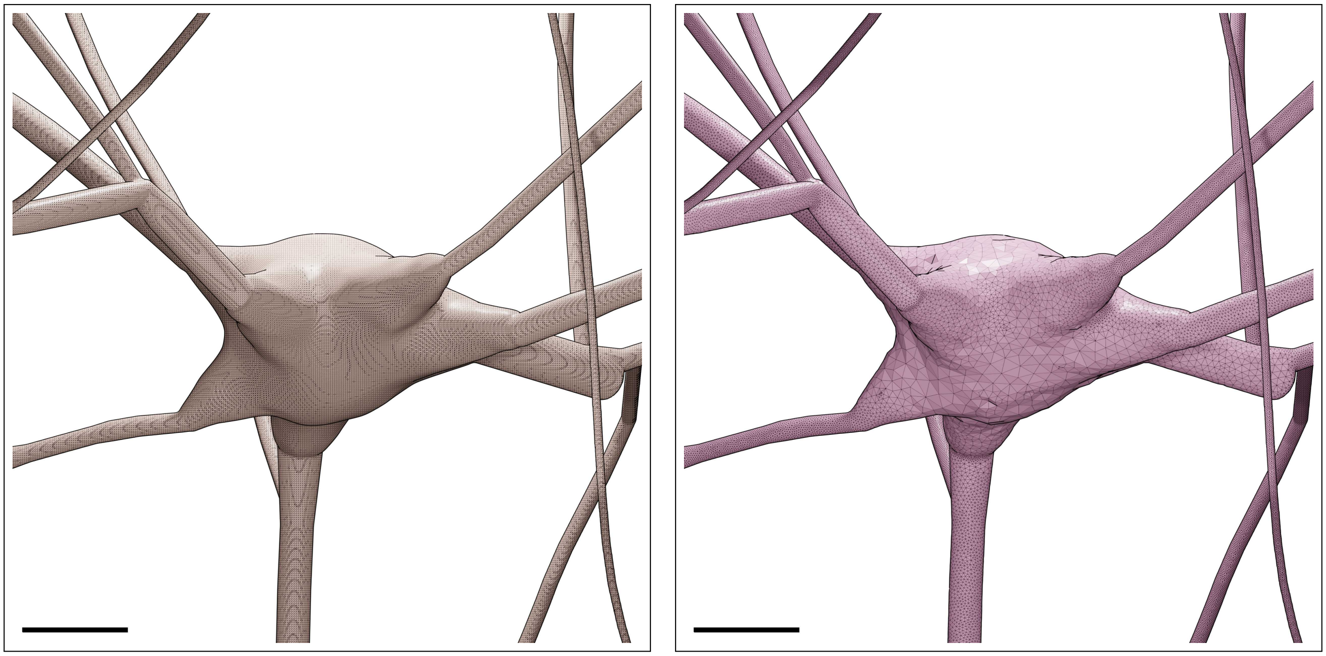

Figure S7: Wireframe visualizations of an L1_DAC neuron showing closeup comparisons between the highly tessellated surface mesh generated from the Voxel remesher (left) and the adaptively optimized surface mesh generated from the optimizer (right). Comparative quantitative and qualitative analyses of the meshes are demonstrated in Figure S8. Scale bars: 5 $\mu$m.

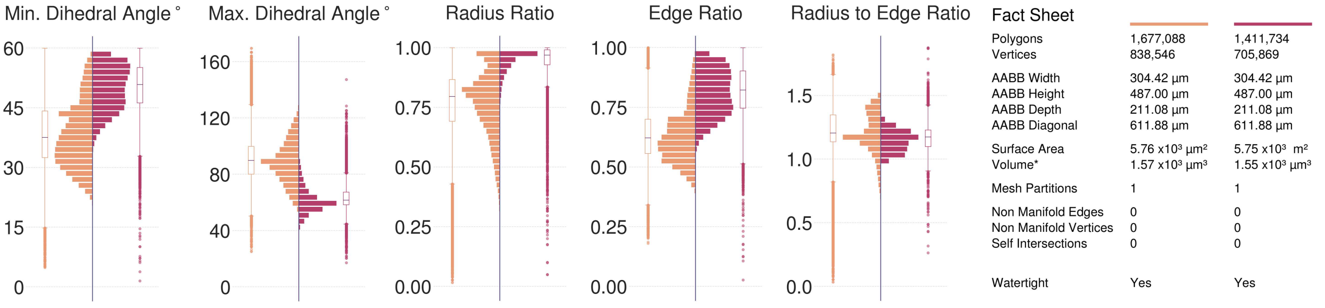

Figure S8: Comparative quantitative and qualitative analyses of the surface mesh models of the L1_DAC neuron visualized in Figure S7.

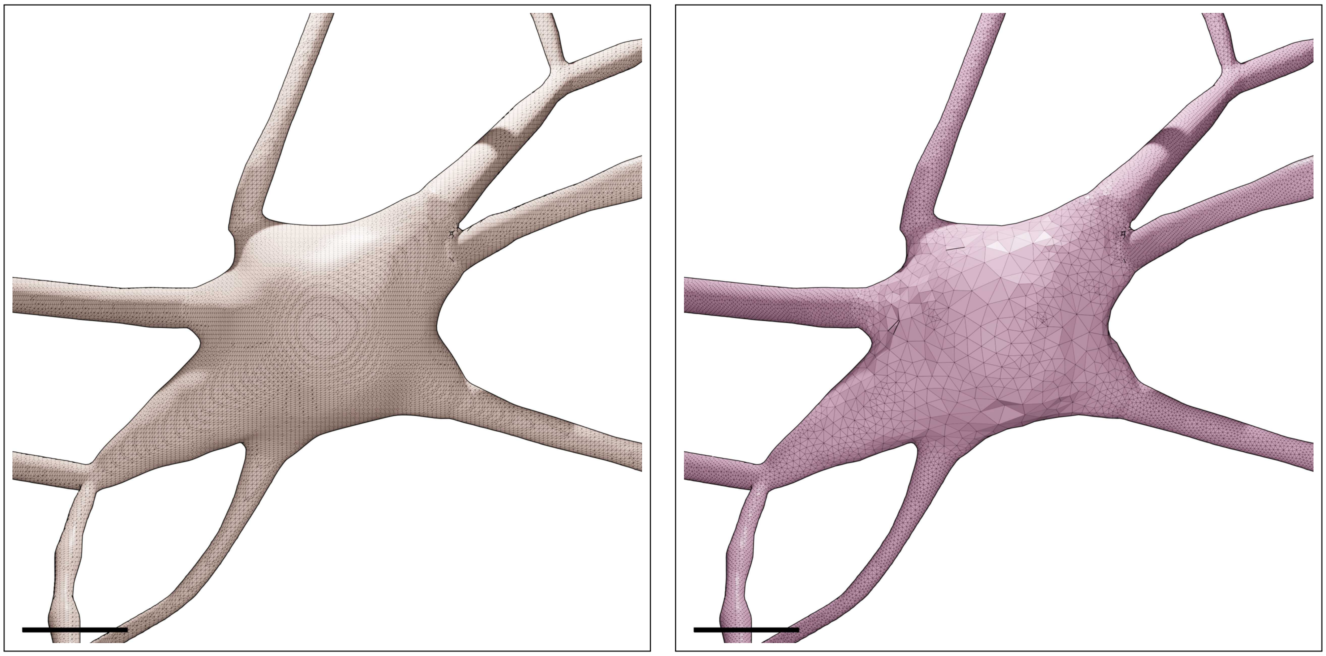

Figure S9: Wireframe visualizations of an L1_HAC neuron showing closeup comparisons between the highly tessellated surface mesh generated from the Voxel remesher (left) and the adaptively optimized surface mesh generated from the optimizer (right). Comparative quantitative and qualitative analyses of the meshes are demonstrated in Figure S10. Scale bars: 5 $\mu$m.

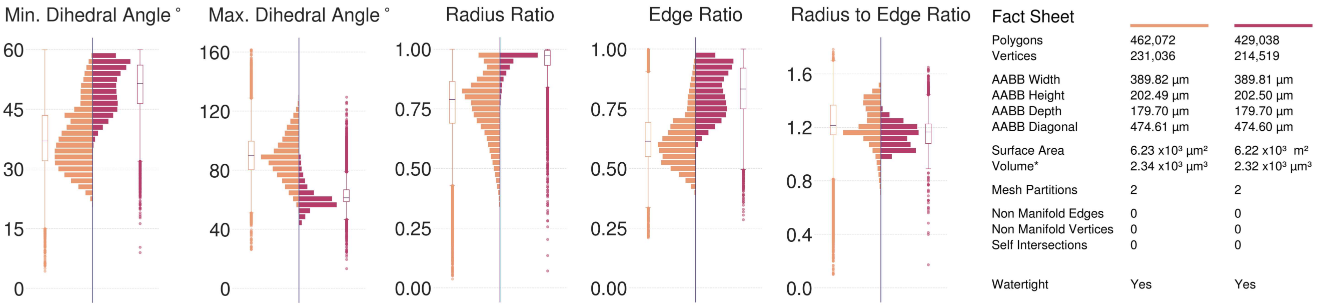

Figure S10: Comparative quantitative and qualitative analyses of the surface mesh models of the L1_HAC neuron visualized in Figure S9.

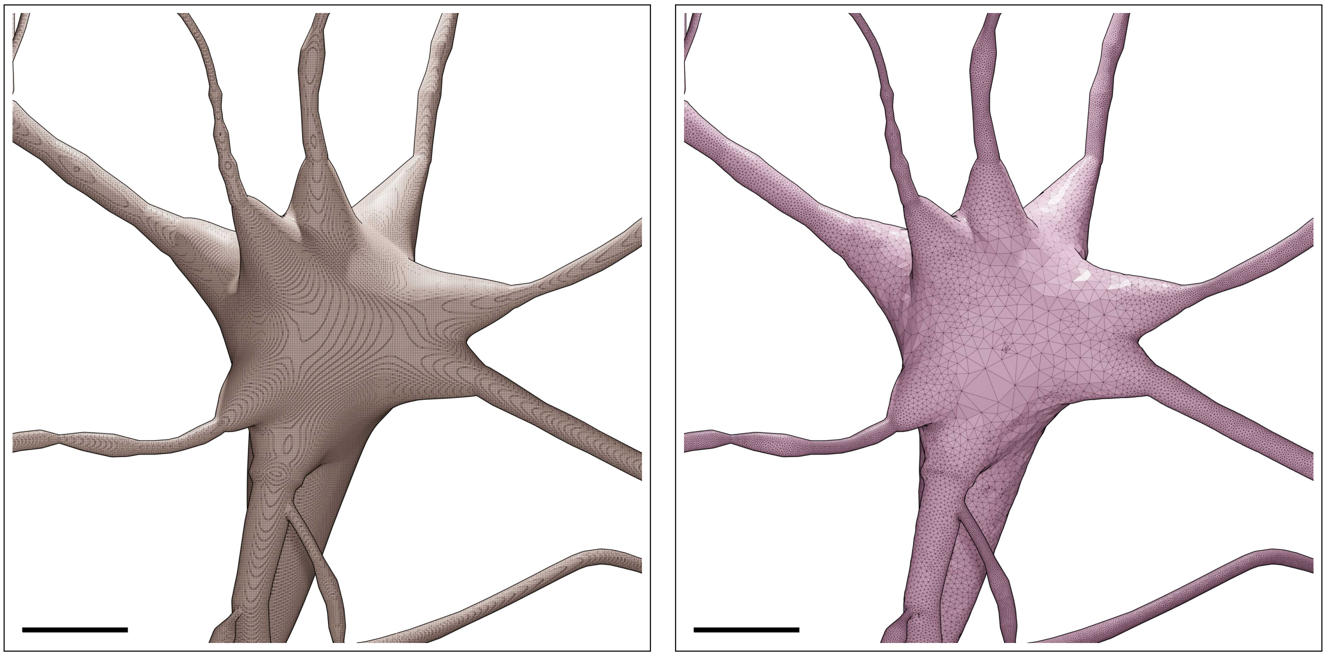

Figure S11: Wireframe visualizations of an L1_LAC neuron showing closeup comparisons between the highly tessellated surface mesh generated from the Voxel remesher (left) and the adaptively optimized surface mesh generated from the optimizer (right). Comparative quantitative and qualitative analyses of the meshes are demonstrated in Figure S12. Scale bars: 5 $\mu$m.

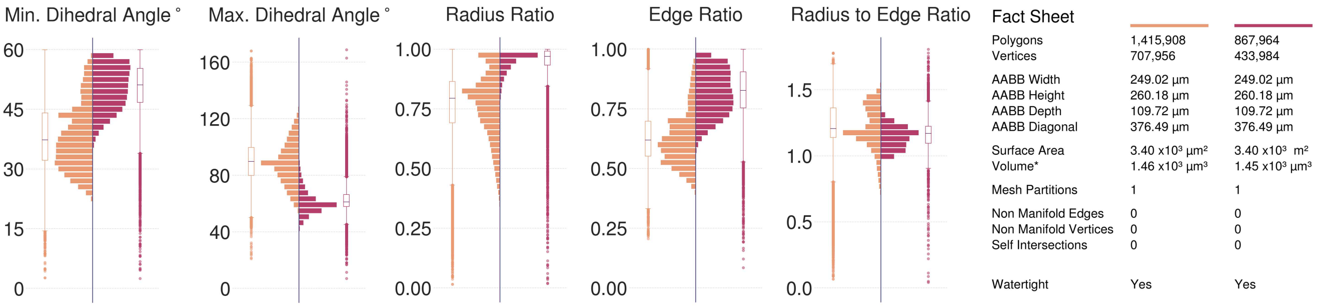

Figure S12: Comparative quantitative and qualitative analyses of the surface mesh models of the L1_LAC neuron visualized in Figure S11.

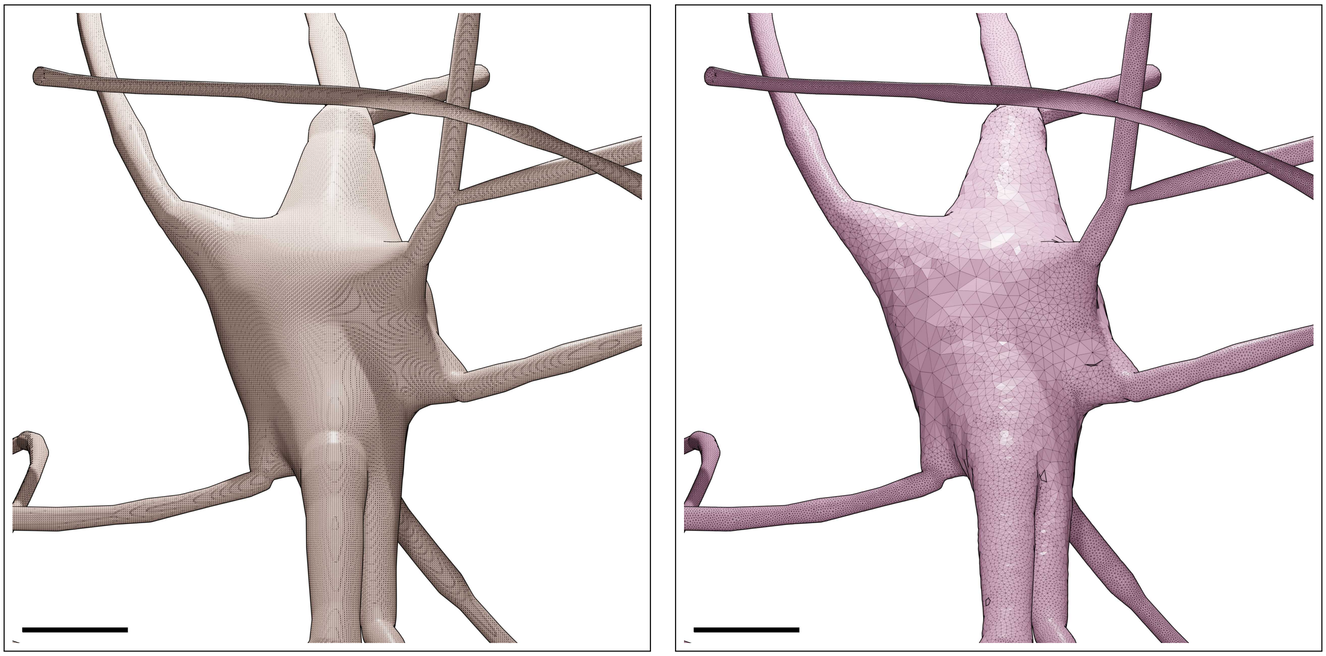

Figure S13: Wireframe visualizations of an L1_NGC-DA neuron showing closeup comparisons between the highly tessellated surface mesh generated from the Voxel remesher (left) and the adaptively optimized surface mesh generated from the optimizer (right). Comparative quantitative and qualitative analyses of the meshes are demonstrated in Figure S14. Scale bars: 5 $\mu$m.

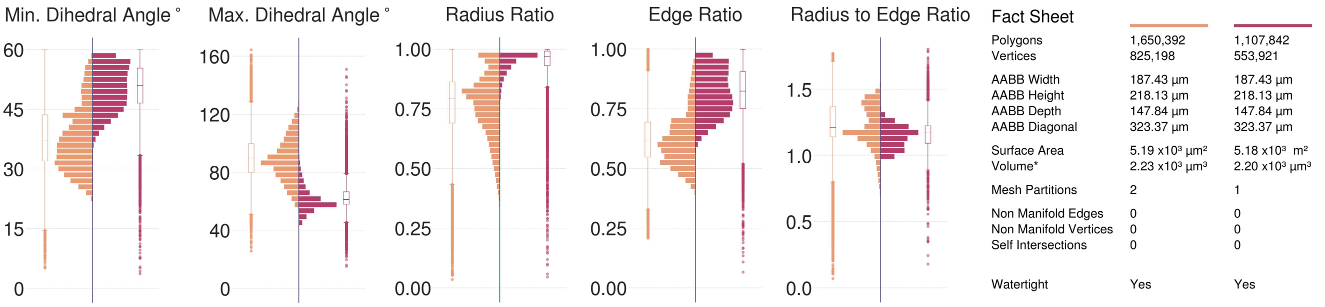

Figure S14: Comparative quantitative and qualitative analyses of the surface mesh models of the L1_NGC-DA neuron visualized in Figure S13.

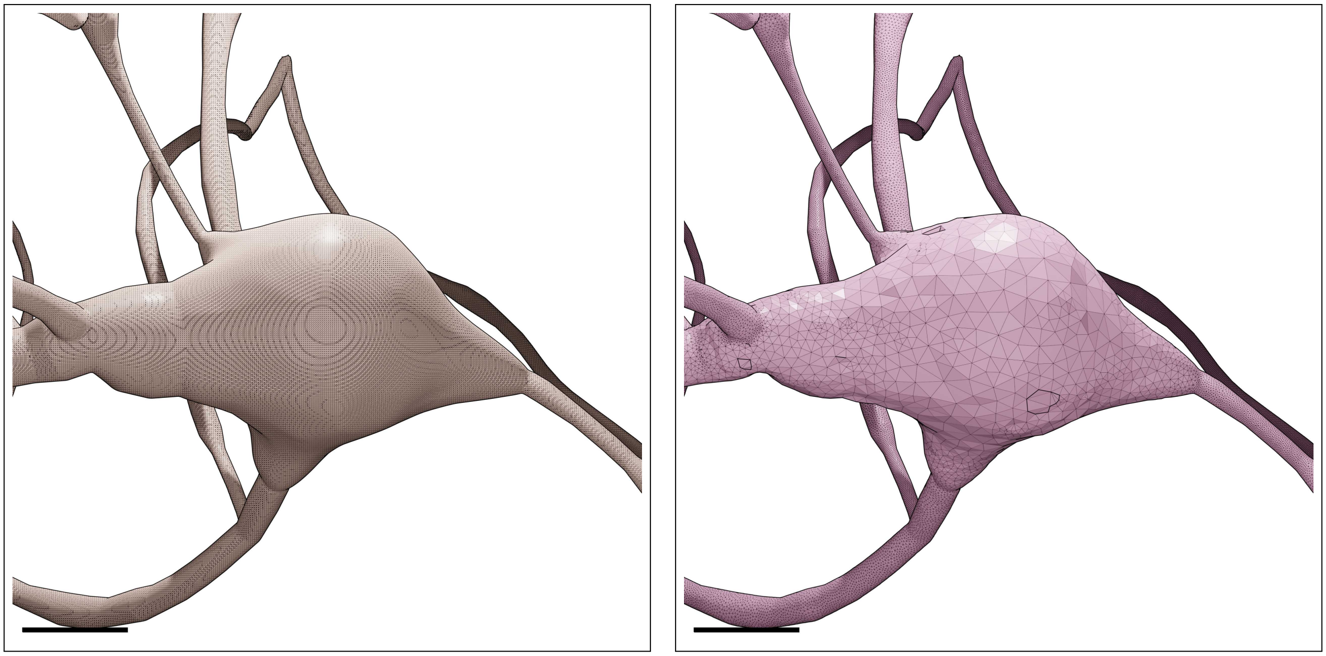

Figure S15: Wireframe visualizations of an L1_NGC-SA neuron showing closeup comparisons between the highly tessellated surface mesh generated from the Voxel remesher (left) and the adaptively optimized surface mesh generated from the optimizer (right). Comparative quantitative and qualitative analyses of the meshes are demonstrated in Figure S16. Scale bars: 5 $\mu$m.

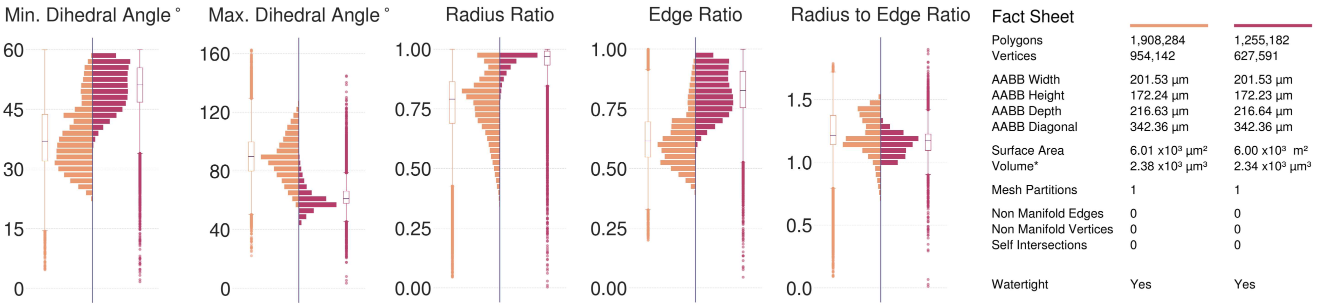

Figure S16: Comparative quantitative and qualitative analyses of the surface mesh models of the L1_NGC-SA neuron visualized in Figure S15.

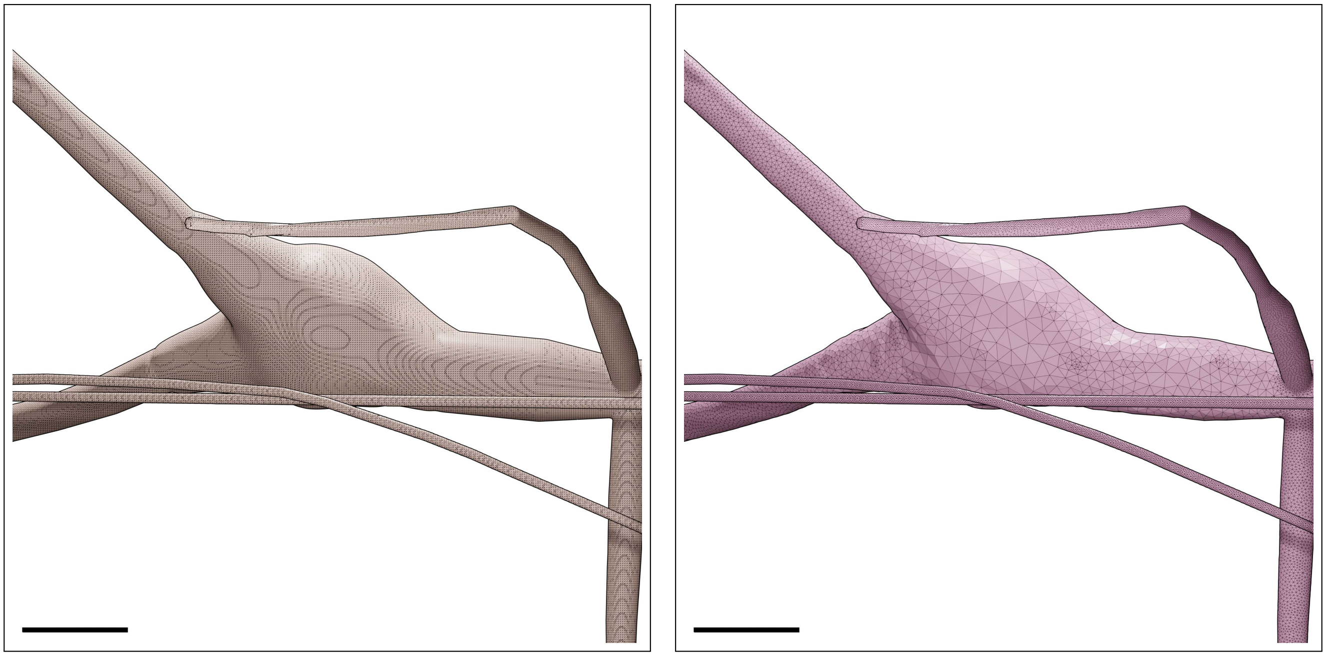

Figure S17: Wireframe visualizations of an L1_SAC neuron showing closeup comparisons between the highly tessellated surface mesh generated from the Voxel remesher (left) and the adaptively optimized surface mesh generated from the optimizer (right). Comparative quantitative and qualitative analyses of the meshes are demonstrated in Figure S18. Scale bars: 5 $\mu$m.

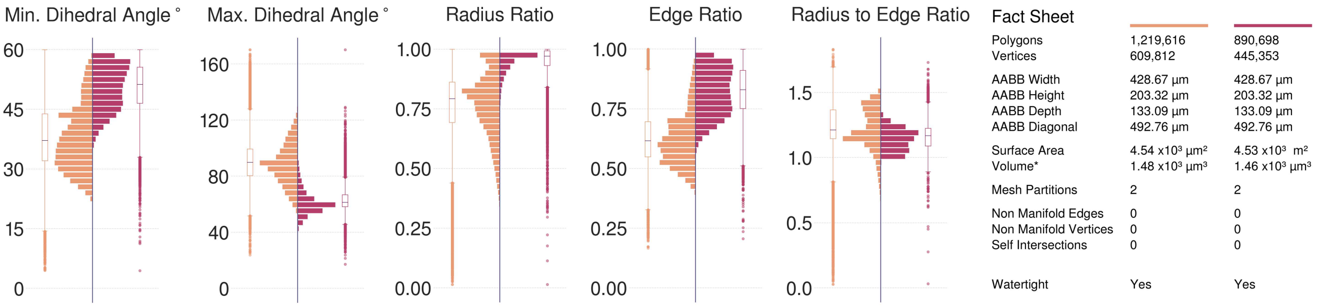

Figure S18: Comparative quantitative and qualitative analyses of the surface mesh models of the L1_SAC neuron visualized in Figure S17.

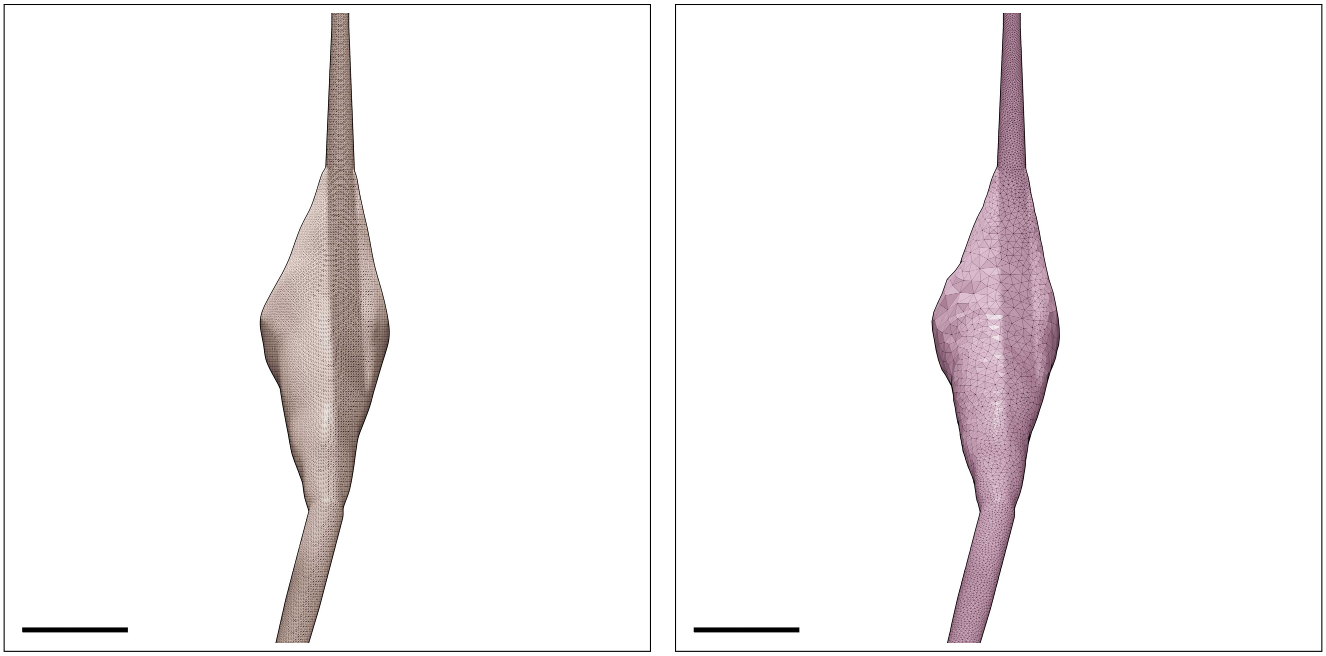

Figure S19: Wireframe visualizations of an L23_BP neuron showing closeup comparisons between the highly tessellated surface mesh generated from the Voxel remesher (left) and the adaptively optimized surface mesh generated from the optimizer (right). Comparative quantitative and qualitative analyses of the meshes are demonstrated in Figure S20. Scale bars: 5 $\mu$m.

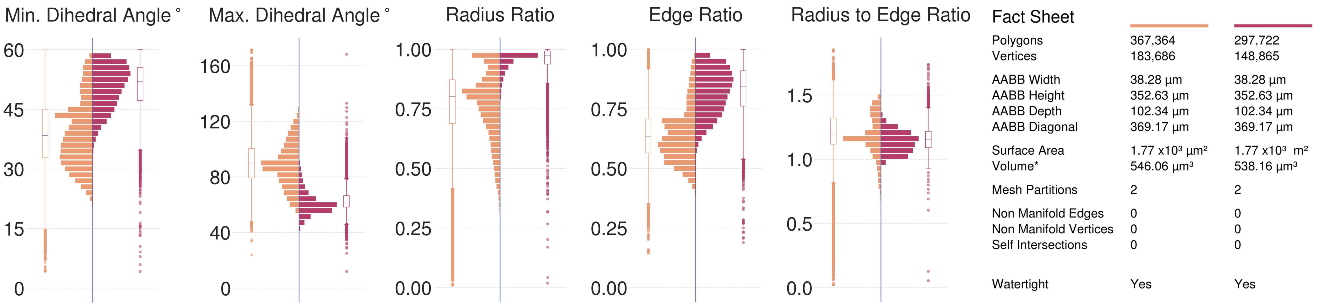

Figure S20: Comparative quantitative and qualitative analyses of the surface mesh models of the L23_BP neuron visualized in Figure S19.

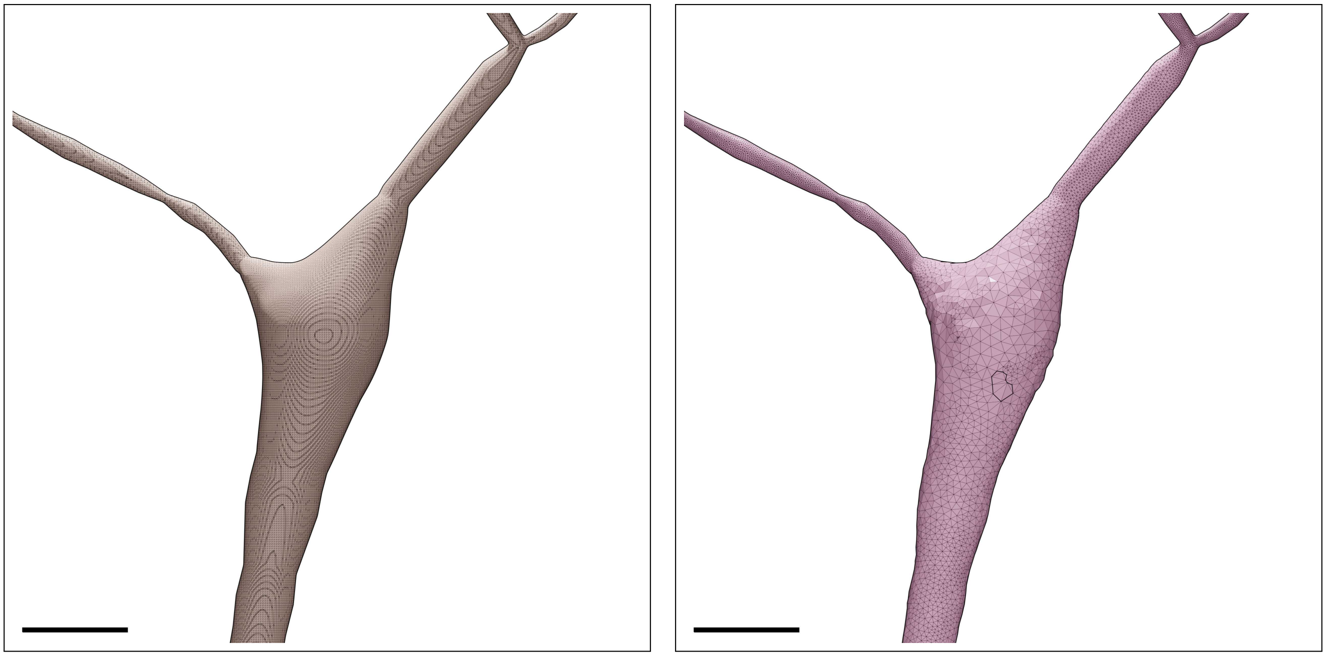

Figure S21: Wireframe visualizations of an L23_BTC neuron showing closeup comparisons between the highly tessellated surface mesh generated from the Voxel remesher (left) and the adaptively optimized surface mesh generated from the optimizer (right). Comparative quantitative and qualitative analyses of the meshes are demonstrated in Figure S22. Scale bars: 5 $\mu$m.

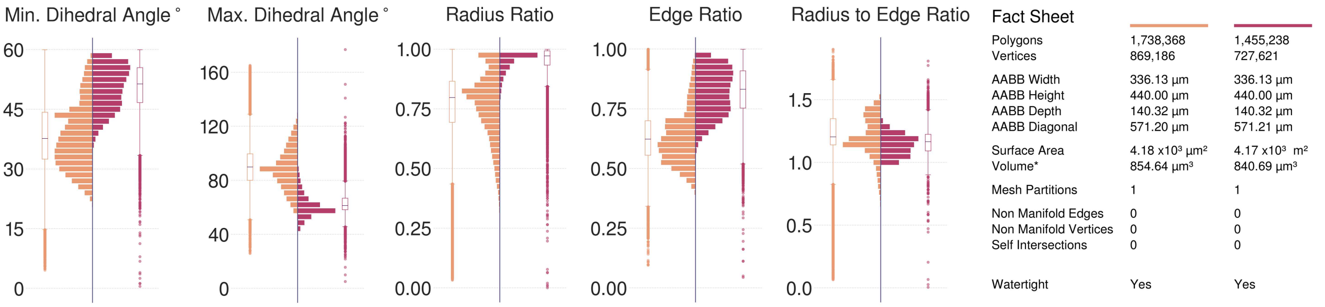

Figure S22: Comparative quantitative and qualitative analyses of the surface mesh models of the L23_BTC neuron visualized in Figure S21.

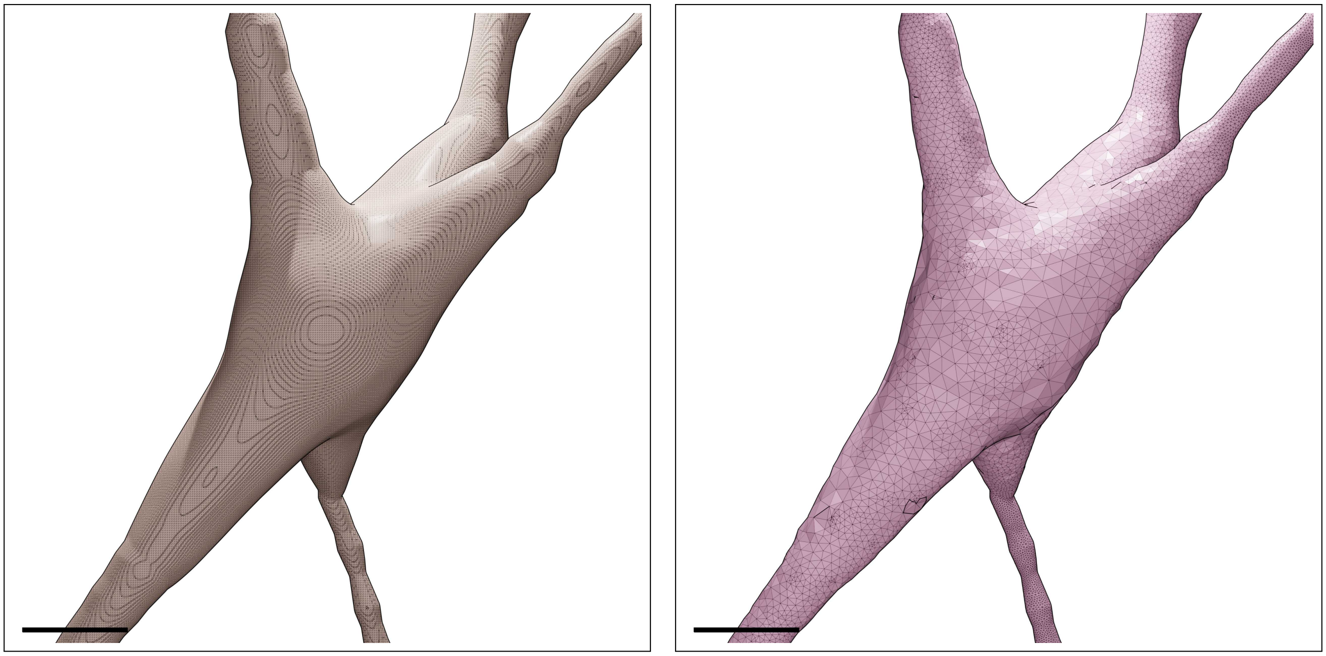

Figure S23: Wireframe visualizations of an L23_CHC neuron showing closeup comparisons between the highly tessellated surface mesh generated from the Voxel remesher (left) and the adaptively optimized surface mesh generated from the optimizer (right). Comparative quantitative and qualitative analyses of the meshes are demonstrated in Figure S24. Scale bars: 5 $\mu$m.

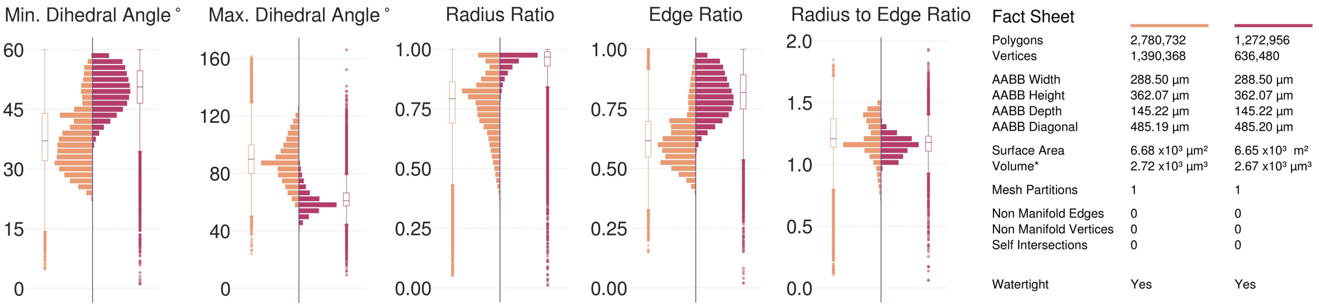

Figure S24: Comparative quantitative and qualitative analyses of the surface mesh models of the L23_CHC neuron visualized in Figure S23.

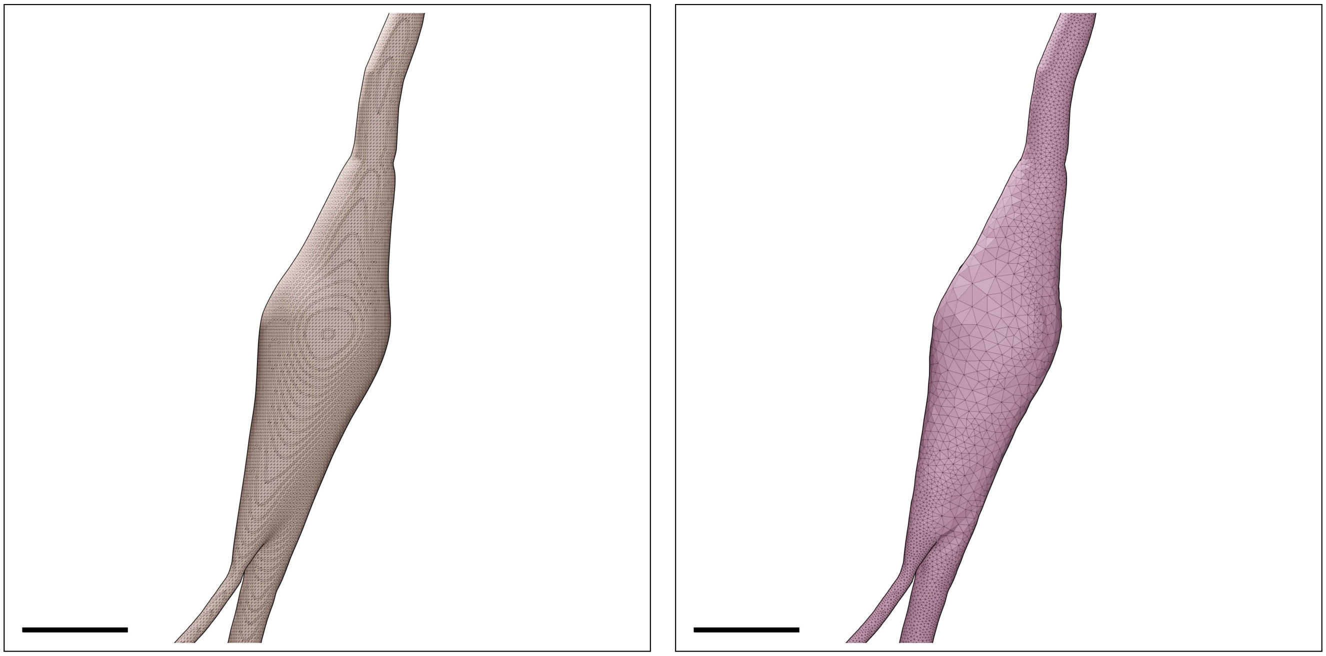

Figure S25: Wireframe visualizations of an L23_DBC neuron showing closeup comparisons between the highly tessellated surface mesh generated from the Voxel remesher (left) and the adaptively optimized surface mesh generated from the optimizer (right). Comparative quantitative and qualitative analyses of the meshes are demonstrated in Figure S26. Scale bars: 5 $\mu$m.

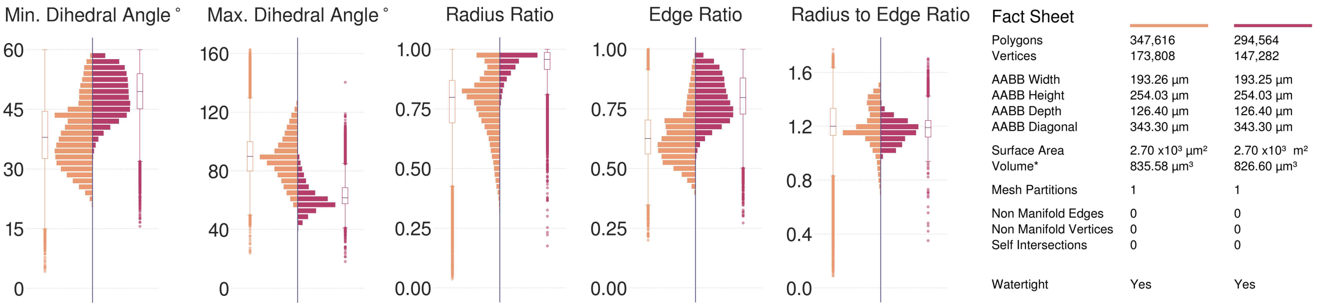

Figure S26: Comparative quantitative and qualitative analyses of the surface mesh models of the L23_DBC neuron visualized in Figure S25.

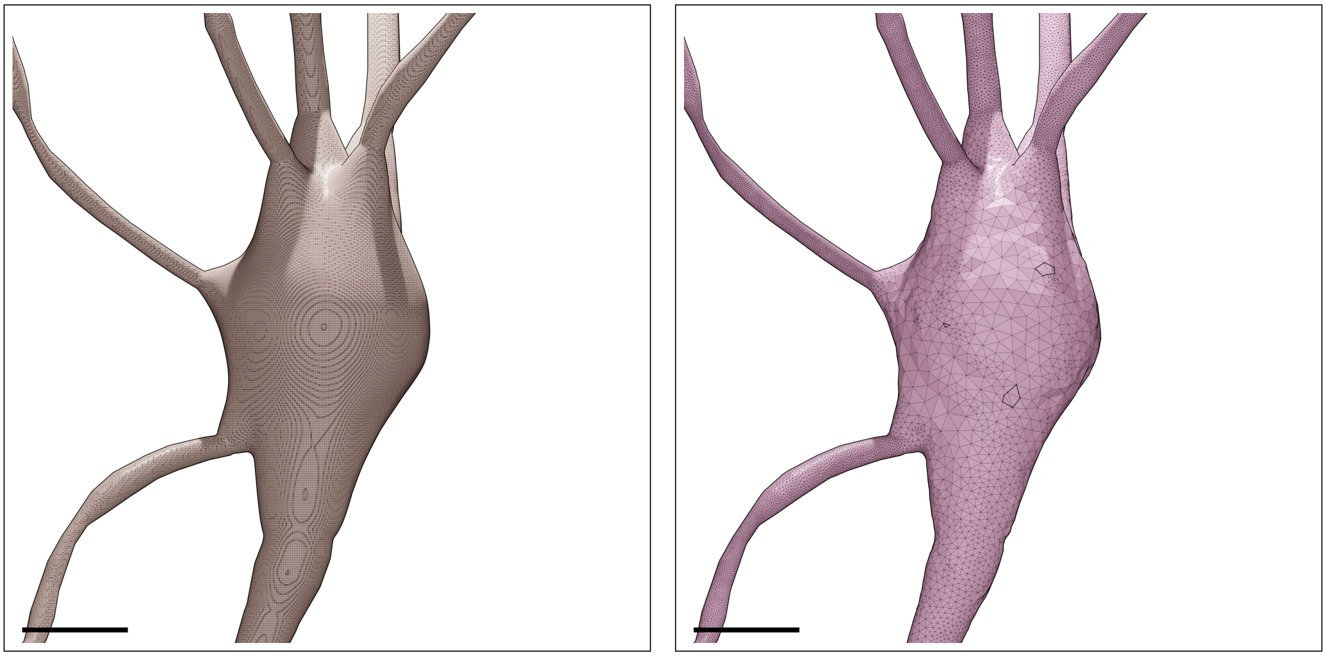

Figure S27: Wireframe visualizations of an L23_LBC neuron showing closeup comparisons between the highly tessellated surface mesh generated from the Voxel remesher (left) and the adaptively optimized surface mesh generated from the optimizer (right). Comparative quantitative and qualitative analyses of the meshes are demonstrated in Figure S28. Scale bars: 5 $\mu$m.

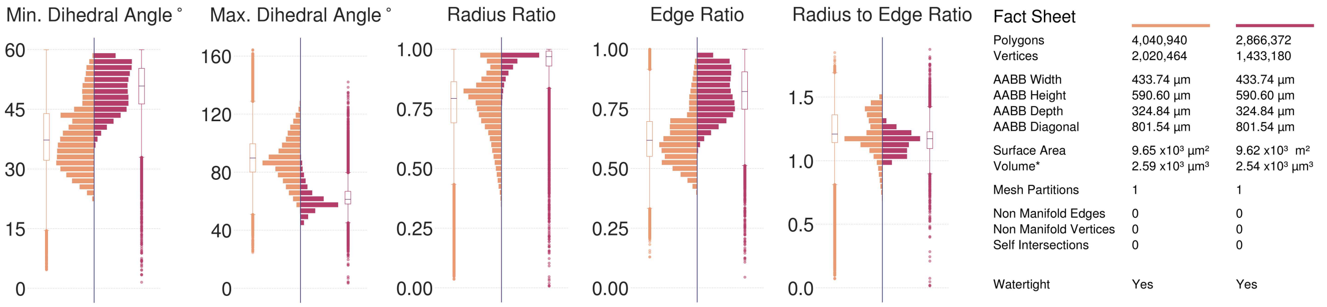

Figure S28: Comparative quantitative and qualitative analyses of the surface mesh models of the L23_LBC neuron visualized in Figure S27.

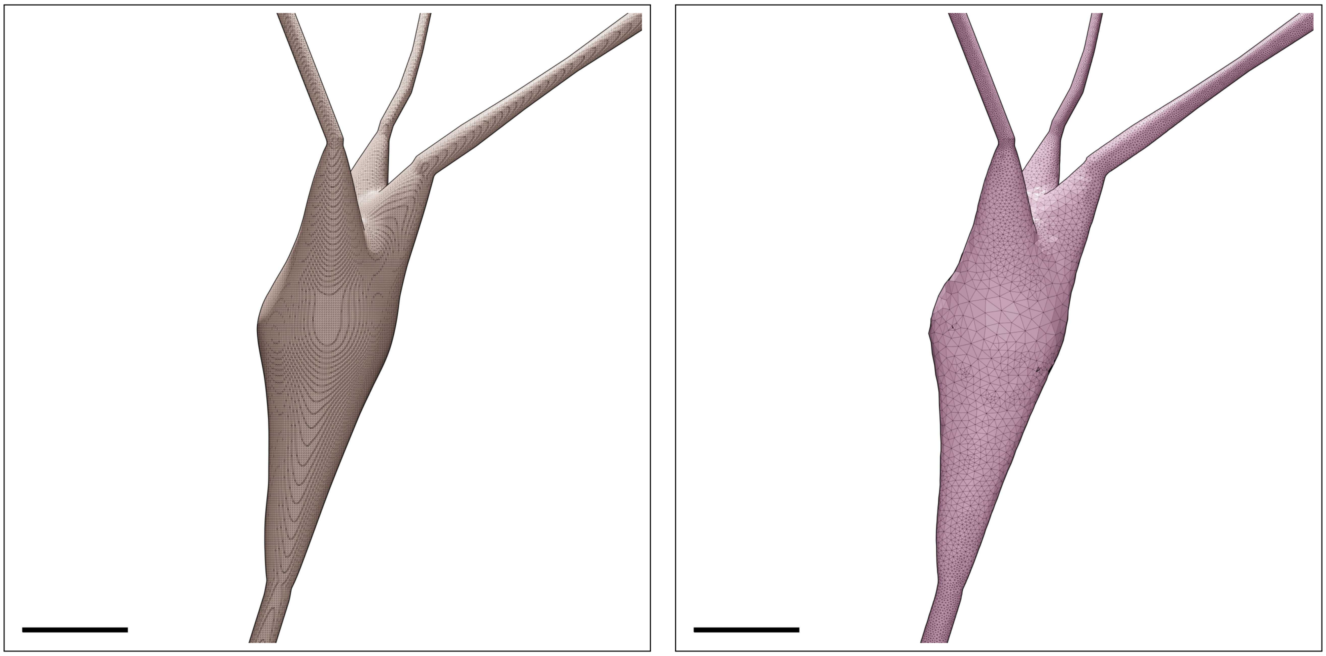

Figure S29: Wireframe visualizations of an L23_MC neuron showing closeup comparisons between the highly tessellated surface mesh generated from the Voxel remesher (left) and the adaptively optimized surface mesh generated from the optimizer (right). Comparative quantitative and qualitative analyses of the meshes are demonstrated in Figure S30. Scale bars: 5 $\mu$m.

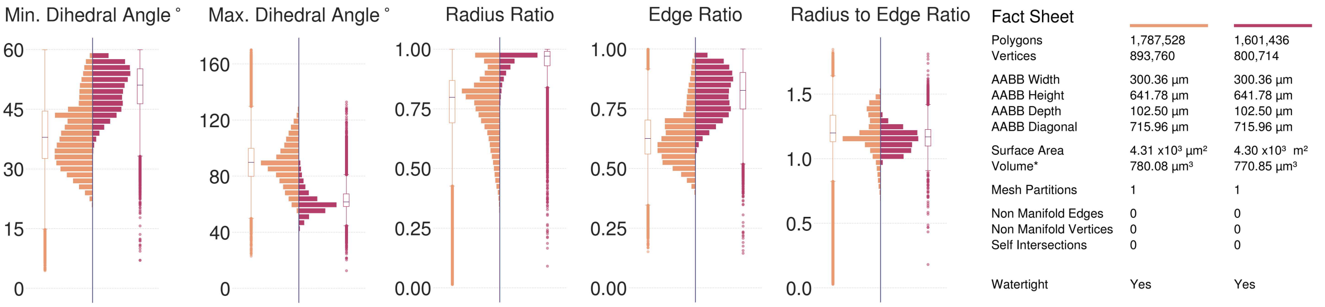

Figure S30: Comparative quantitative and qualitative analyses of the surface mesh models of the L23_MC neuron visualized in Figure S29.

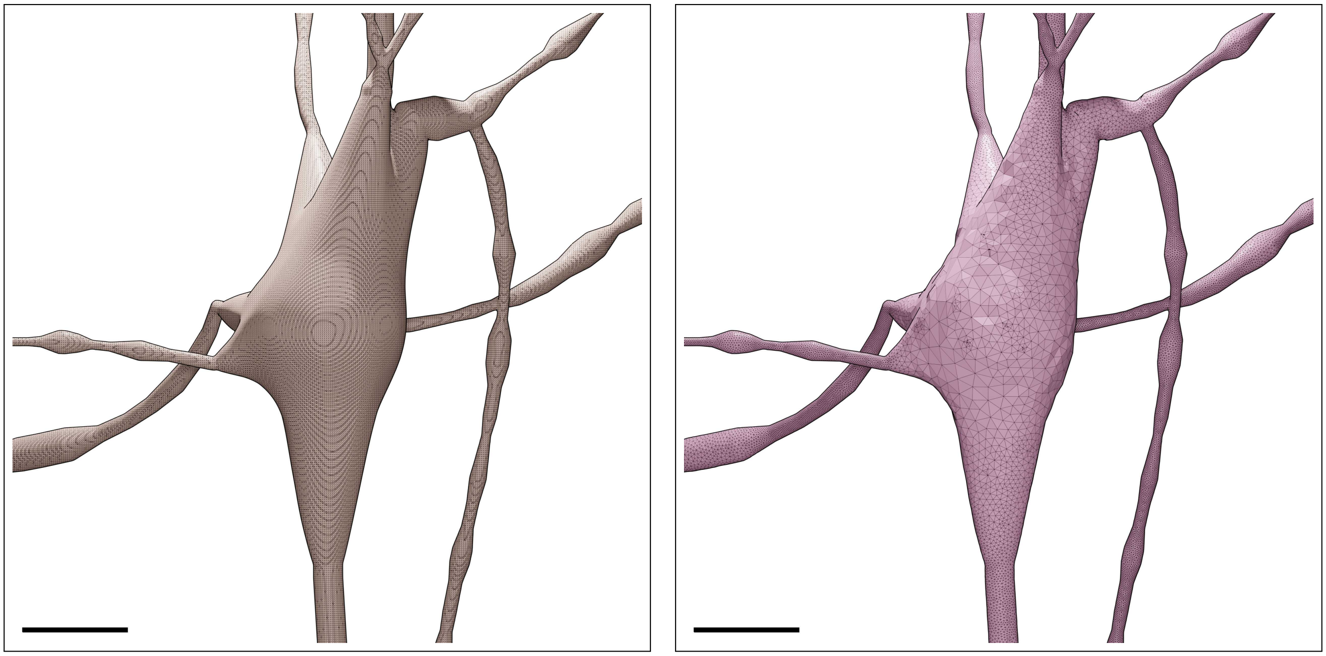

Figure S31: Wireframe visualizations of an L23_NBC neuron showing closeup comparisons between the highly tessellated surface mesh generated from the Voxel remesher (left) and the adaptively optimized surface mesh generated from the optimizer (right). Comparative quantitative and qualitative analyses of the meshes are demonstrated in Figure S32. Scale bars: 5 $\mu$m.

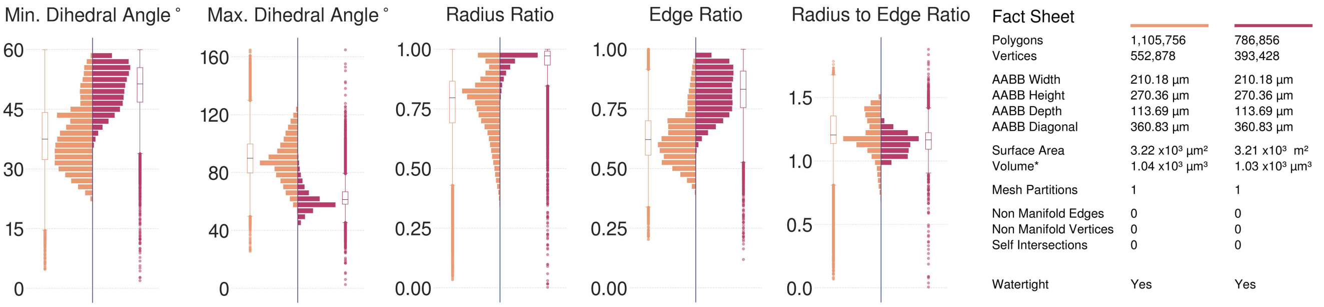

Figure S32: Comparative quantitative and qualitative analyses of the surface mesh models of the L23_NBC neuron visualized in Figure S31.

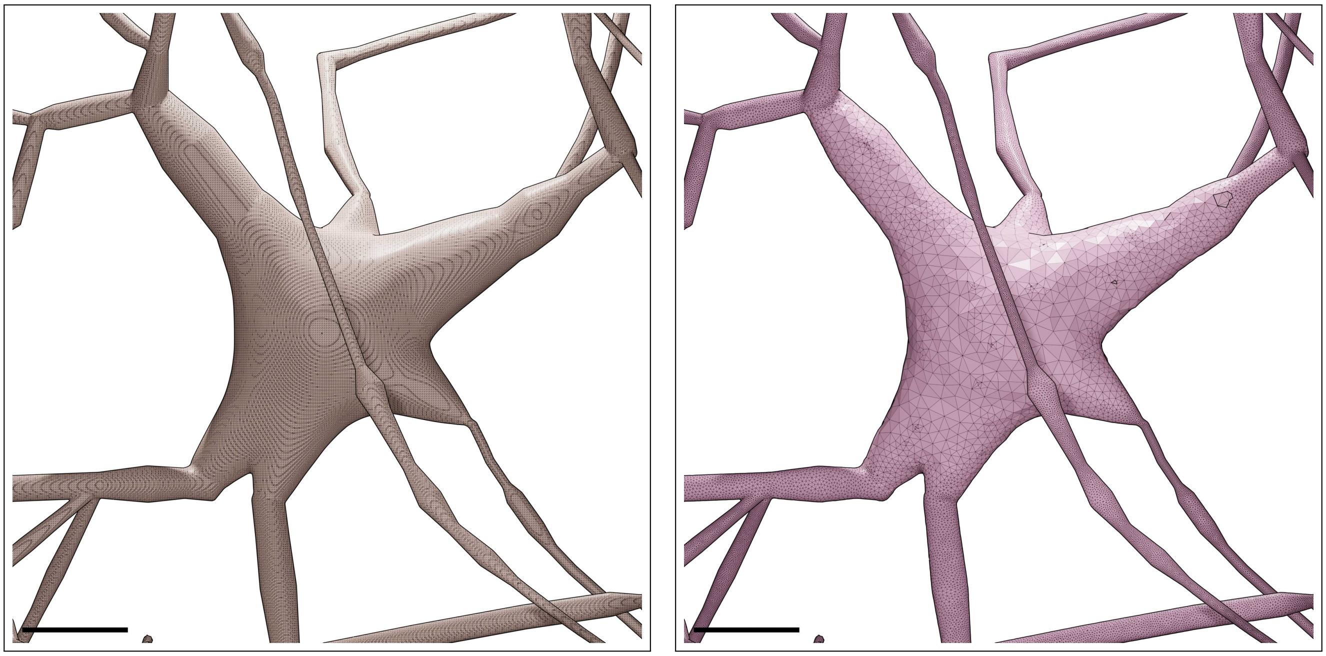

Figure S33: Wireframe visualizations of an L23_NGC neuron showing closeup comparisons between the highly tessellated surface mesh generated from the Voxel remesher (left) and the adaptively optimized surface mesh generated from the optimizer (right). Comparative quantitative and qualitative analyses of the meshes are demonstrated in Figure S34. Scale bars: 5 $\mu$m.

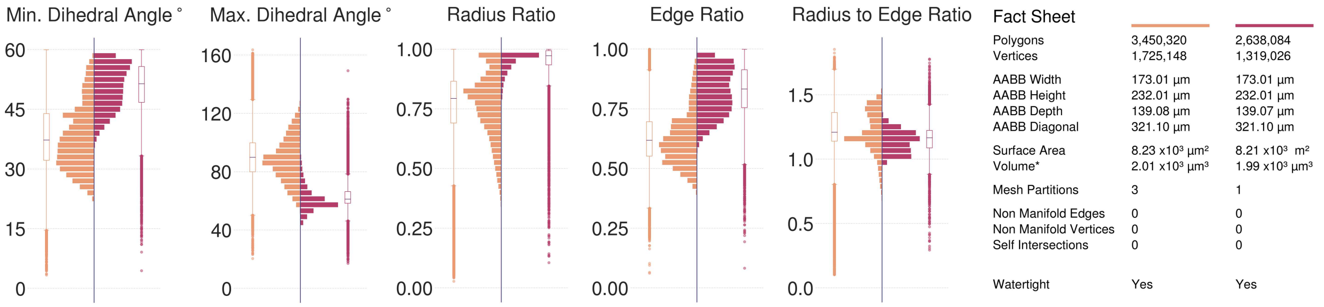

Figure S34: Comparative quantitative and qualitative analyses of the surface mesh models of the L23_NGC neuron visualized in Figure S33.

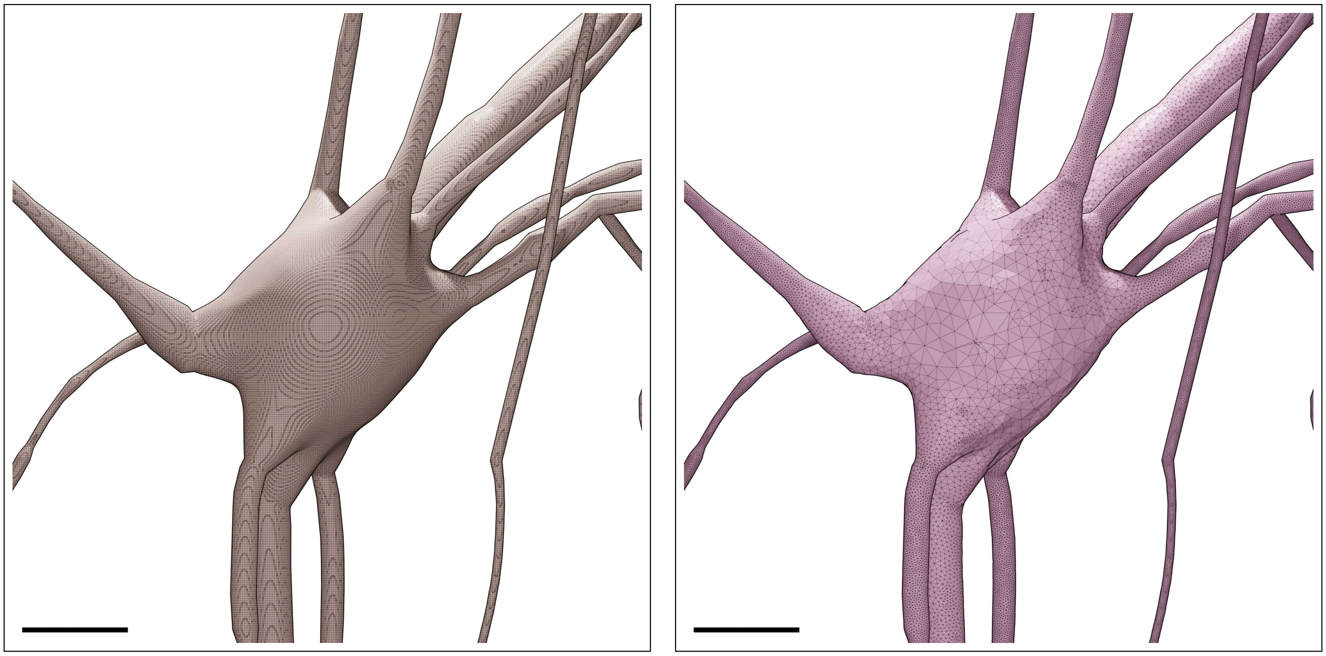

Figure S35: Wireframe visualizations of an L23_SBC neuron showing closeup comparisons between the highly tessellated surface mesh generated from the Voxel remesher (left) and the adaptively optimized surface mesh generated from the optimizer (right). Comparative quantitative and qualitative analyses of the meshes are demonstrated in Figure S36. Scale bars: 5 $\mu$m.

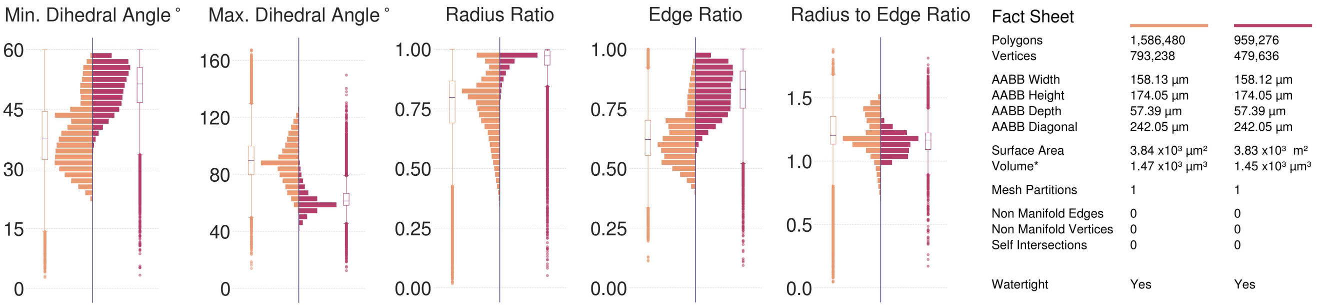

Figure S36: Comparative quantitative and qualitative analyses of the surface mesh models of the L23_SBC neuron visualized in Figure S35.

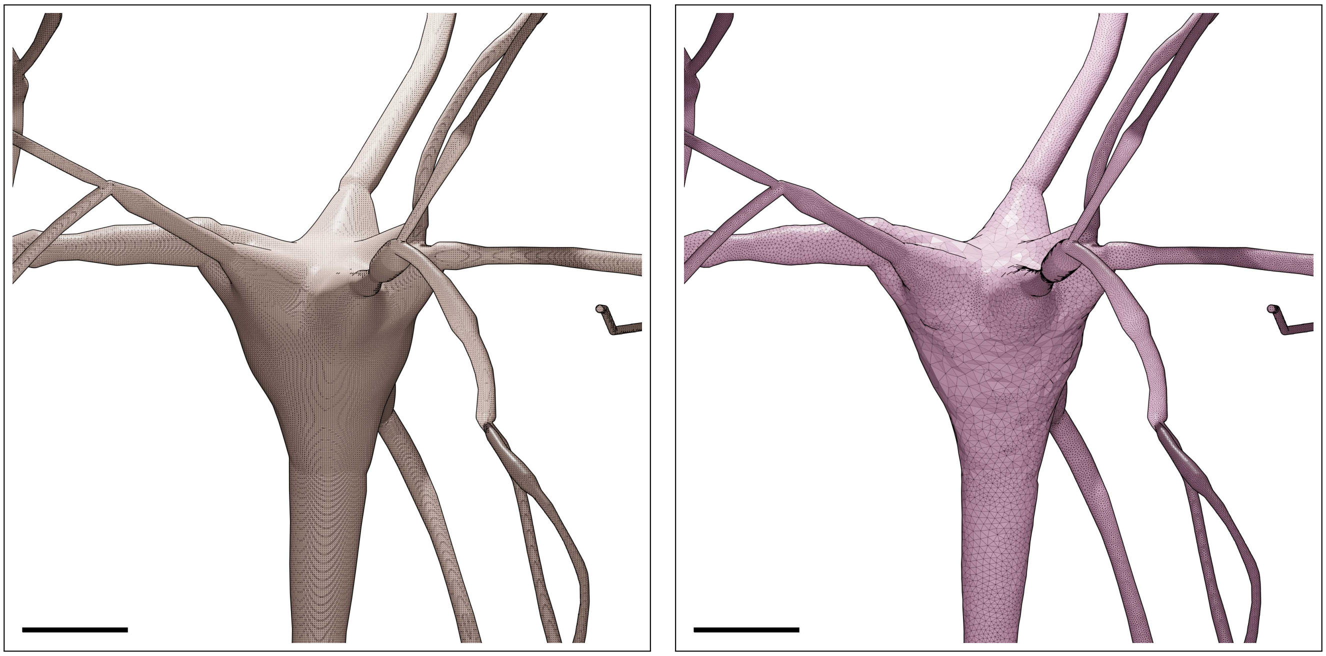

Figure S37: Wireframe visualizations of an L2_IPC neuron showing closeup comparisons between the highly tessellated surface mesh generated from the Voxel remesher (left) and the adaptively optimized surface mesh generated from the optimizer (right). Comparative quantitative and qualitative analyses of the meshes are demonstrated in Figure S38. Scale bars: 5 $\mu$m.

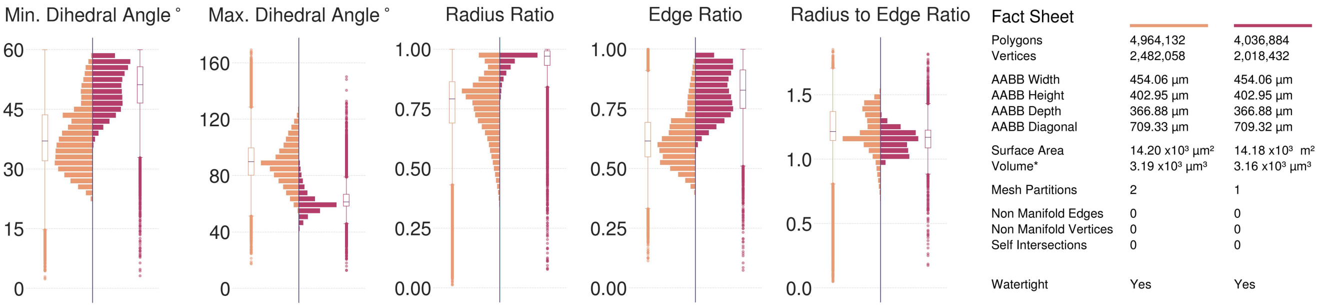

Figure S38: Comparative quantitative and qualitative analyses of the surface mesh models of the L2_IPC neuron visualized in Figure S37.

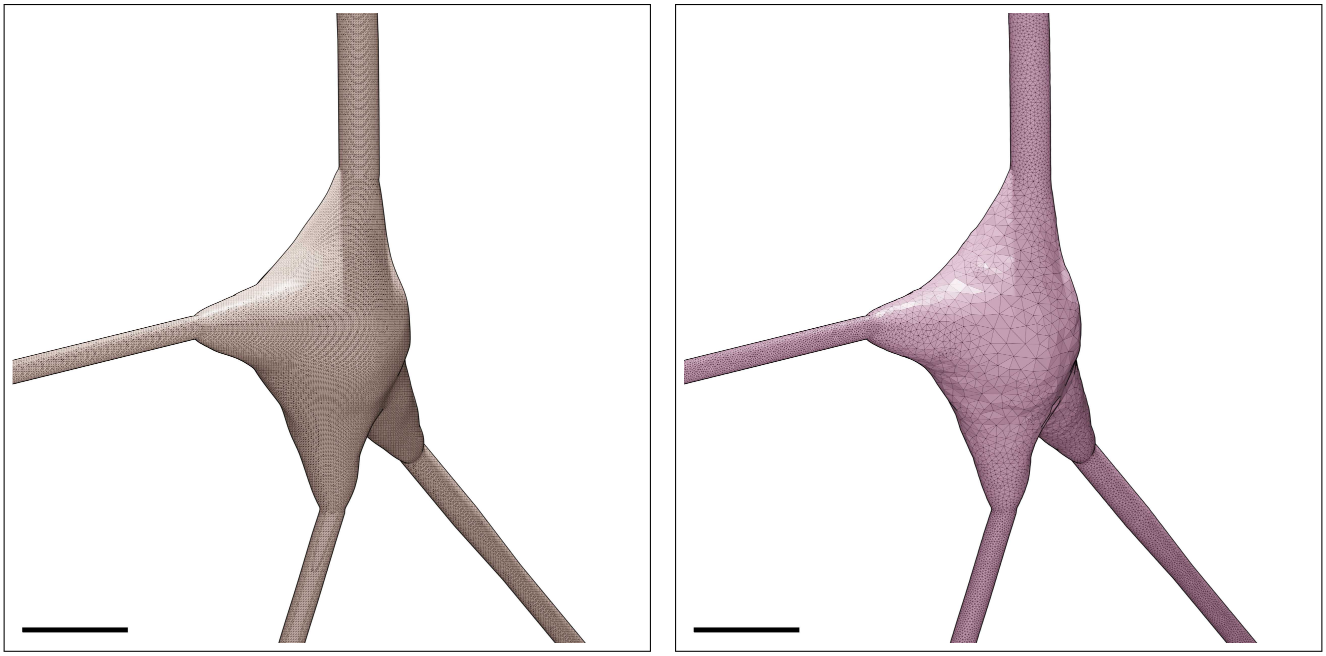

Figure S39: Wireframe visualizations of an L2_TPC:A neuron showing closeup comparisons between the highly tessellated surface mesh generated from the Voxel remesher (left) and the adaptively optimized surface mesh generated from the optimizer (right). Comparative quantitative and qualitative analyses of the meshes are demonstrated in Figure S40. Scale bars: 5 $\mu$m.

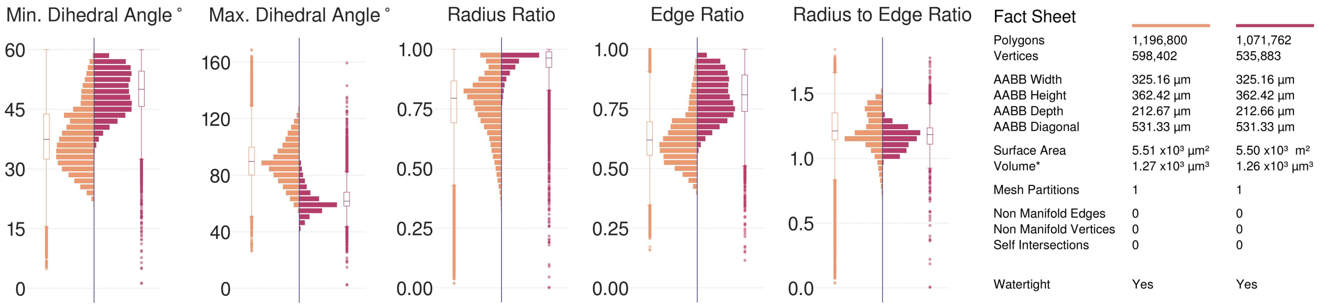

Figure S40: Comparative quantitative and qualitative analyses of the surface mesh models of the L2_TPC:A neuron visualized in Figure S39.

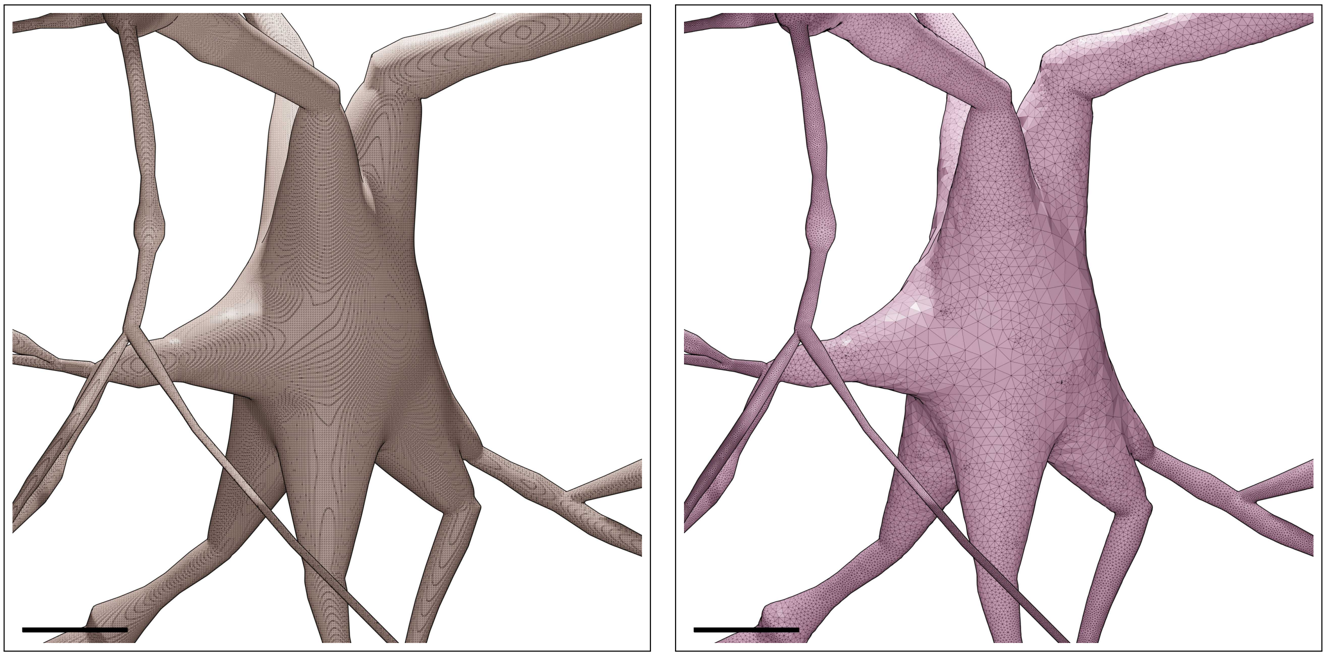

Figure S41: Wireframe visualizations of an L2_TPC:B neuron showing closeup comparisons between the highly tessellated surface mesh generated from the Voxel remesher (left) and the adaptively optimized surface mesh generated from the optimizer (right). Comparative quantitative and qualitative analyses of the meshes are demonstrated in Figure S42. Scale bars: 5 $\mu$m.

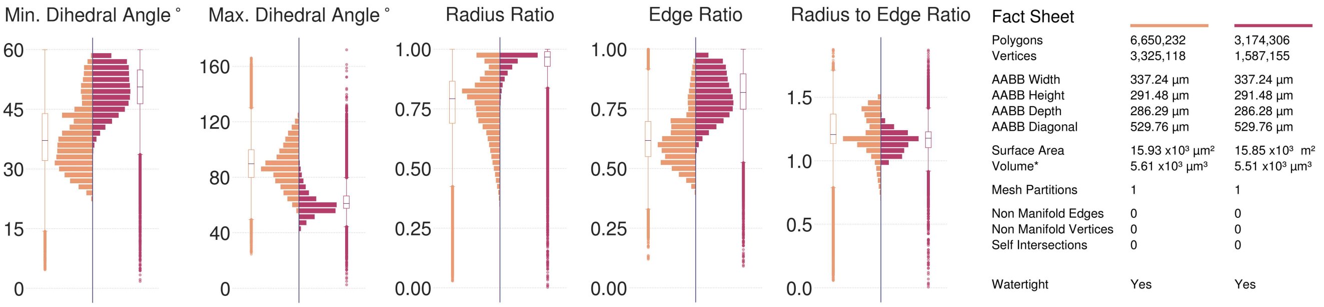

Figure S42: Comparative quantitative and qualitative analyses of the surface mesh models of the L2_TPC:B neuron visualized in Figure S41.

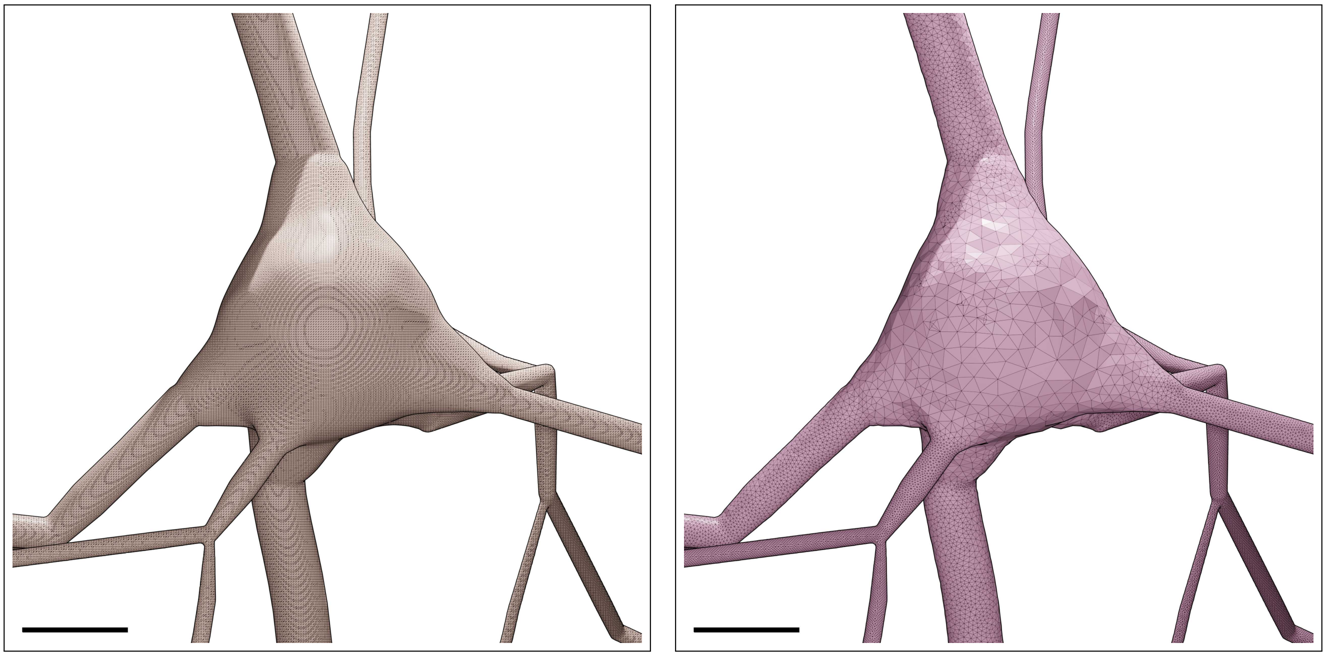

Figure S43: Wireframe visualizations of an L3_TPC:A neuron showing closeup comparisons between the highly tessellated surface mesh generated from the Voxel remesher (left) and the adaptively optimized surface mesh generated from the optimizer (right). Comparative quantitative and qualitative analyses of the meshes are demonstrated in Figure S44. Scale bars: 5 $\mu$m.

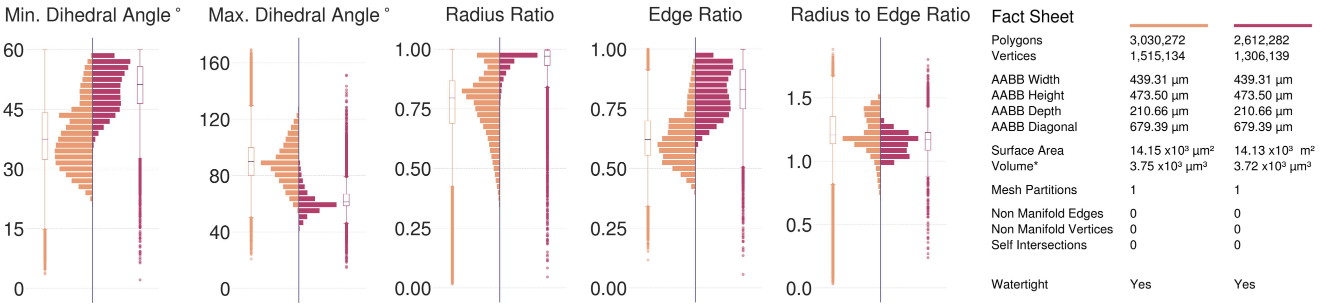

Figure S44: Comparative quantitative and qualitative analyses of the surface mesh models of the L3_TPC:A neuron visualized in Figure S43.

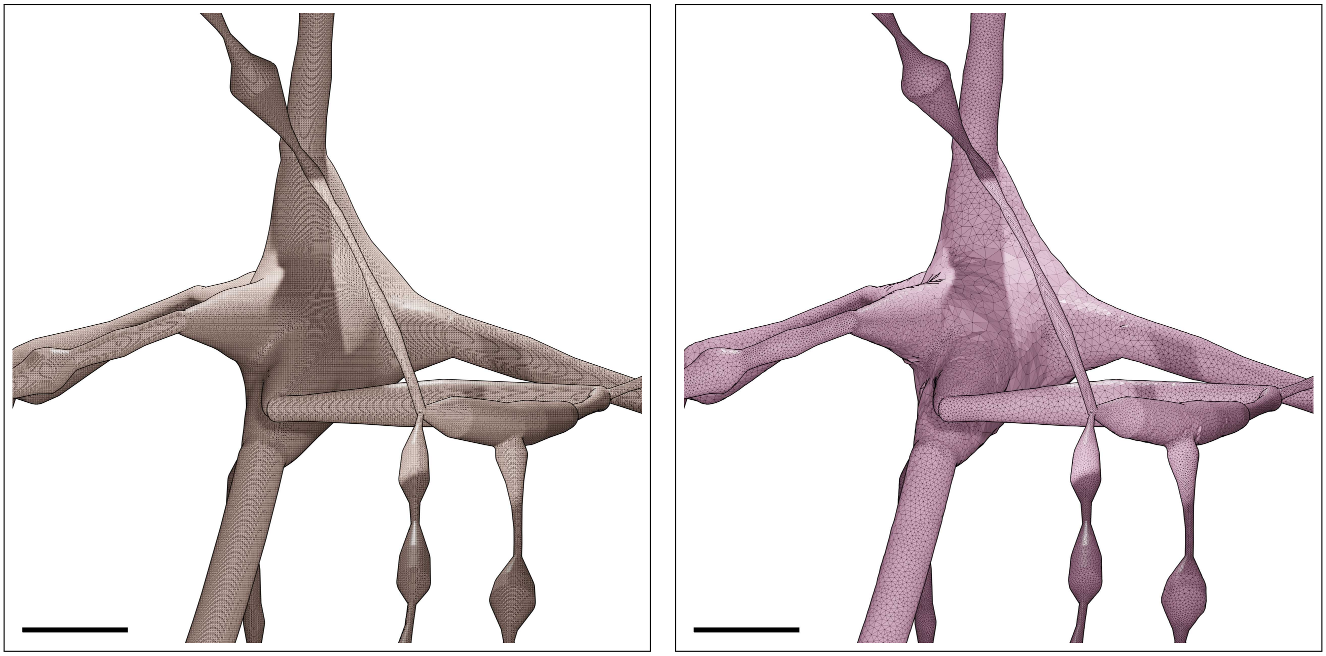

Figure S45: Wireframe visualizations of an L3_TPC:C neuron showing closeup comparisons between the highly tessellated surface mesh generated from the Voxel remesher (left) and the adaptively optimized surface mesh generated from the optimizer (right). Comparative quantitative and qualitative analyses of the meshes are demonstrated in Figure S46. Scale bars: 5 $\mu$m.

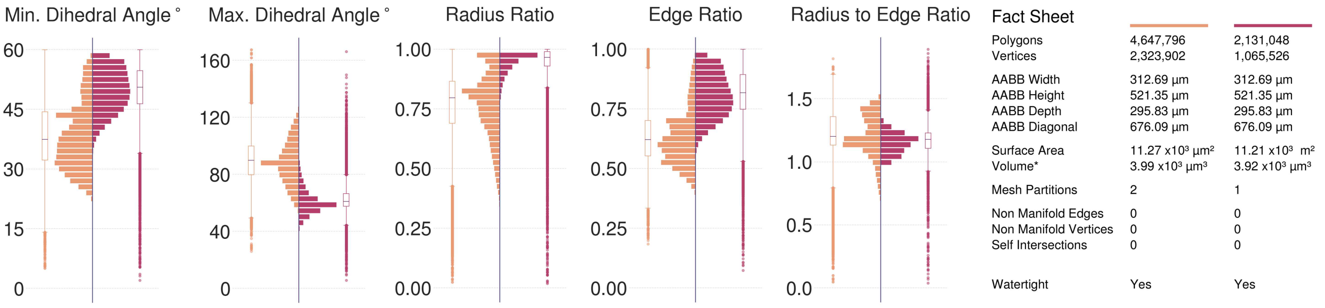

Figure S46: Comparative quantitative and qualitative analyses of the surface mesh models of the L3_TPC:C neuron visualized in Figure S45.

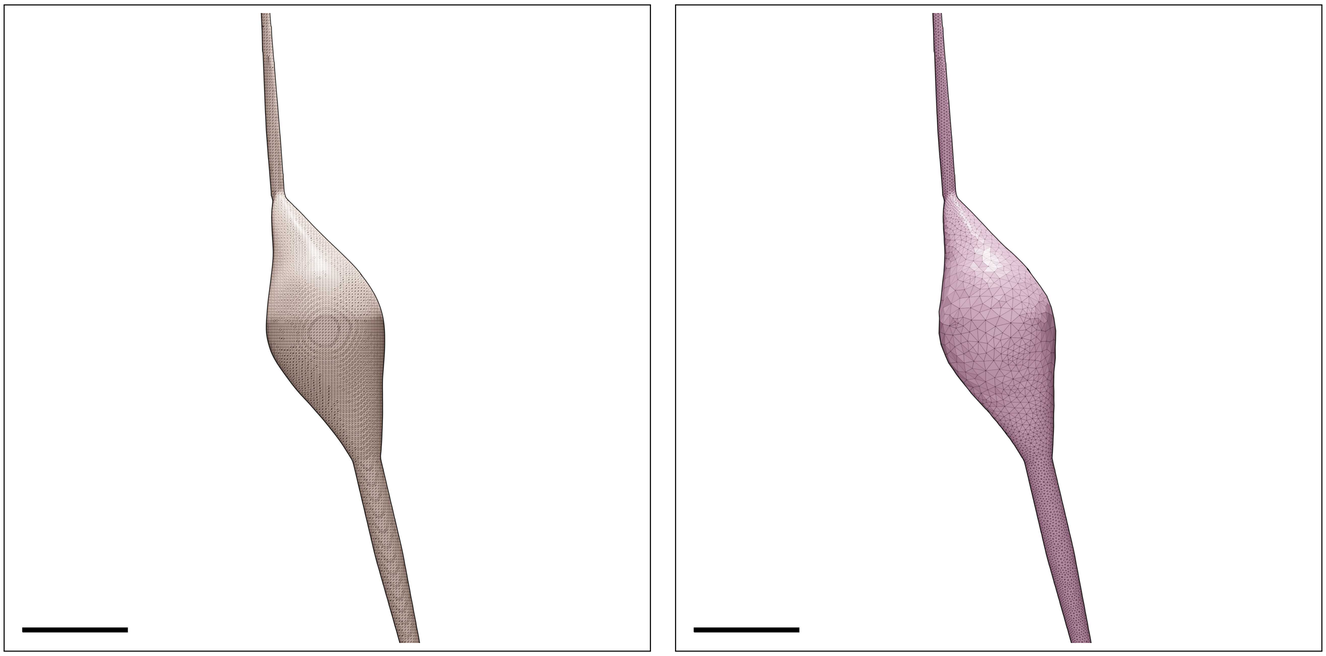

Figure S47: Wireframe visualizations of an L4_BP neuron showing closeup comparisons between the highly tessellated surface mesh generated from the Voxel remesher (left) and the adaptively optimized surface mesh generated from the optimizer (right). Comparative quantitative and qualitative analyses of the meshes are demonstrated in Figure S48. Scale bars: 5 $\mu$m.

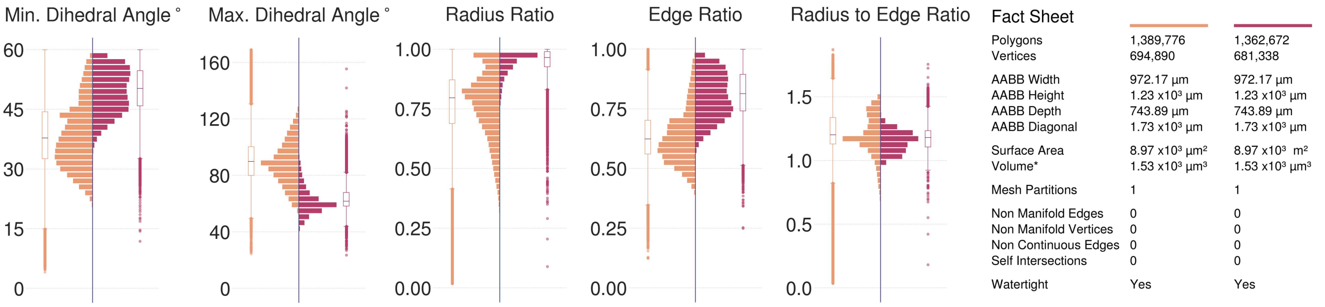

Figure S48: Comparative quantitative and qualitative analyses of the surface mesh models of the L4_BP neuron visualized in Figure S47.

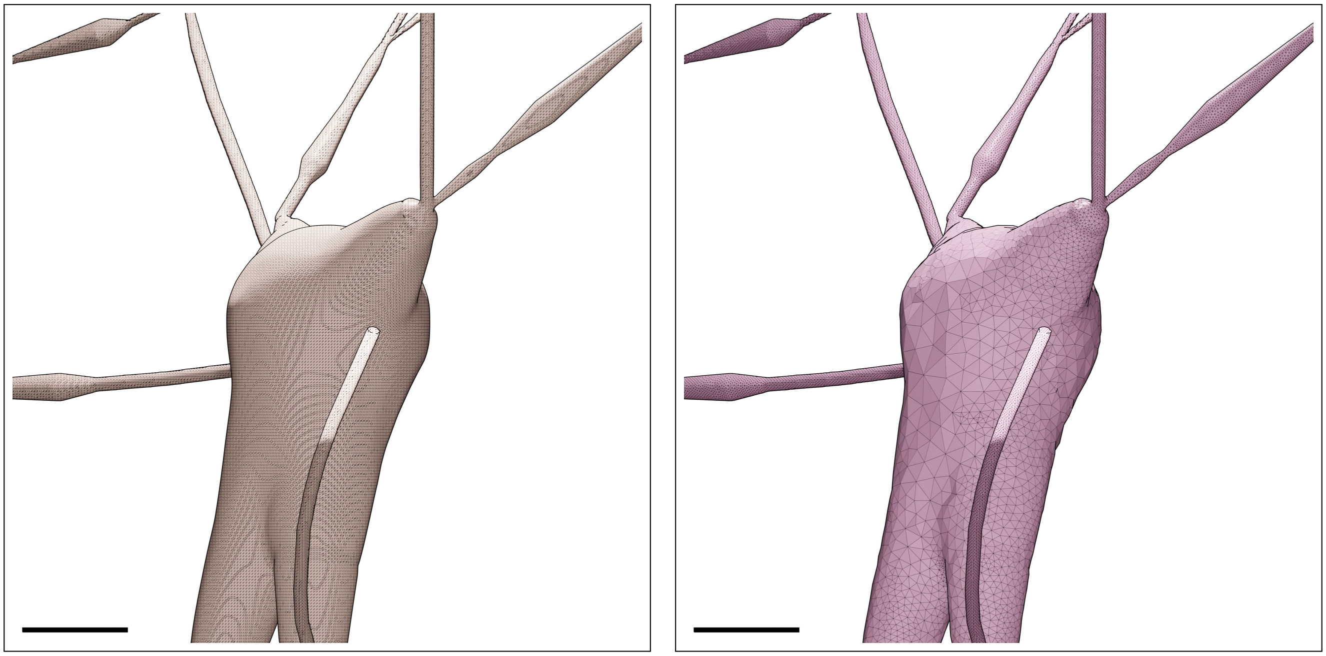

Figure S49: Wireframe visualizations of an L4_BTC neuron showing closeup comparisons between the highly tessellated surface mesh generated from the Voxel remesher (left) and the adaptively optimized surface mesh generated from the optimizer (right). Comparative quantitative and qualitative analyses of the meshes are demonstrated in Figure S50. Scale bars: 5 $\mu$m.

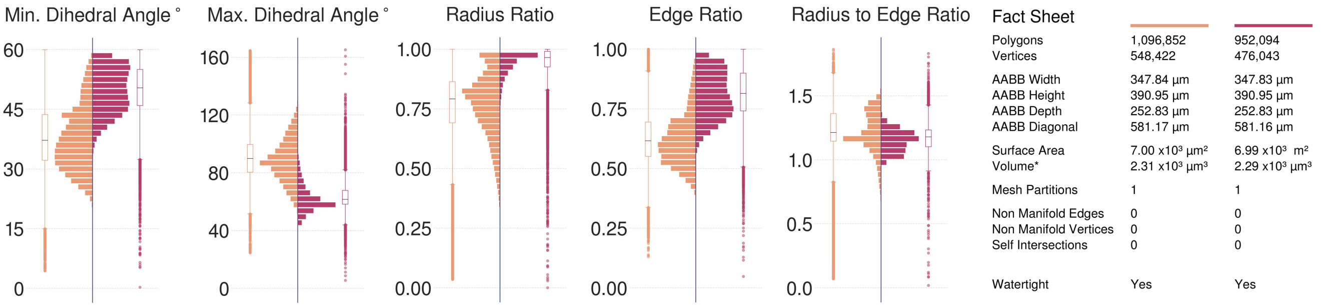

Figure S50: Comparative quantitative and qualitative analyses of the surface mesh models of the L4_BTC neuron visualized in Figure S49.

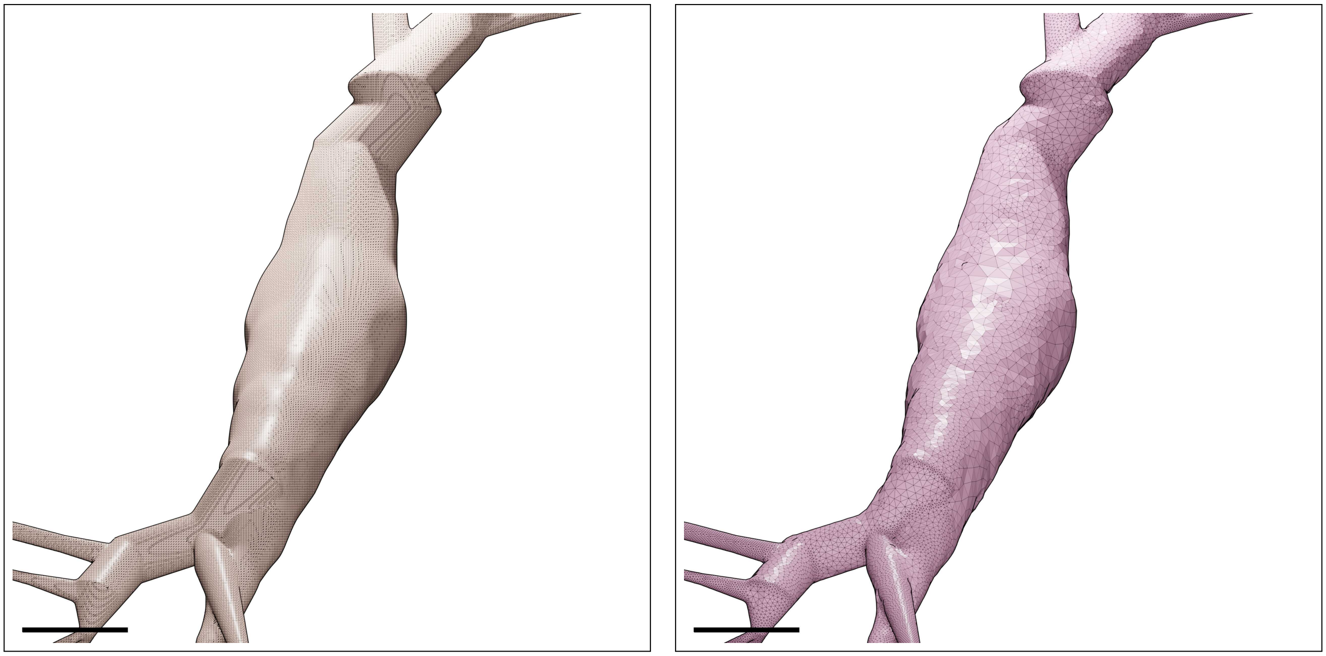

Figure S51: Wireframe visualizations of an L4_CHC neuron showing closeup comparisons between the highly tessellated surface mesh generated from the Voxel remesher (left) and the adaptively optimized surface mesh generated from the optimizer (right). Comparative quantitative and qualitative analyses of the meshes are demonstrated in Figure S52. Scale bars: 5 $\mu$m.

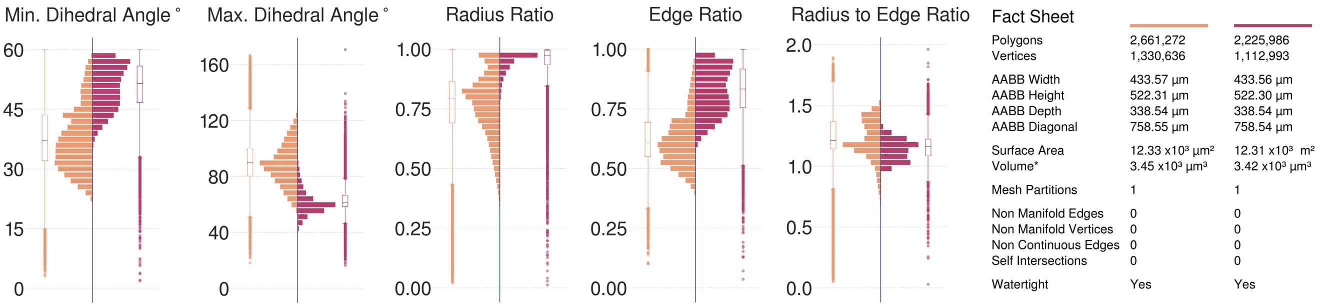

Figure S52: Comparative quantitative and qualitative analyses of the surface mesh models of the L4_CHC neuron visualized in Figure S51.

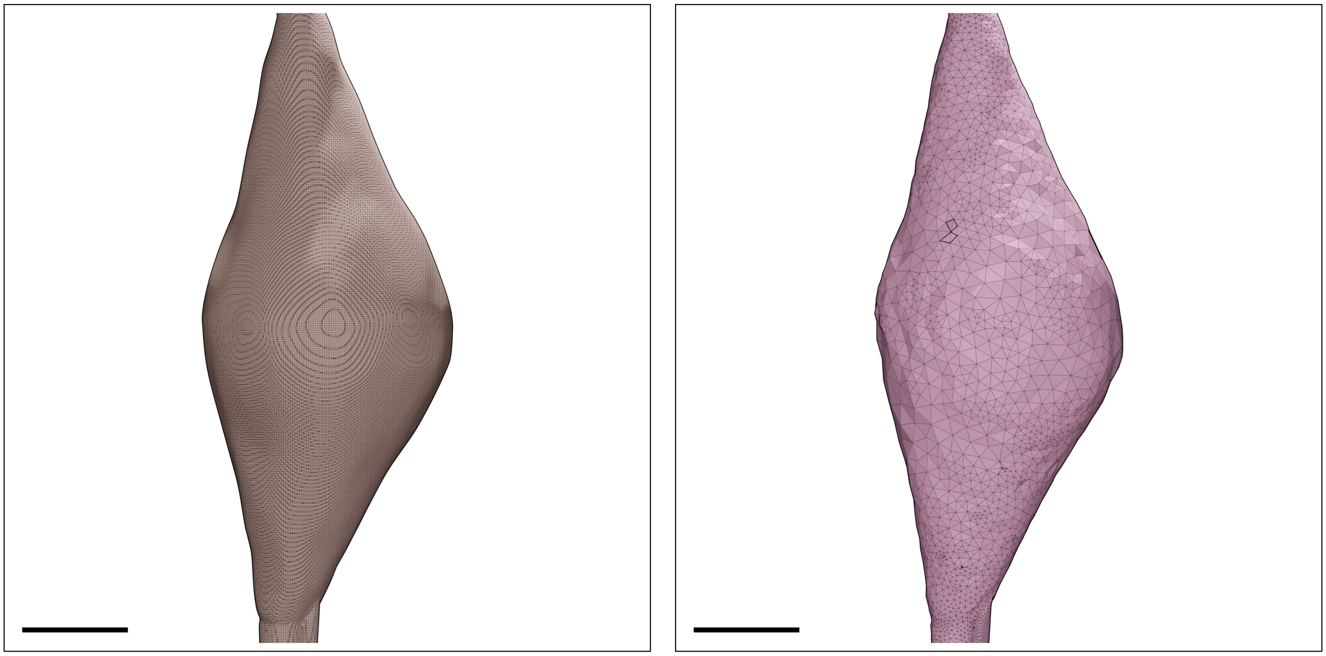

Figure S53: Wireframe visualizations of an L4_DBC neuron showing closeup comparisons between the highly tessellated surface mesh generated from the Voxel remesher (left) and the adaptively optimized surface mesh generated from the optimizer (right). Comparative quantitative and qualitative analyses of the meshes are demonstrated in Figure S54. Scale bars: 5 $\mu$m.

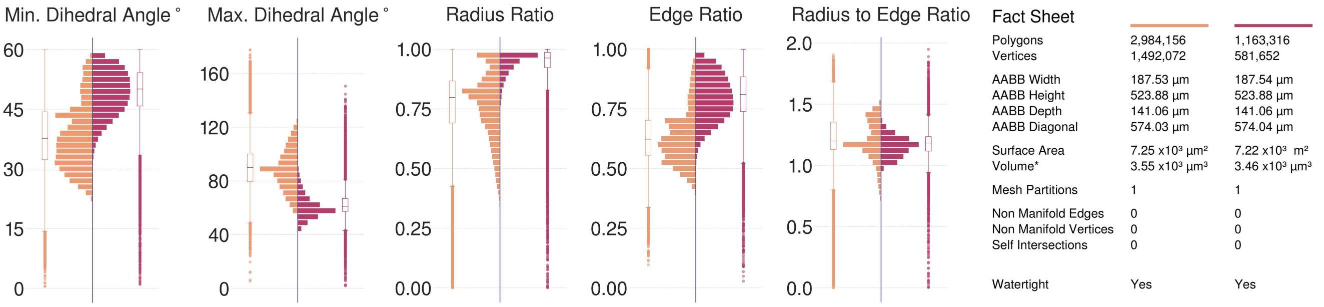

Figure S54: Comparative quantitative and qualitative analyses of the surface mesh models of the L4_DBC neuron visualized in Figure S53.

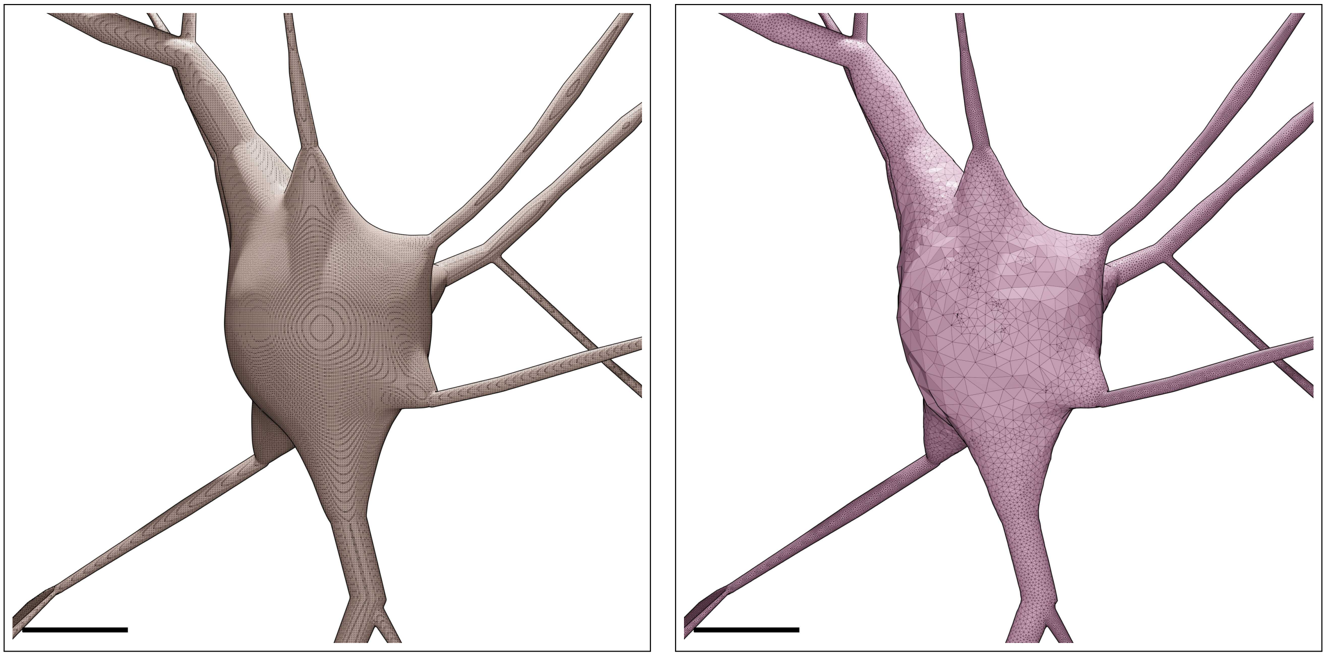

Figure S55: Wireframe visualizations of an L4_LBC neuron showing closeup comparisons between the highly tessellated surface mesh generated from the Voxel remesher (left) and the adaptively optimized surface mesh generated from the optimizer (right). Comparative quantitative and qualitative analyses of the meshes are demonstrated in Figure S56. Scale bars: 5 $\mu$m.

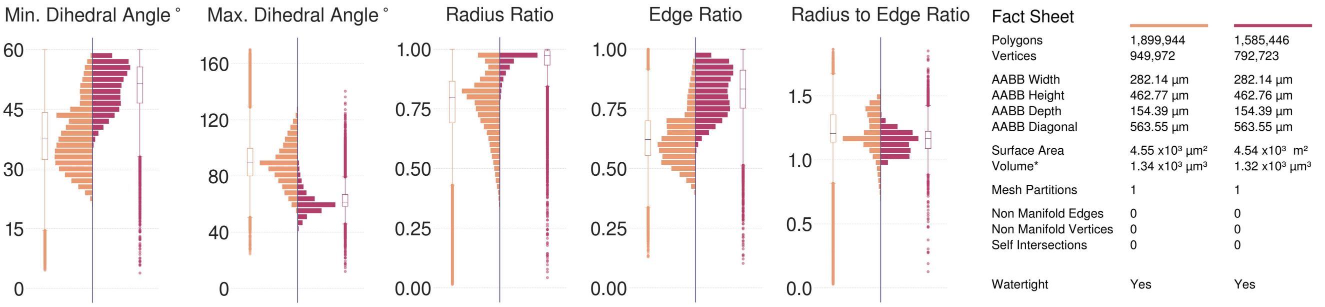

Figure S56: Comparative quantitative and qualitative analyses of the surface mesh models of the L4_LBC neuron visualized in Figure S55.

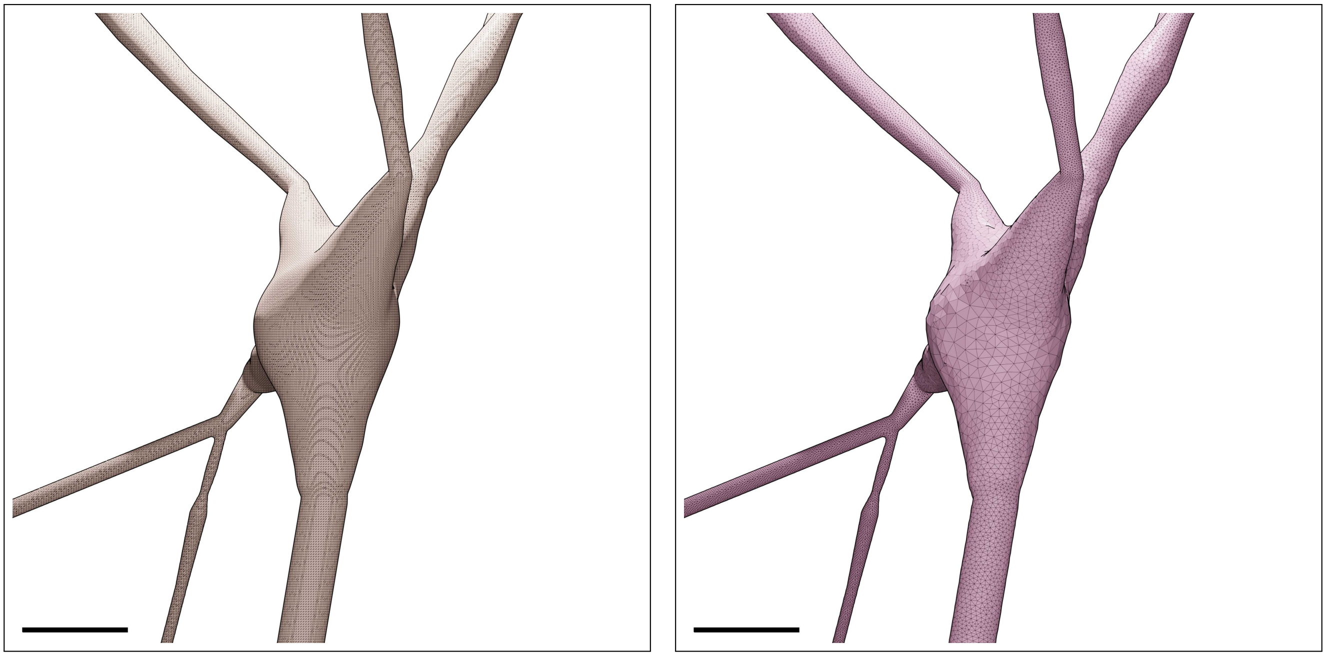

Figure S57: Wireframe visualizations of an L4_MC neuron showing closeup comparisons between the highly tessellated surface mesh generated from the Voxel remesher (left) and the adaptively optimized surface mesh generated from the optimizer (right). Comparative quantitative and qualitative analyses of the meshes are demonstrated in Figure S58. Scale bars: 5 $\mu$m.

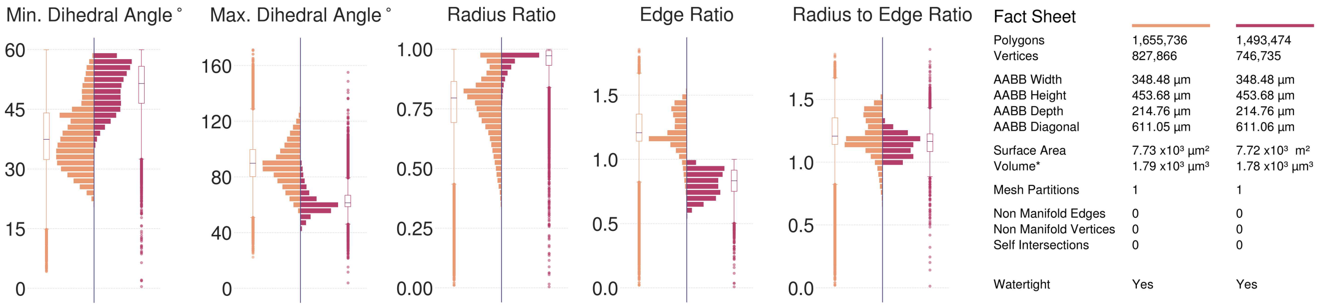

Figure S58: Comparative quantitative and qualitative analyses of the surface mesh models of the L4_MC neuron visualized in Figure S57.

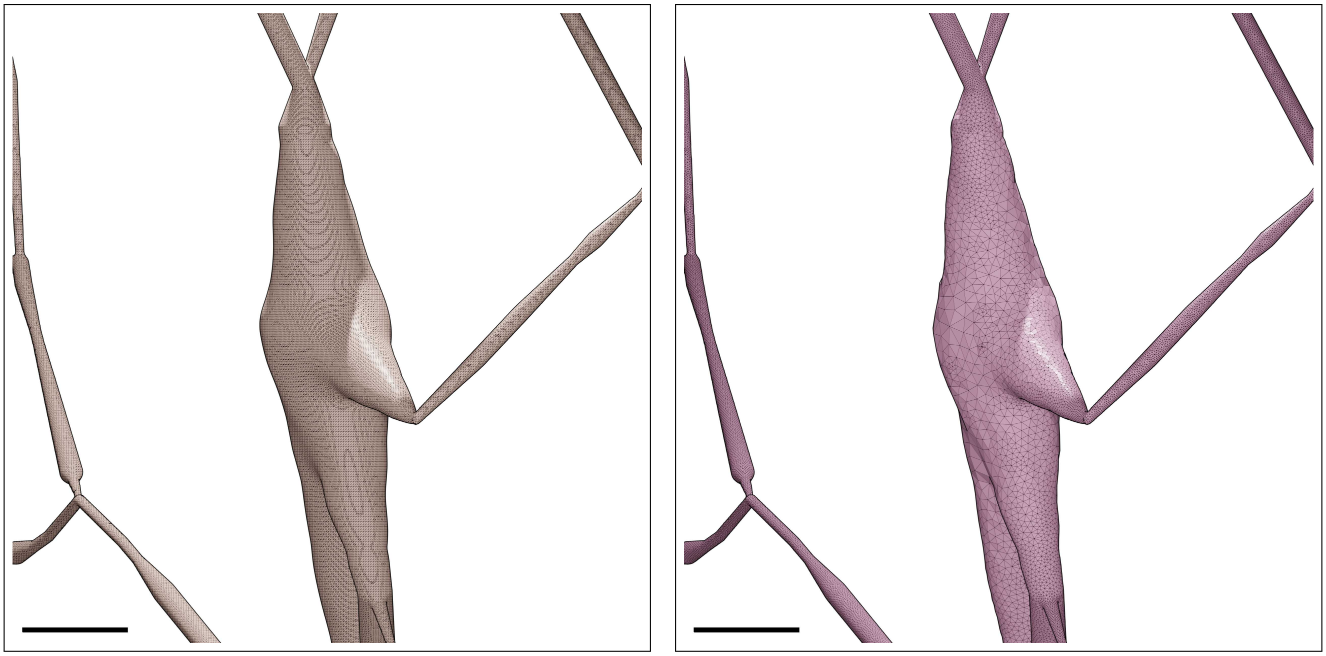

Figure S59: Wireframe visualizations of an L4_NBC neuron showing closeup comparisons between the highly tessellated surface mesh generated from the Voxel remesher (left) and the adaptively optimized surface mesh generated from the optimizer (right). Comparative quantitative and qualitative analyses of the meshes are demonstrated in Figure S60. Scale bars: 5 $\mu$m.

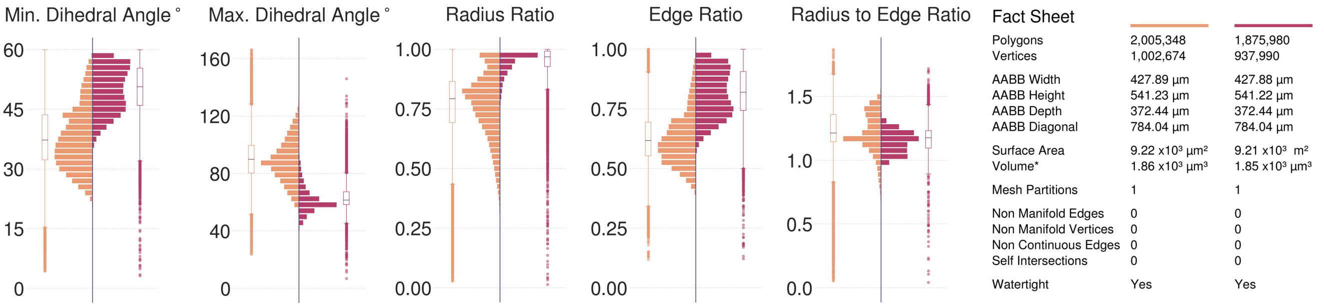

Figure S60: Comparative quantitative and qualitative analyses of the surface mesh models of the L4_NBC neuron visualized in Figure S59.

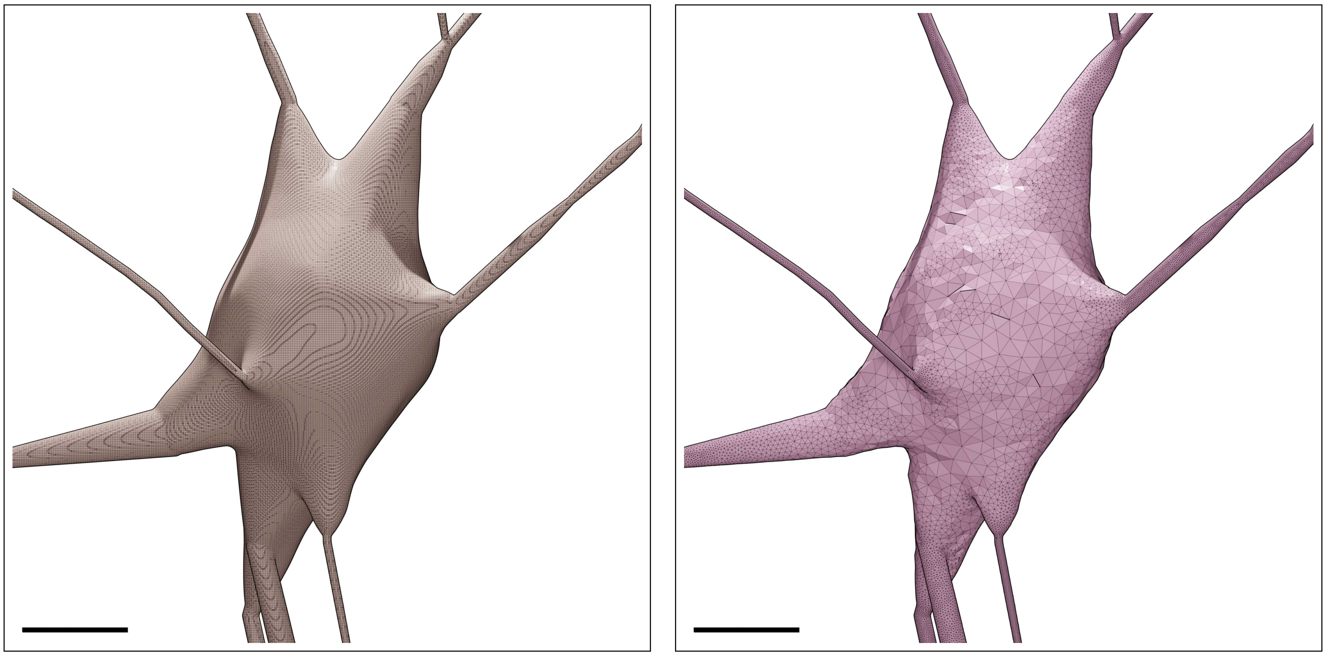

Figure S61: Wireframe visualizations of an L4_NGC neuron showing closeup comparisons between the highly tessellated surface mesh generated from the Voxel remesher (left) and the adaptively optimized surface mesh generated from the optimizer (right). Comparative quantitative and qualitative analyses of the meshes are demonstrated in Figure S62. Scale bars: 5 $\mu$m.

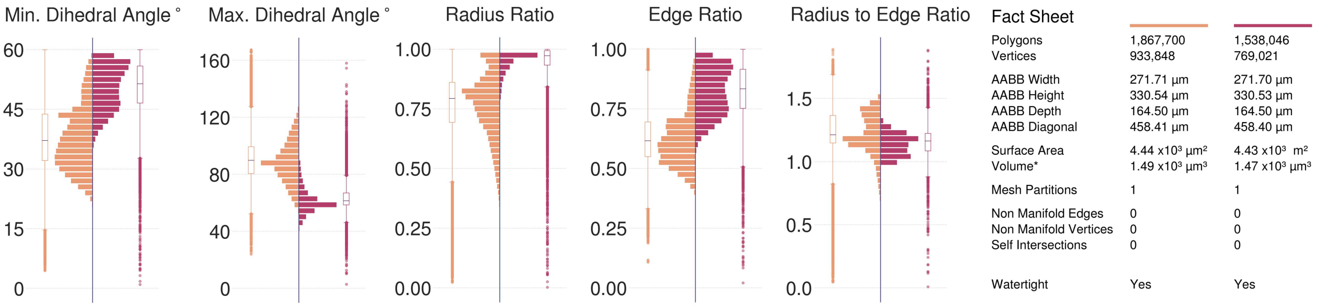

Figure S62: Comparative quantitative and qualitative analyses of the surface mesh models of the L4_NGC neuron visualized in Figure S61.

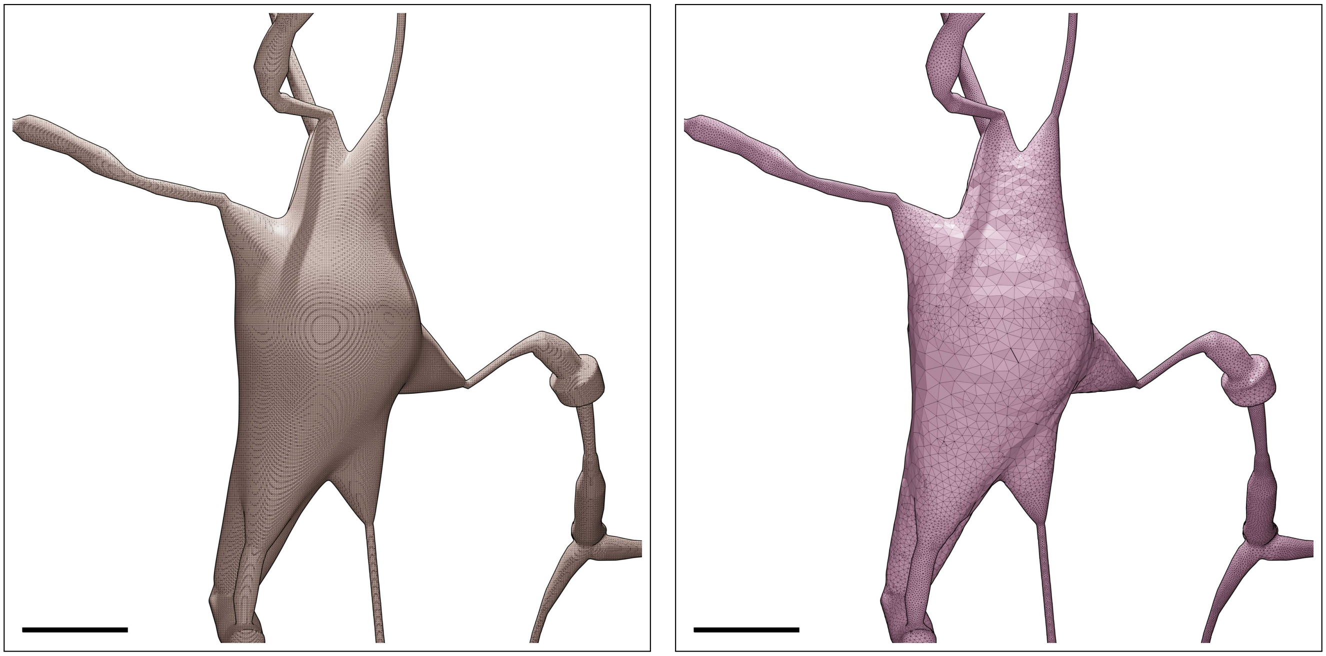

Figure S63: Wireframe visualizations of an L4_SBC neuron showing closeup comparisons between the highly tessellated surface mesh generated from the Voxel remesher (left) and the adaptively optimized surface mesh generated from the optimizer (right). Comparative quantitative and qualitative analyses of the meshes are demonstrated in Figure S64. Scale bars: 5 $\mu$m.

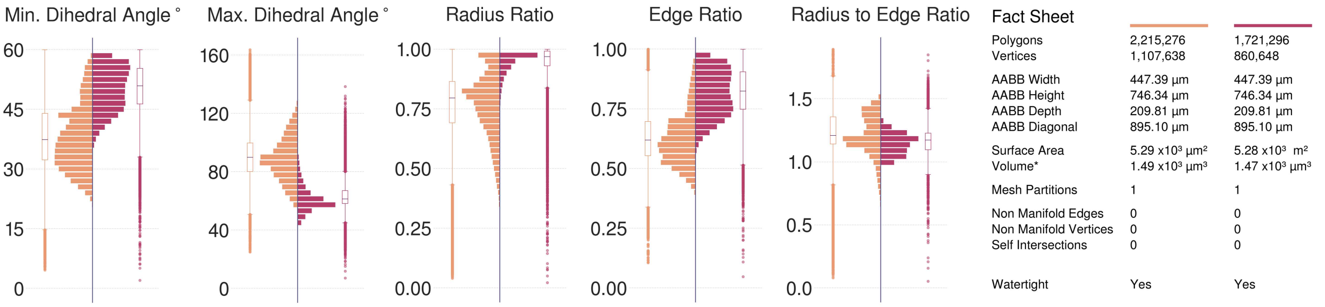

Figure S64: Comparative quantitative and qualitative analyses of the surface mesh models of the L4_SBC neuron visualized in Figure S63.

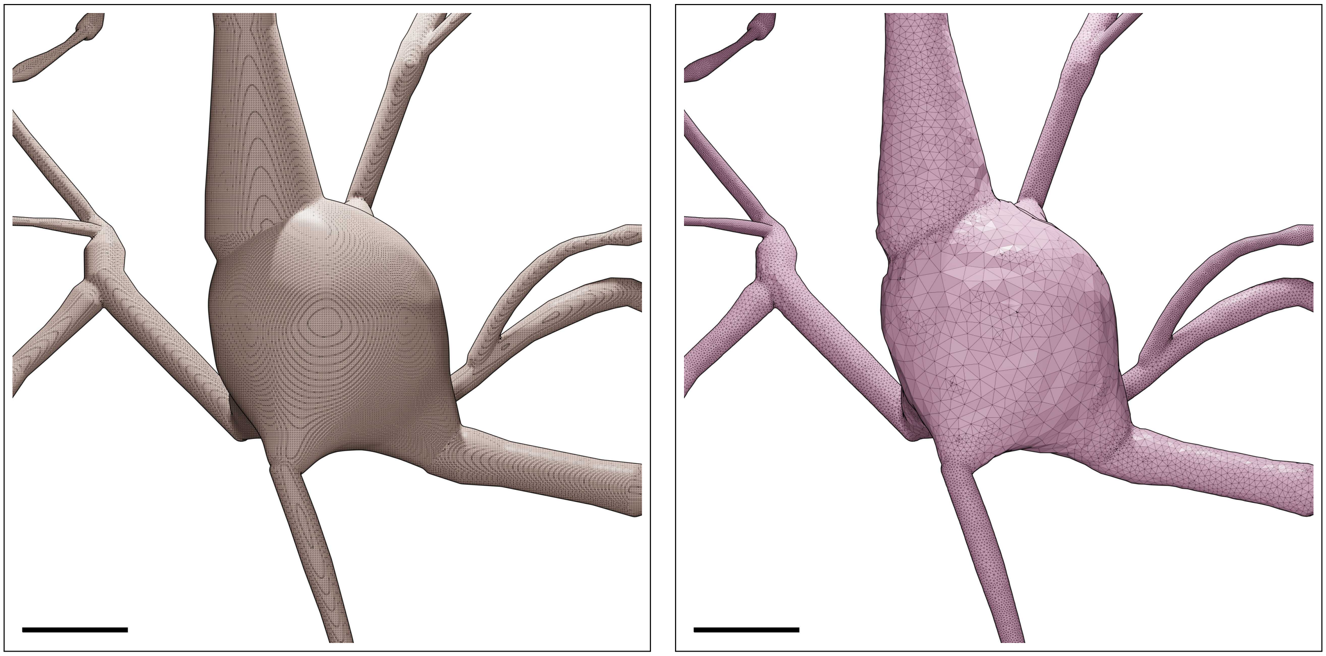

Figure S65: Wireframe visualizations of an L4_SSC neuron showing closeup comparisons between the highly tessellated surface mesh generated from the Voxel remesher (left) and the adaptively optimized surface mesh generated from the optimizer (right). Comparative quantitative and qualitative analyses of the meshes are demonstrated in Figure S66. Scale bars: 5 $\mu$m.

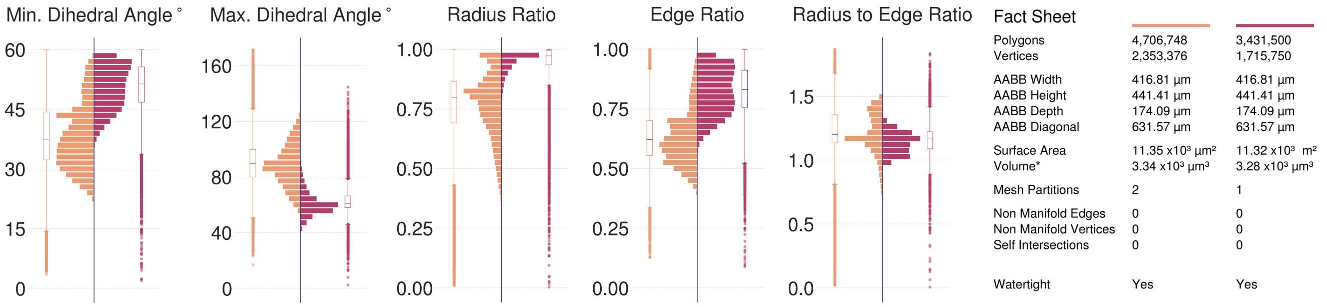

Figure S66: Comparative quantitative and qualitative analyses of the surface mesh models of the L4_SSC neuron visualized in Figure S65.

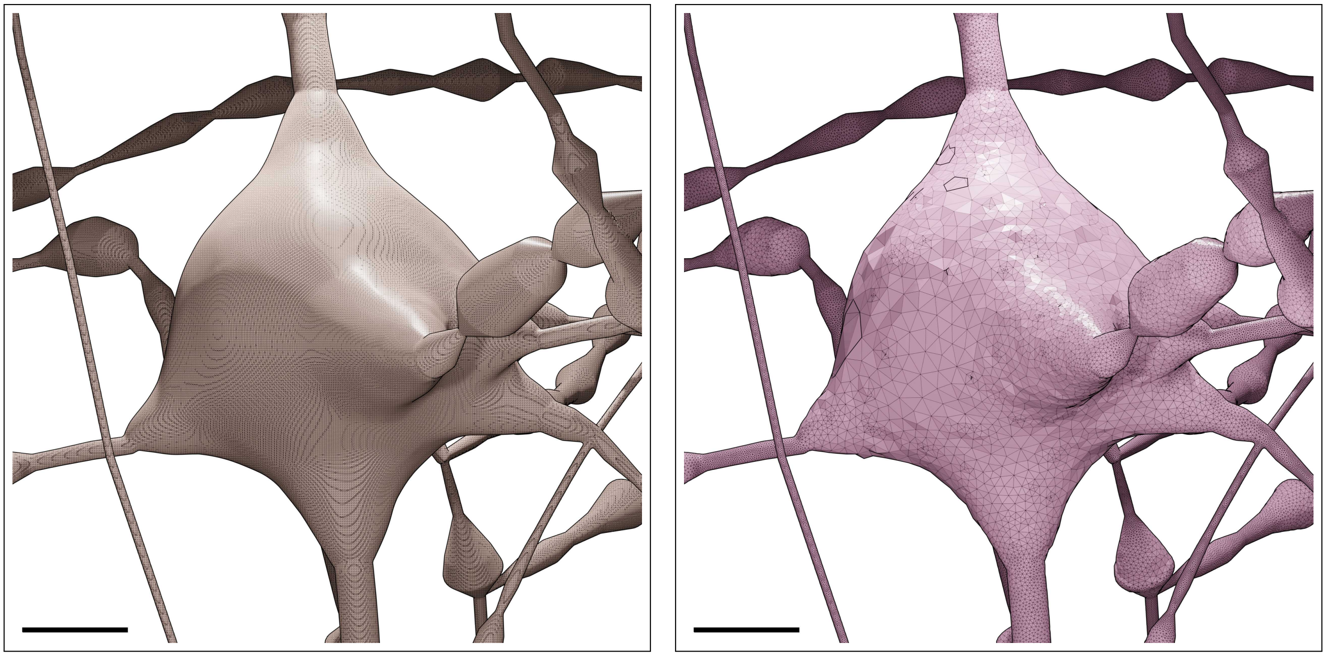

Figure S67: Wireframe visualizations of an L4_TPC neuron showing closeup comparisons between the highly tessellated surface mesh generated from the Voxel remesher (left) and the adaptively optimized surface mesh generated from the optimizer (right). Comparative quantitative and qualitative analyses of the meshes are demonstrated in Figure S68. Scale bars: 5 $\mu$m.

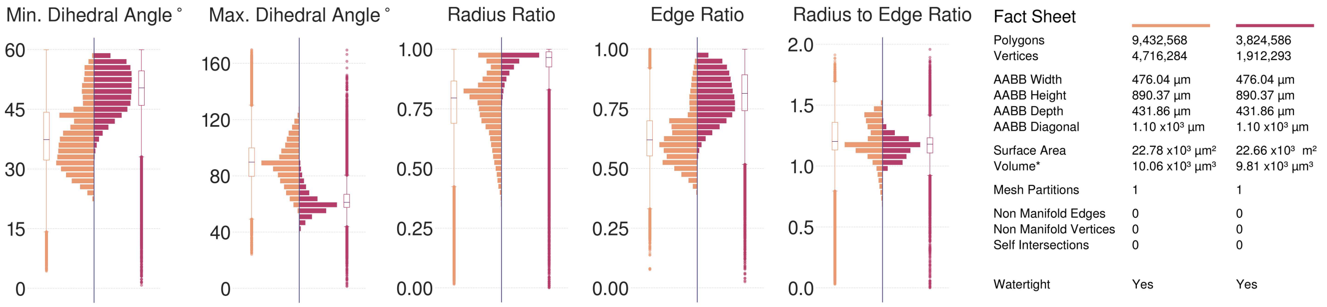

Figure S68: Comparative quantitative and qualitative analyses of the surface mesh models of the L4_TPC neuron visualized in Figure S67.

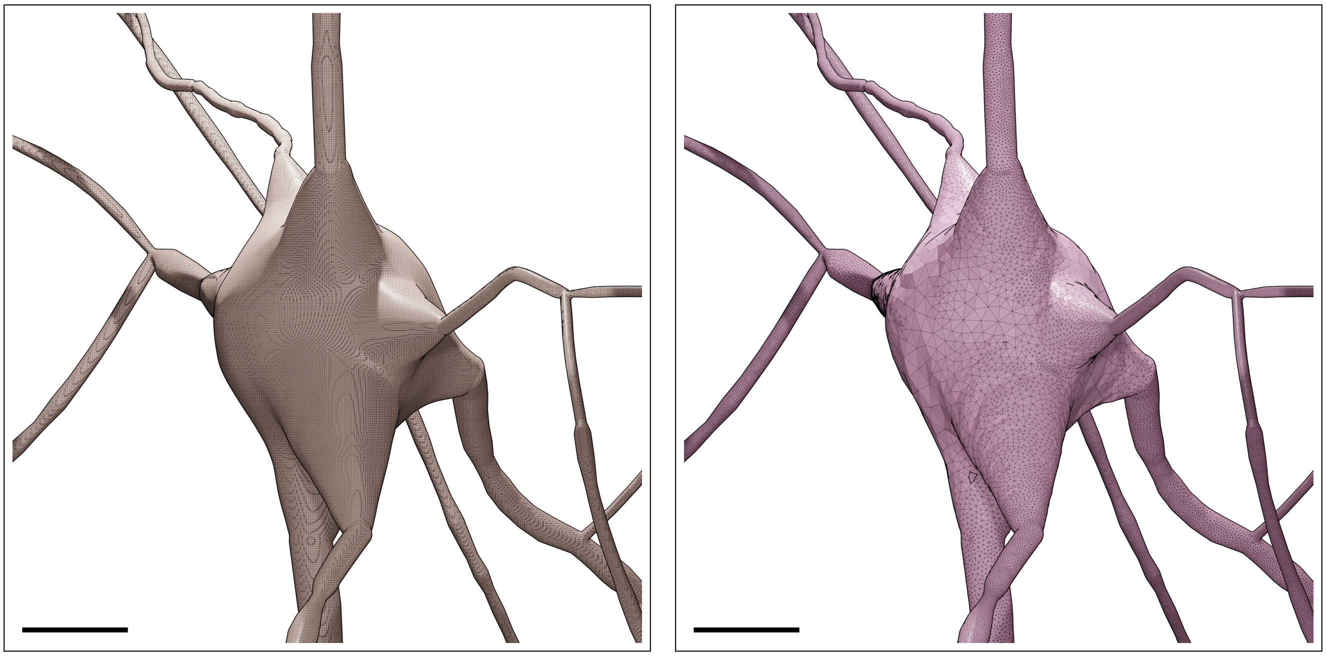

Figure S69: Wireframe visualizations of an L4_UPC neuron showing closeup comparisons between the highly tessellated surface mesh generated from the Voxel remesher (left) and the adaptively optimized surface mesh generated from the optimizer (right). Comparative quantitative and qualitative analyses of the meshes are demonstrated in Figure S70. Scale bars: 5 $\mu$m.

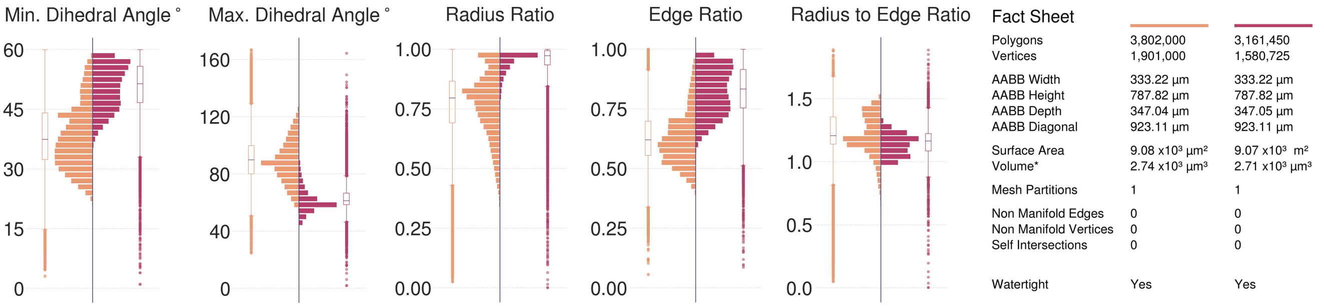

Figure S70: Comparative quantitative and qualitative analyses of the surface mesh models of the L4_UPC neuron visualized in Figure S69.

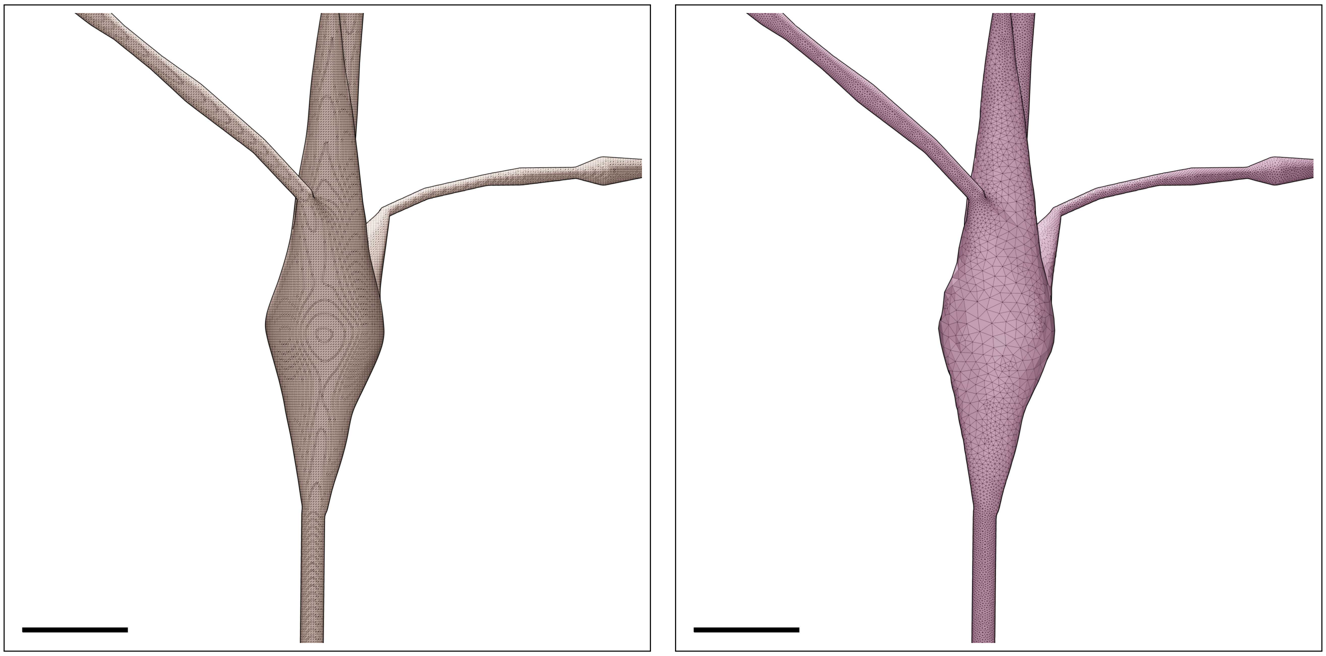

Figure S71: Wireframe visualizations of an L5_BP neuron showing closeup comparisons between the highly tessellated surface mesh generated from the Voxel remesher (left) and the adaptively optimized surface mesh generated from the optimizer (right). Comparative quantitative and qualitative analyses of the meshes are demonstrated in Figure S72. Scale bars: 5 $\mu$m.

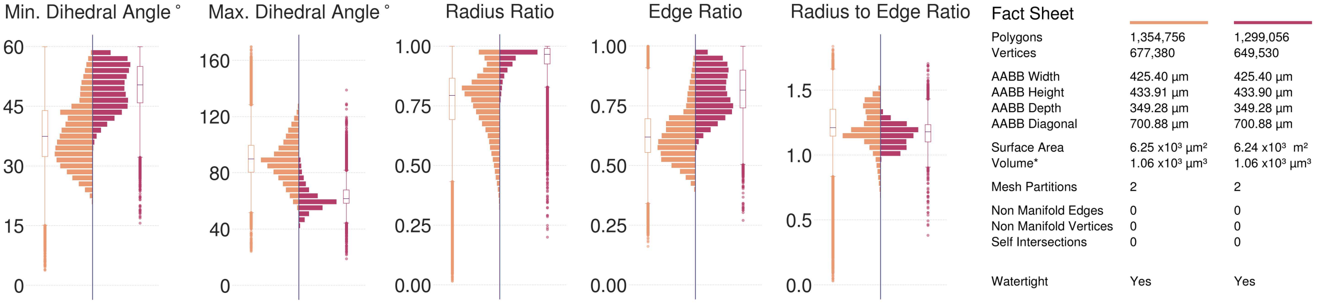

Figure S72: Comparative quantitative and qualitative analyses of the surface mesh models of the L5_BP neuron visualized in Figure S71.

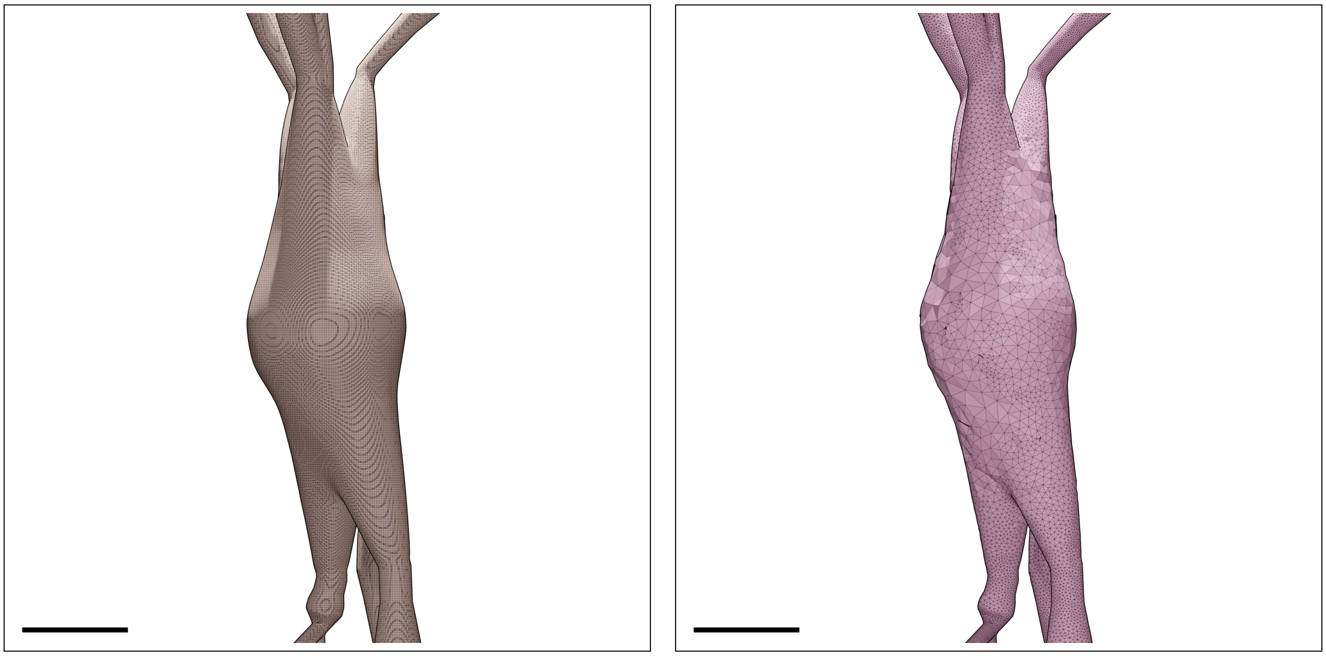

Figure S73: Wireframe visualizations of an L5_BTC neuron showing closeup comparisons between the highly tessellated surface mesh generated from the Voxel remesher (left) and the adaptively optimized surface mesh generated from the optimizer (right). Comparative quantitative and qualitative analyses of the meshes are demonstrated in Figure S74. Scale bars: 5 $\mu$m.

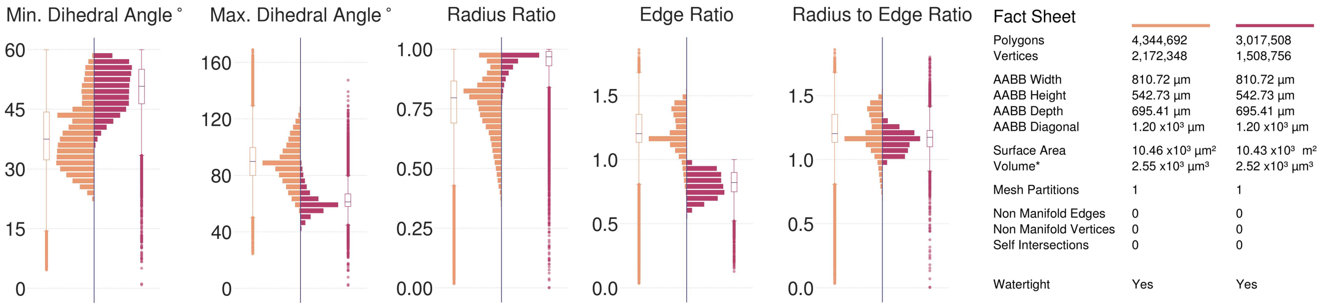

Figure S74: Comparative quantitative and qualitative analyses of the surface mesh models of the L5_BTC neuron visualized in Figure S73.

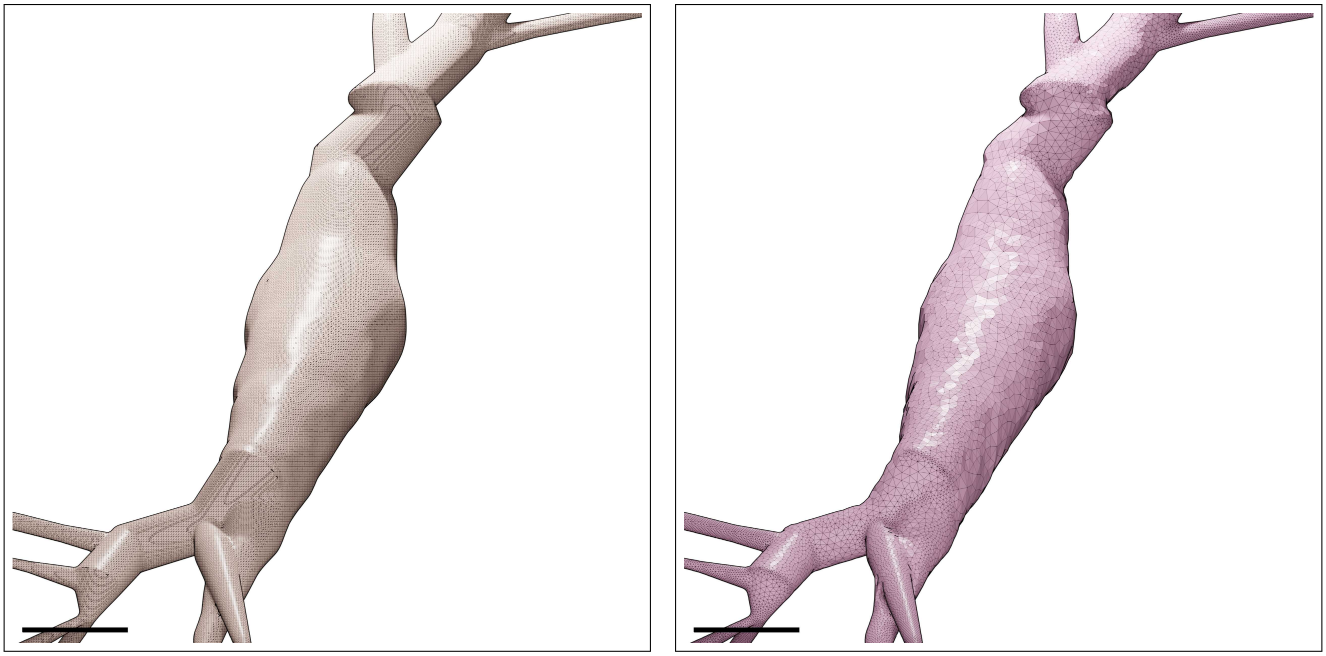

Figure S75: Wireframe visualizations of an L5_CHC neuron showing closeup comparisons between the highly tessellated surface mesh generated from the Voxel remesher (left) and the adaptively optimized surface mesh generated from the optimizer (right). Comparative quantitative and qualitative analyses of the meshes are demonstrated in Figure S76. Scale bars: 5 $\mu$m.

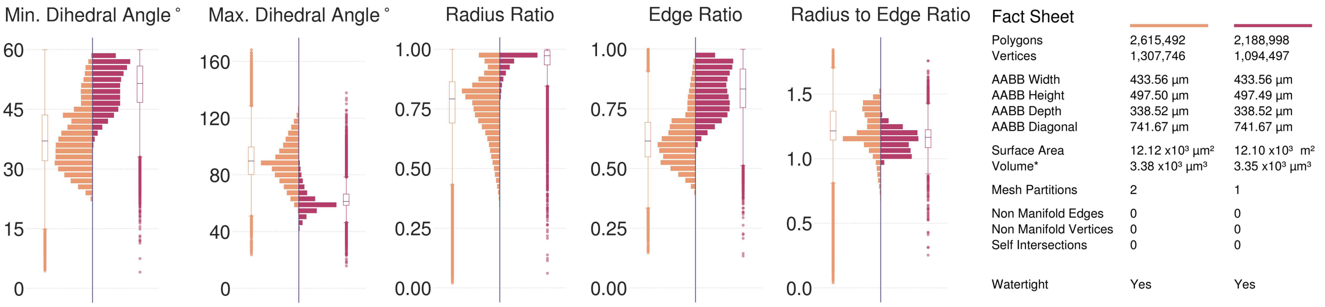

Figure S76: Comparative quantitative and qualitative analyses of the surface mesh models of the L5_CHC neuron visualized in Figure S75.

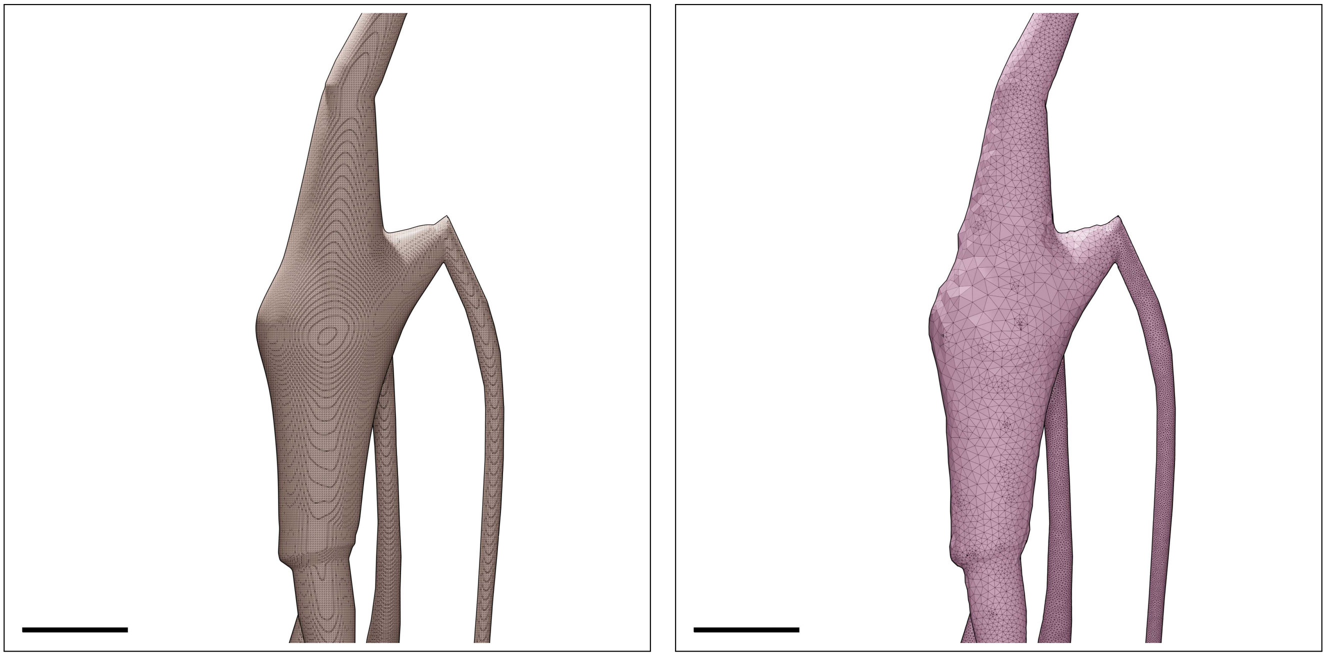

Figure S77: Wireframe visualizations of an L5_DBC neuron showing closeup comparisons between the highly tessellated surface mesh generated from the Voxel remesher (left) and the adaptively optimized surface mesh generated from the optimizer (right). Comparative quantitative and qualitative analyses of the meshes are demonstrated in Figure S78. Scale bars: 5 $\mu$m.

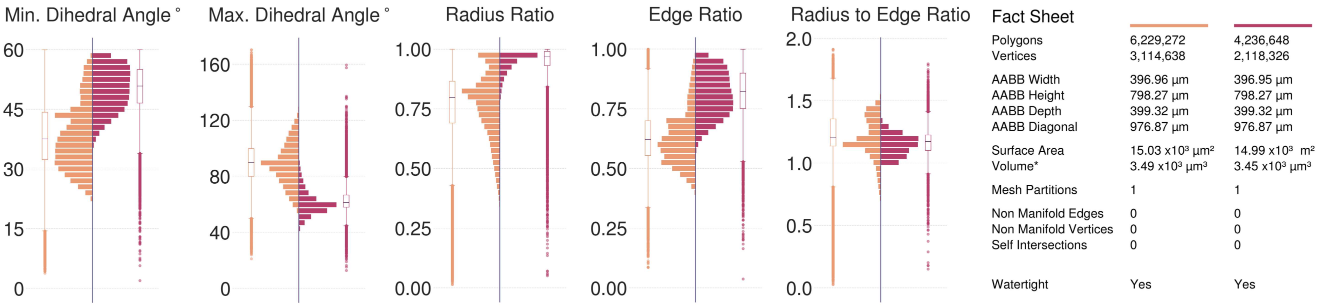

Figure S78: Comparative quantitative and qualitative analyses of the surface mesh models of the L5_DBC neuron visualized in Figure S77.

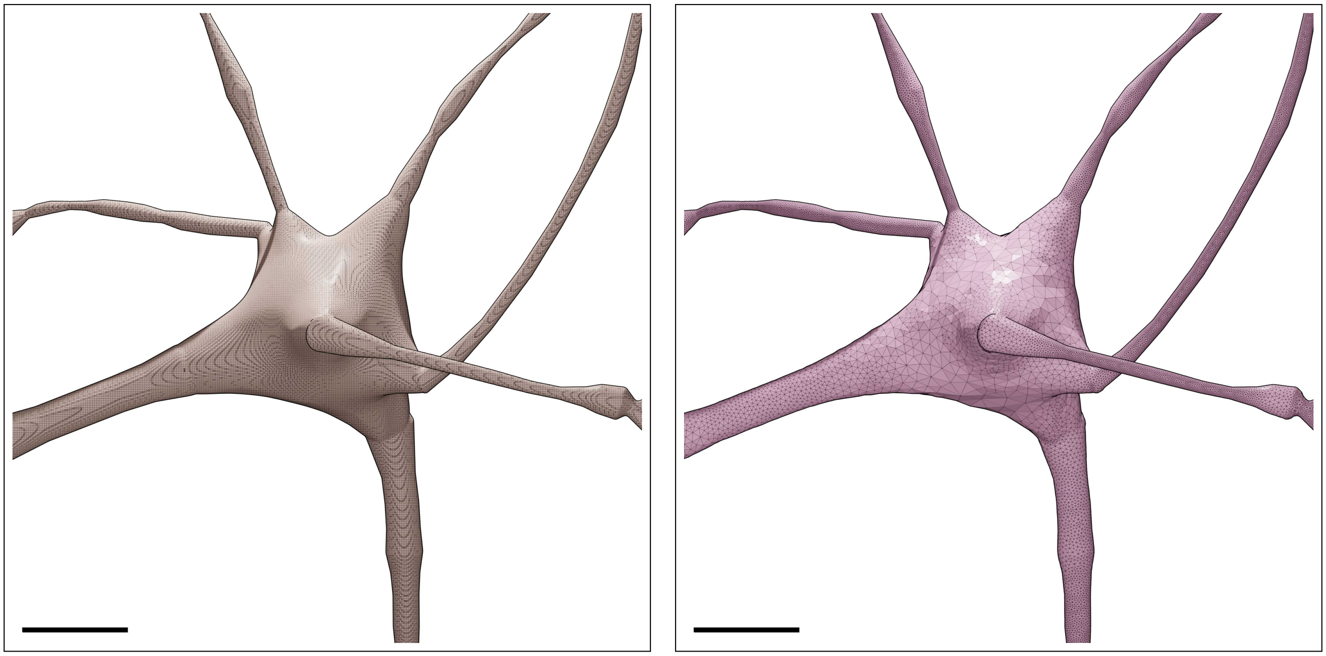

Figure S79: Wireframe visualizations of an L5_LBC neuron showing closeup comparisons between the highly tessellated surface mesh generated from the Voxel remesher (left) and the adaptively optimized surface mesh generated from the optimizer (right). Comparative quantitative and qualitative analyses of the meshes are demonstrated in Figure S80. Scale bars: 5 $\mu$m.

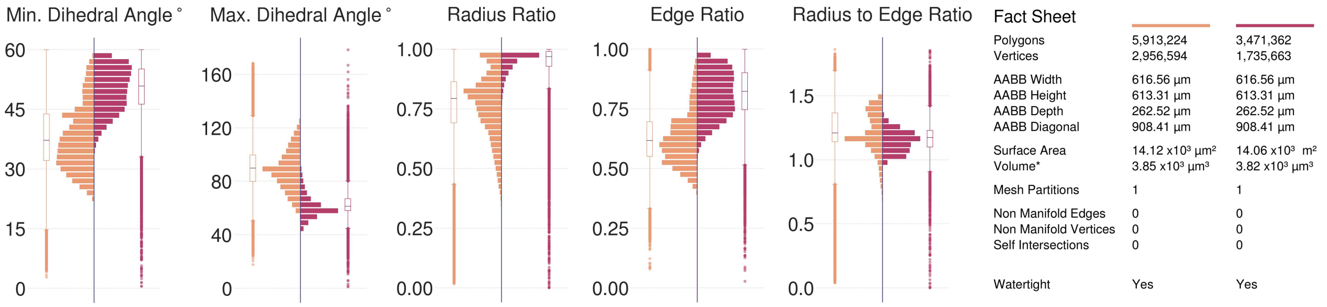

Figure S80: Comparative quantitative and qualitative analyses of the surface mesh models of the L5_LBC neuron visualized in Figure S79.

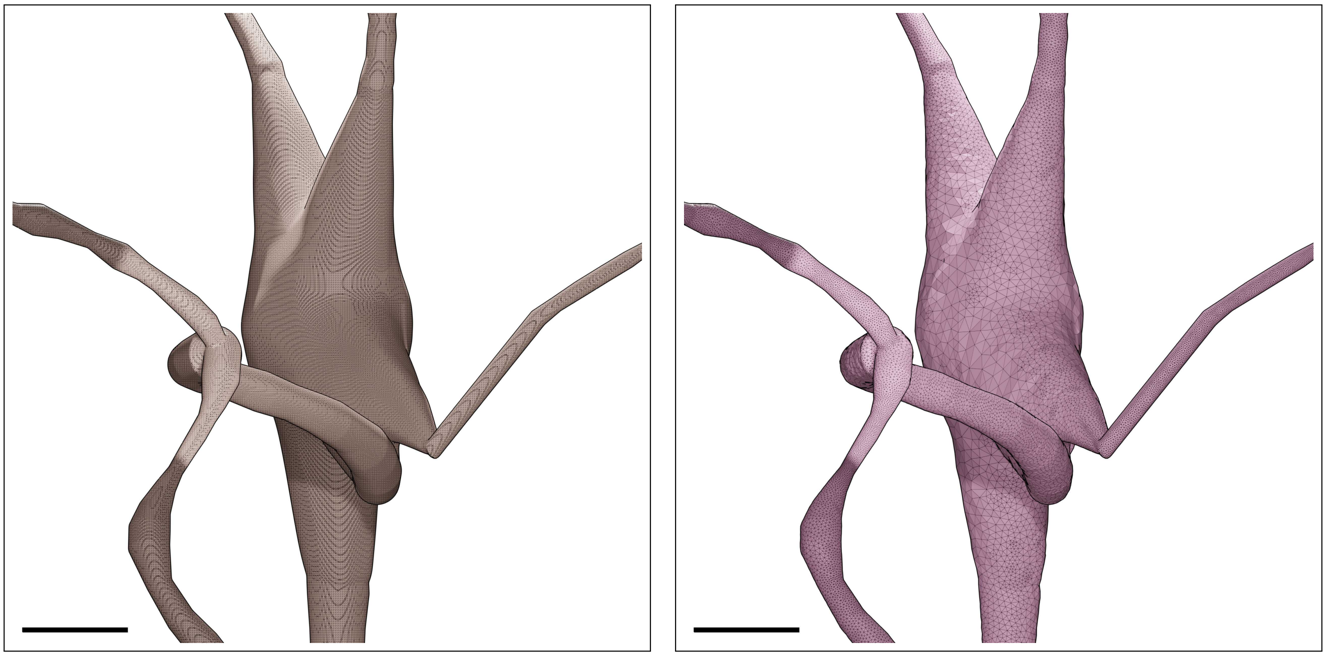

Figure S81: Wireframe visualizations of an L5_MC neuron showing closeup comparisons between the highly tessellated surface mesh generated from the Voxel remesher (left) and the adaptively optimized surface mesh generated from the optimizer (right). Comparative quantitative and qualitative analyses of the meshes are demonstrated in Figure S82. Scale bars: 5 $\mu$m.

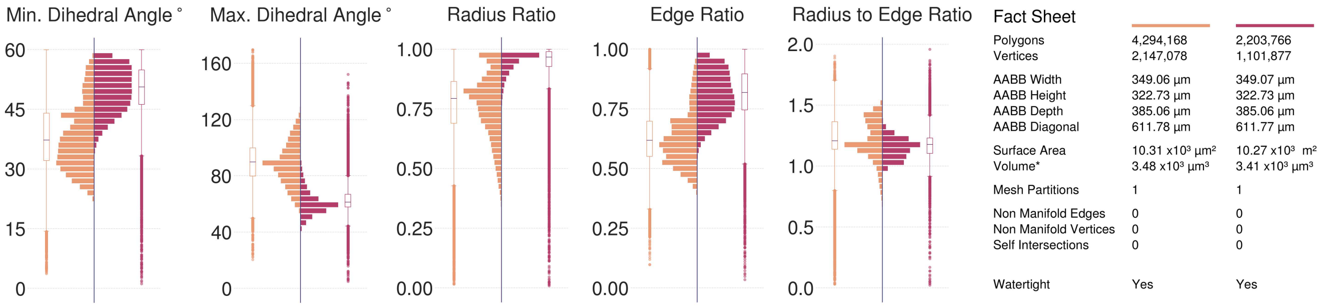

Figure S82: Comparative quantitative and qualitative analyses of the surface mesh models of the L5_MC neuron visualized in Figure S81.

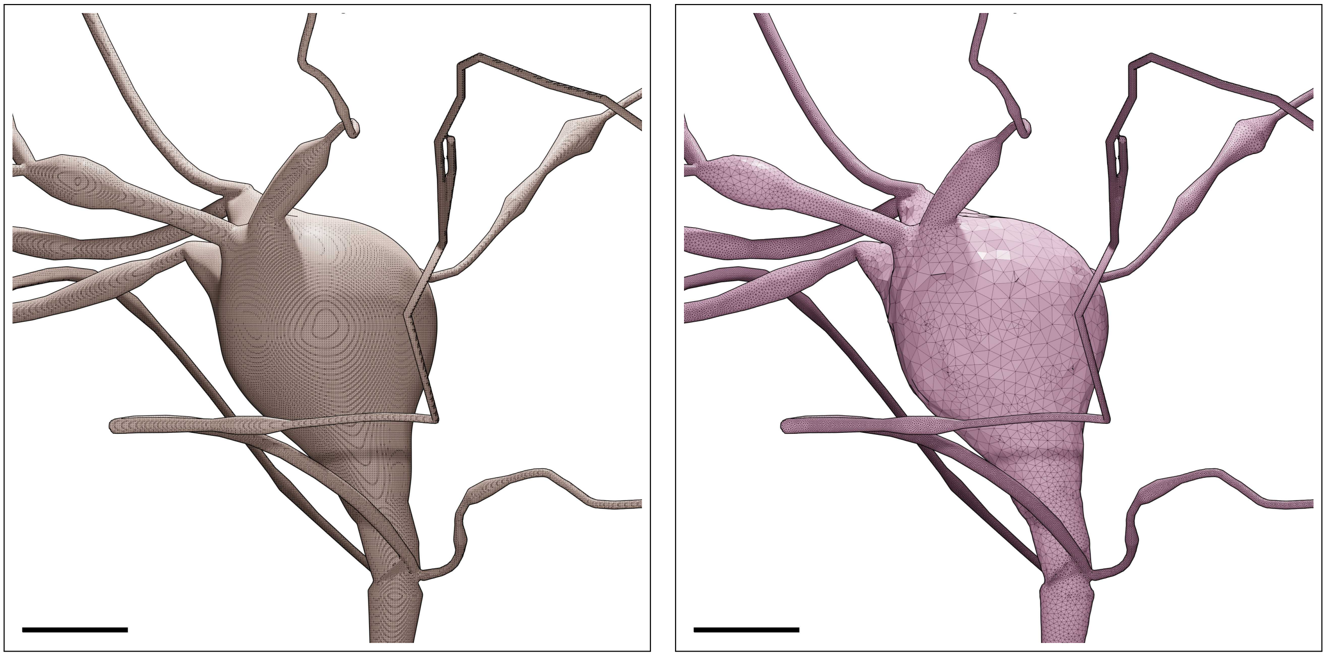

Figure S83: Wireframe visualizations of an L5_NBC neuron showing closeup comparisons between the highly tessellated surface mesh generated from the Voxel remesher (left) and the adaptively optimized surface mesh generated from the optimizer (right). Comparative quantitative and qualitative analyses of the meshes are demonstrated in Figure S84. Scale bars: 5 $\mu$m.

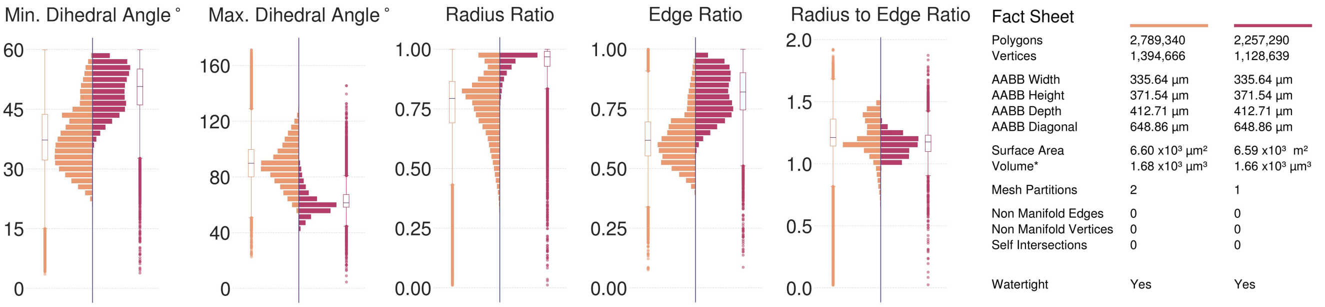

Figure S84: Comparative quantitative and qualitative analyses of the surface mesh models of the L5_NBC neuron visualized in Figure S83.

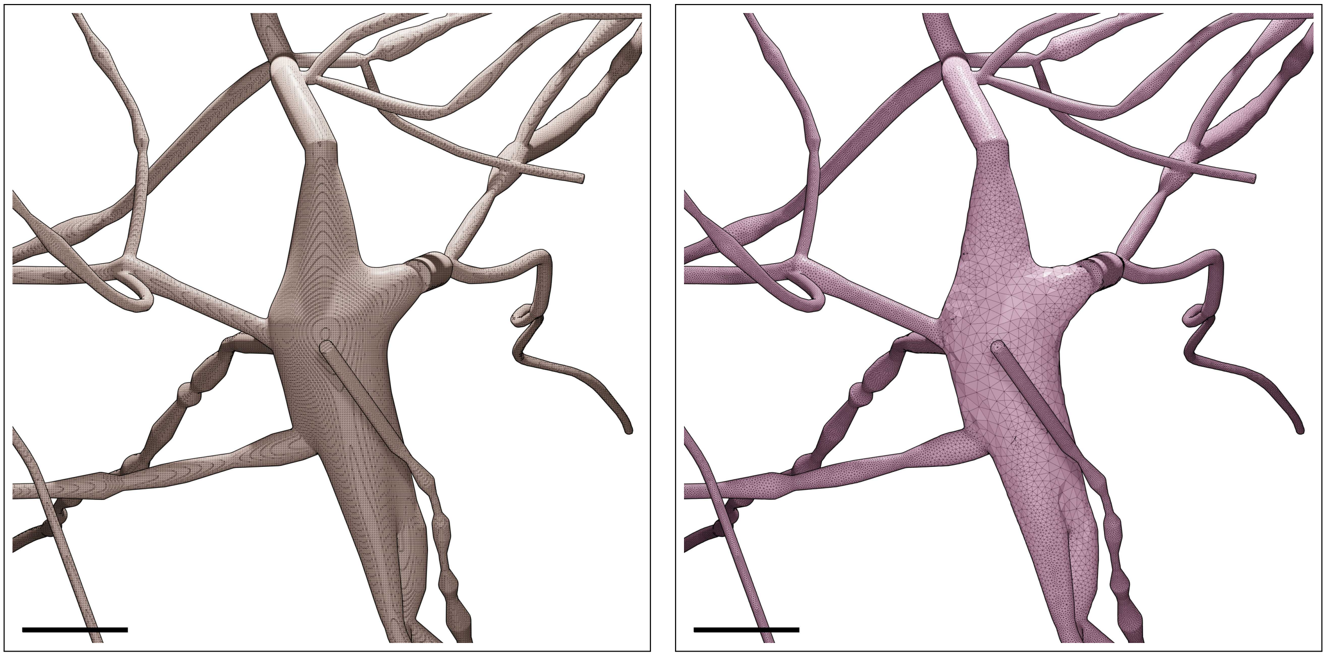

Figure S85: Wireframe visualizations of an L5_NGC neuron showing closeup comparisons between the highly tessellated surface mesh generated from the Voxel remesher (left) and the adaptively optimized surface mesh generated from the optimizer (right). Comparative quantitative and qualitative analyses of the meshes are demonstrated in Figure S86. Scale bars: 5 $\mu$m.

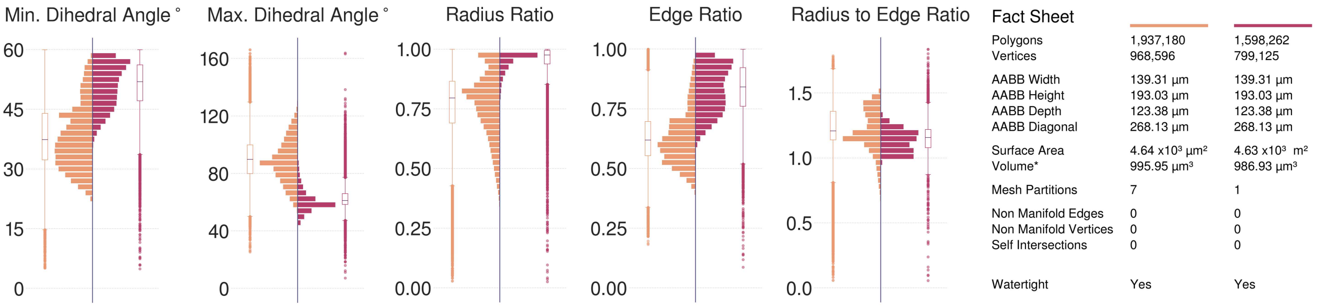

Figure S86: Comparative quantitative and qualitative analyses of the surface mesh models of the L5_NGC neuron visualized in Figure S85.

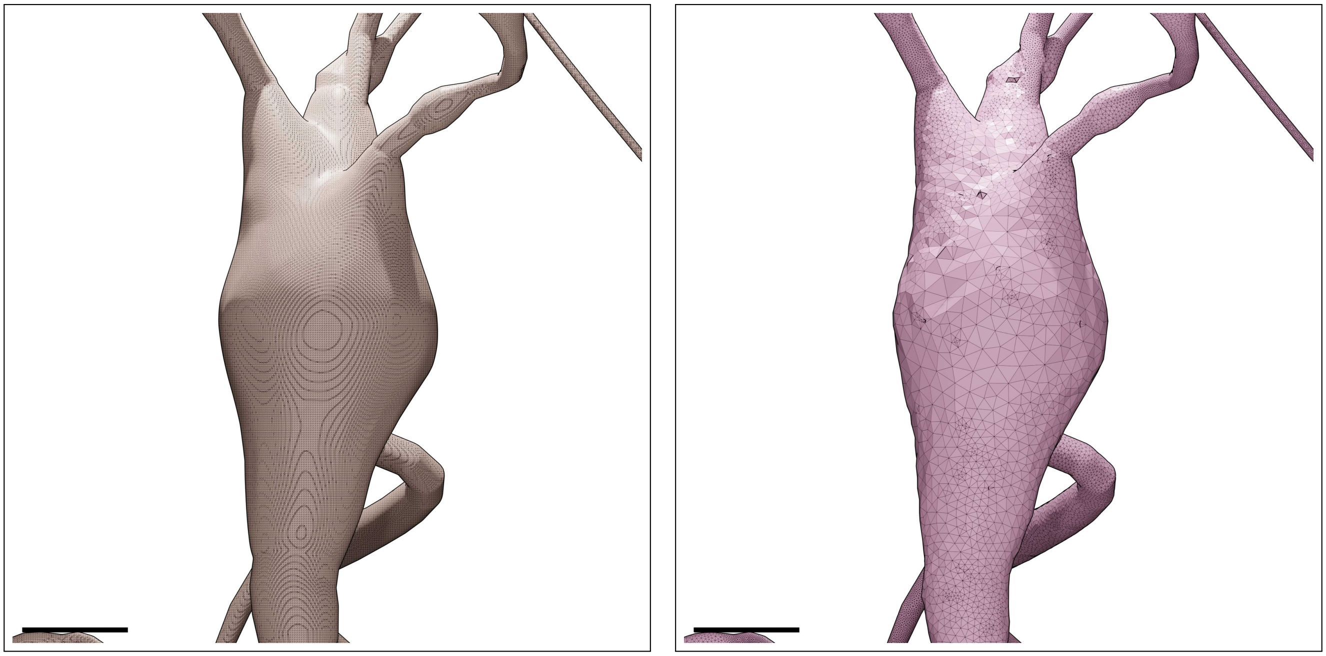

Figure S87: Wireframe visualizations of an L5_SBC neuron showing closeup comparisons between the highly tessellated surface mesh generated from the Voxel remesher (left) and the adaptively optimized surface mesh generated from the optimizer (right). Comparative quantitative and qualitative analyses of the meshes are demonstrated in Figure S88. Scale bars: 5 $\mu$m.

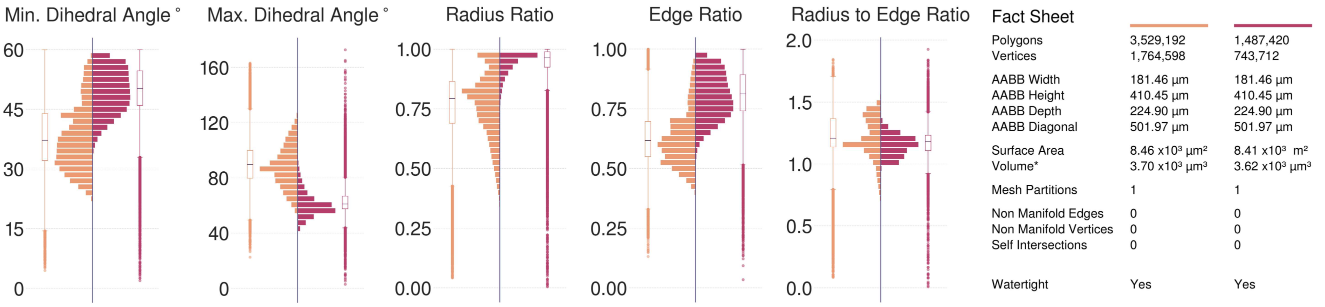

Figure S88: Comparative quantitative and qualitative analyses of the surface mesh models of the L5_SBC neuron visualized in Figure S87.

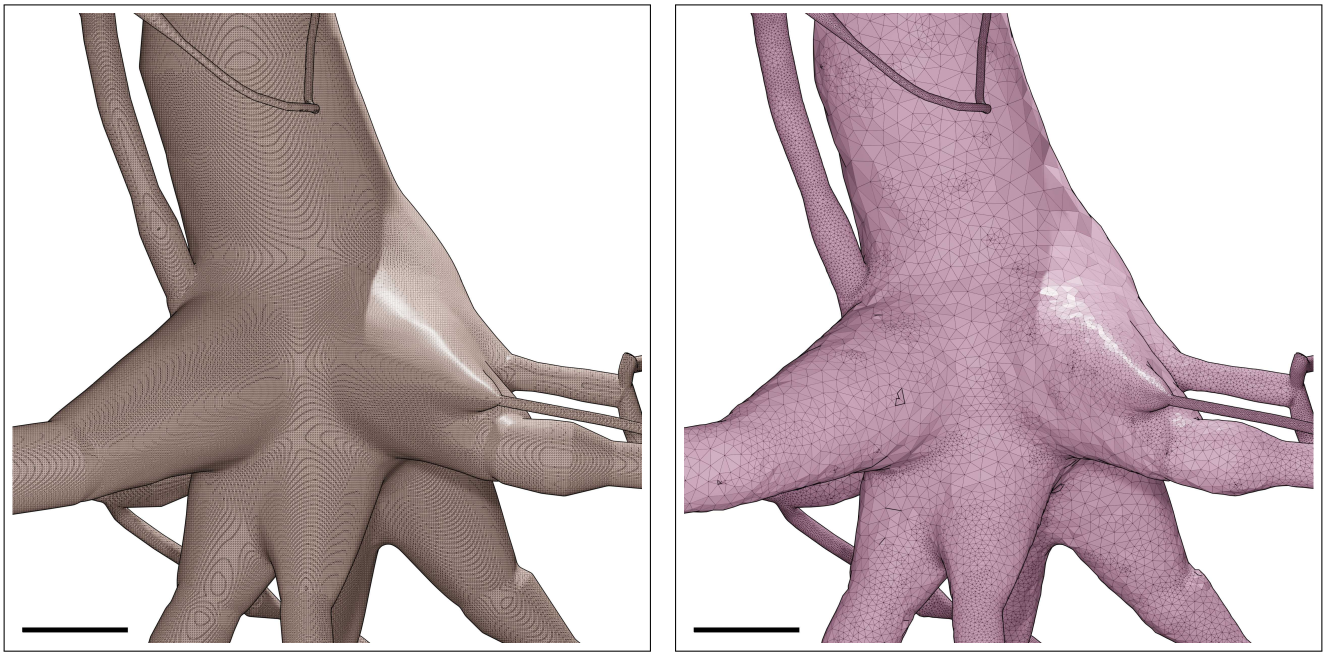

Figure S89: Wireframe visualizations of an L5_TPC:A neuron showing closeup comparisons between the highly tessellated surface mesh generated from the Voxel remesher (left) and the adaptively optimized surface mesh generated from the optimizer (right). Comparative quantitative and qualitative analyses of the meshes are demonstrated in Figure S90. Scale bars: 5 $\mu$m.

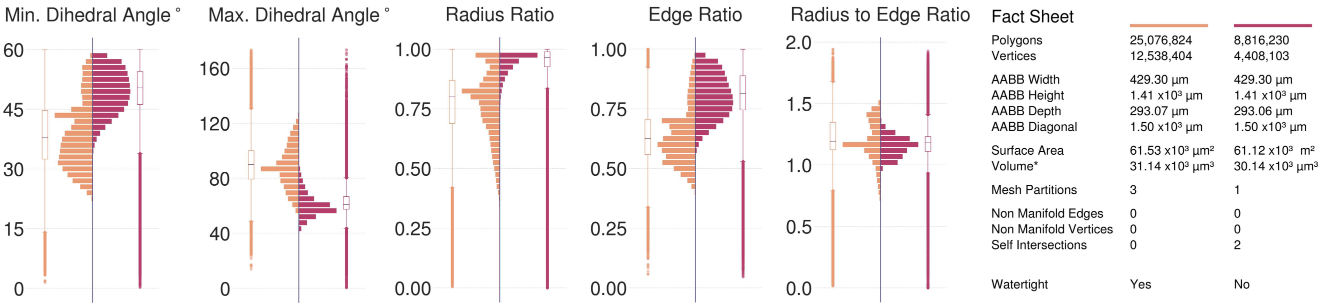

Figure S90: Comparative quantitative and qualitative analyses of the surface mesh models of the L5_TPC:A neuron visualized in Figure S89.

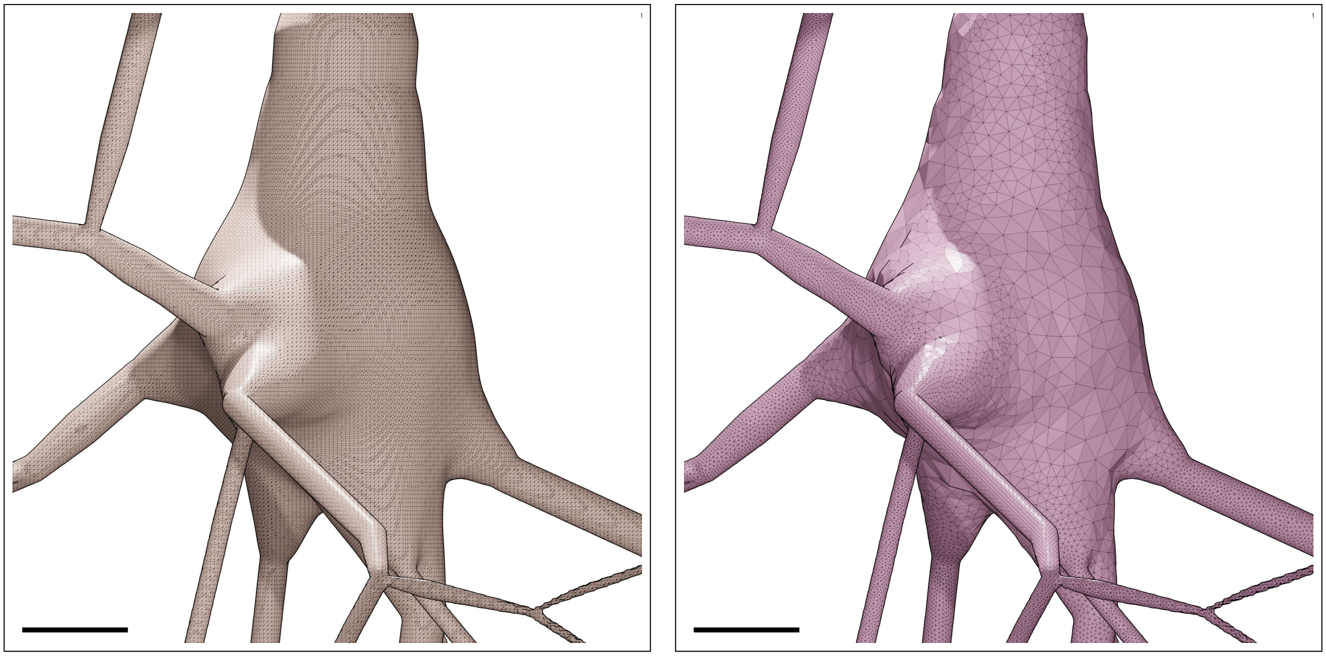

Figure S91: Wireframe visualizations of an L5_TPC:B neuron showing closeup comparisons between the highly tessellated surface mesh generated from the Voxel remesher (left) and the adaptively optimized surface mesh generated from the optimizer (right). Comparative quantitative and qualitative analyses of the meshes are demonstrated in Figure S92. Scale bars: 5 $\mu$m.

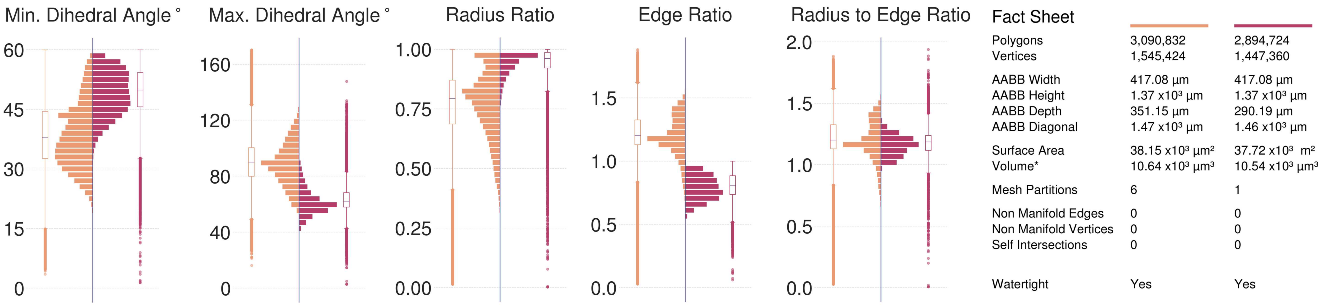

Figure S92: Comparative quantitative and qualitative analyses of the surface mesh models of the L5_TPC:B neuron visualized in Figure S91.

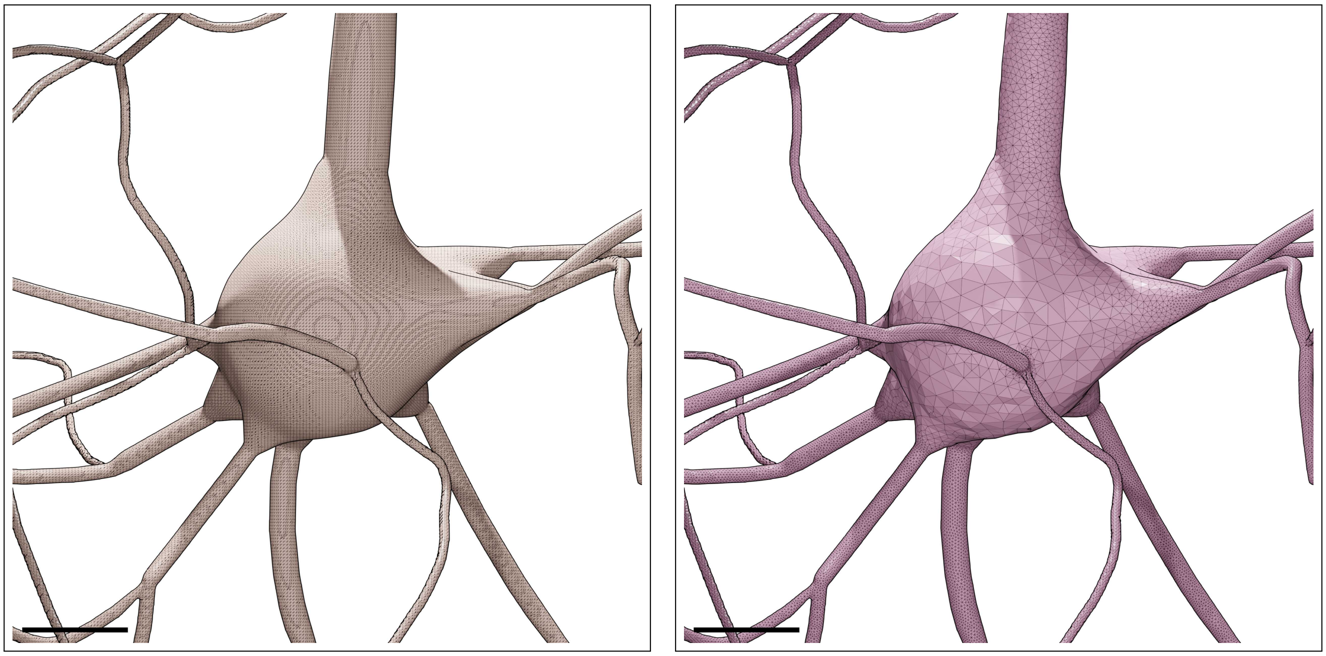

Figure S93: Wireframe visualizations of an L5_TPC:C neuron showing closeup comparisons between the highly tessellated surface mesh generated from the Voxel remesher (left) and the adaptively optimized surface mesh generated from the optimizer (right). Comparative quantitative and qualitative analyses of the meshes are demonstrated in Figure S94. Scale bars: 5 $\mu$m.

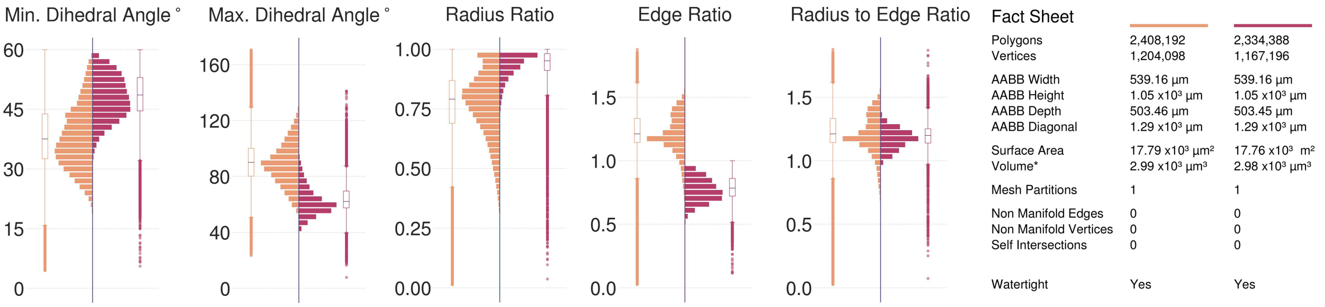

Figure S94: Comparative quantitative and qualitative analyses of the surface mesh models of the L5_TPC:C neuron visualized in Figure S93.

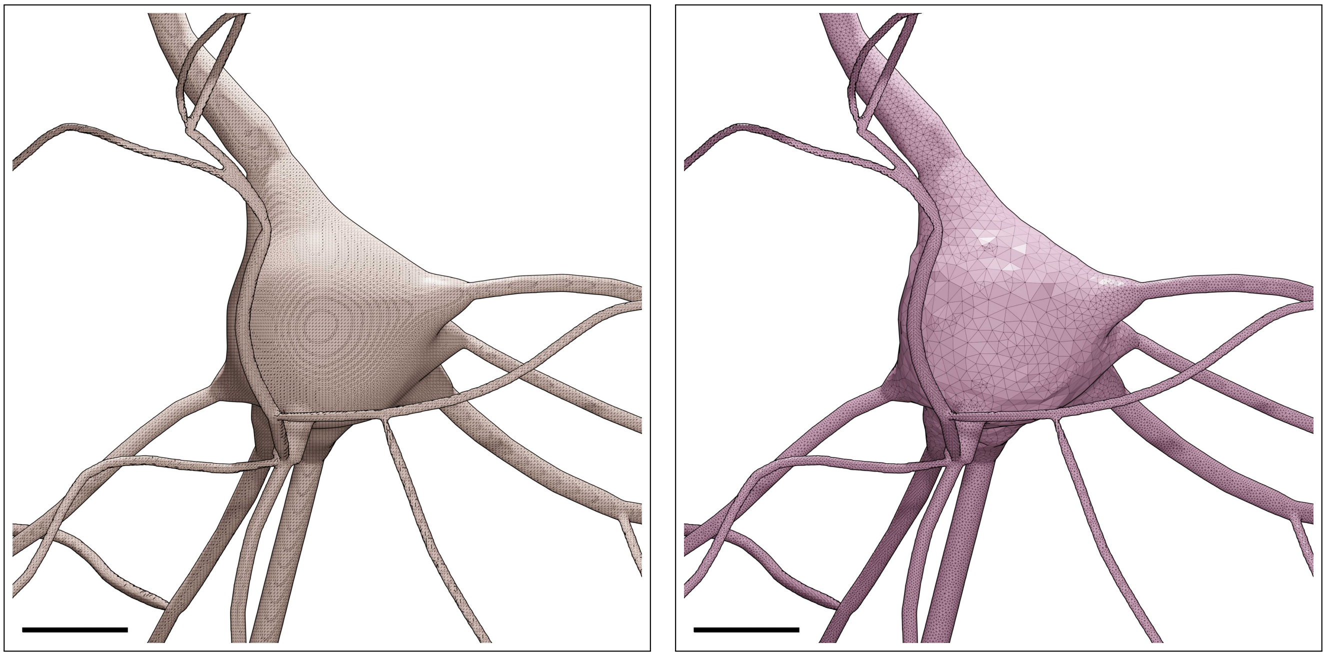

Figure S95: Wireframe visualizations of an L5_UPC neuron showing closeup comparisons between the highly tessellated surface mesh generated from the Voxel remesher (left) and the adaptively optimized surface mesh generated from the optimizer (right). Comparative quantitative and qualitative analyses of the meshes are demonstrated in Figure S96. Scale bars: 5 $\mu$m.

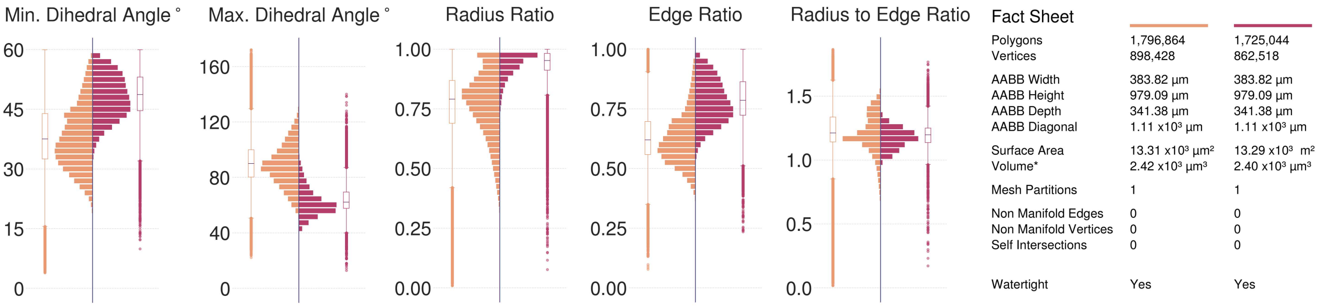

Figure S96: Comparative quantitative and qualitative analyses of the surface mesh models of the L5_UPC neuron visualized in Figure S95.

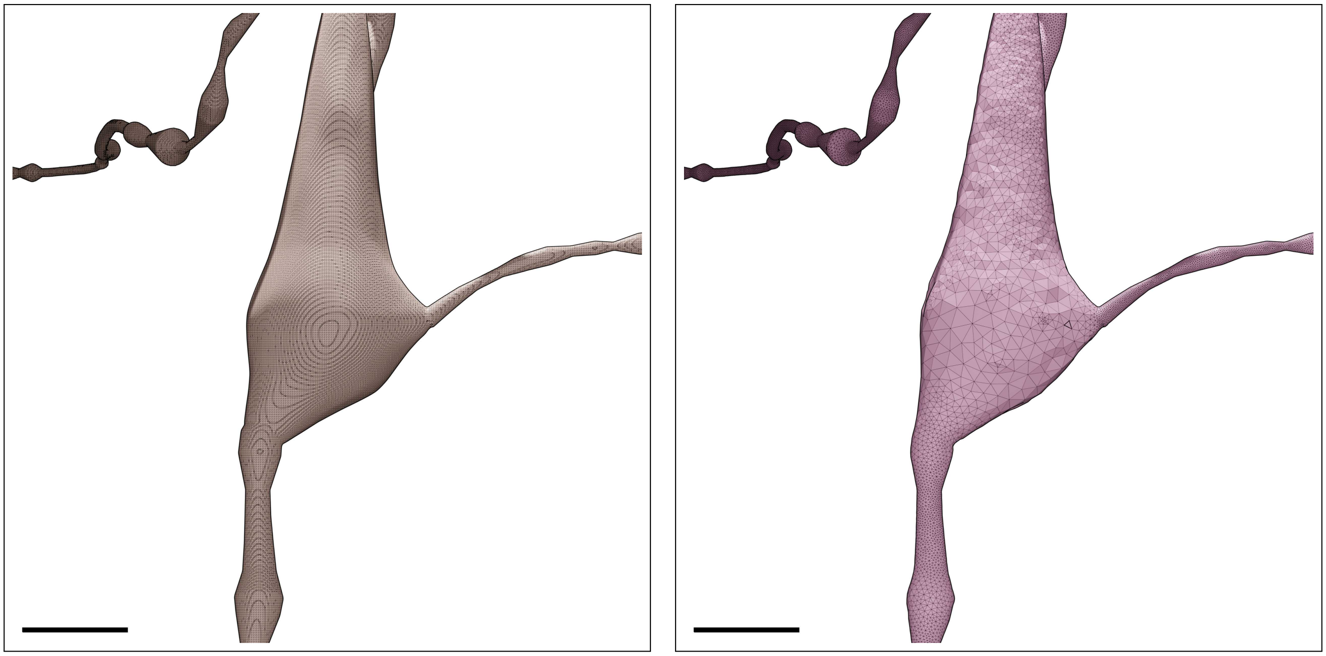

Figure S97: Wireframe visualizations of an L6_BP neuron showing closeup comparisons between the highly tessellated surface mesh generated from the Voxel remesher (left) and the adaptively optimized surface mesh generated from the optimizer (right). Comparative quantitative and qualitative analyses of the meshes are demonstrated in Figure S98. Scale bars: 5 $\mu$m.

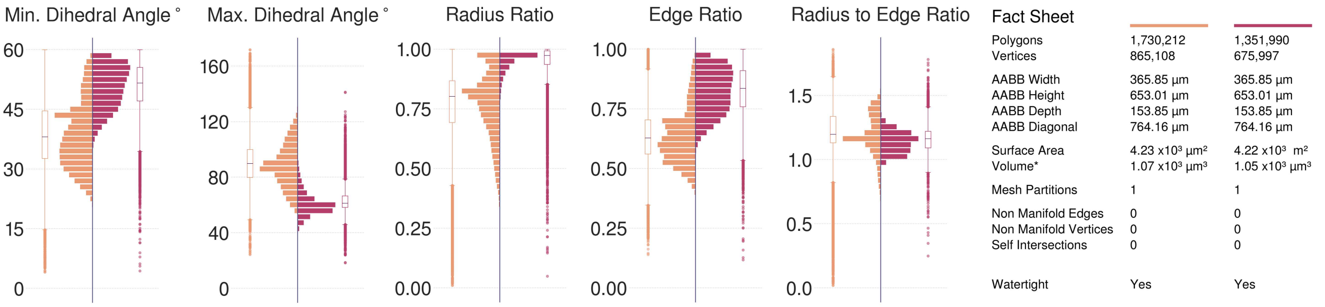

Figure S98: Comparative quantitative and qualitative analyses of the surface mesh models of the L6_BP neuron visualized in Figure S97.

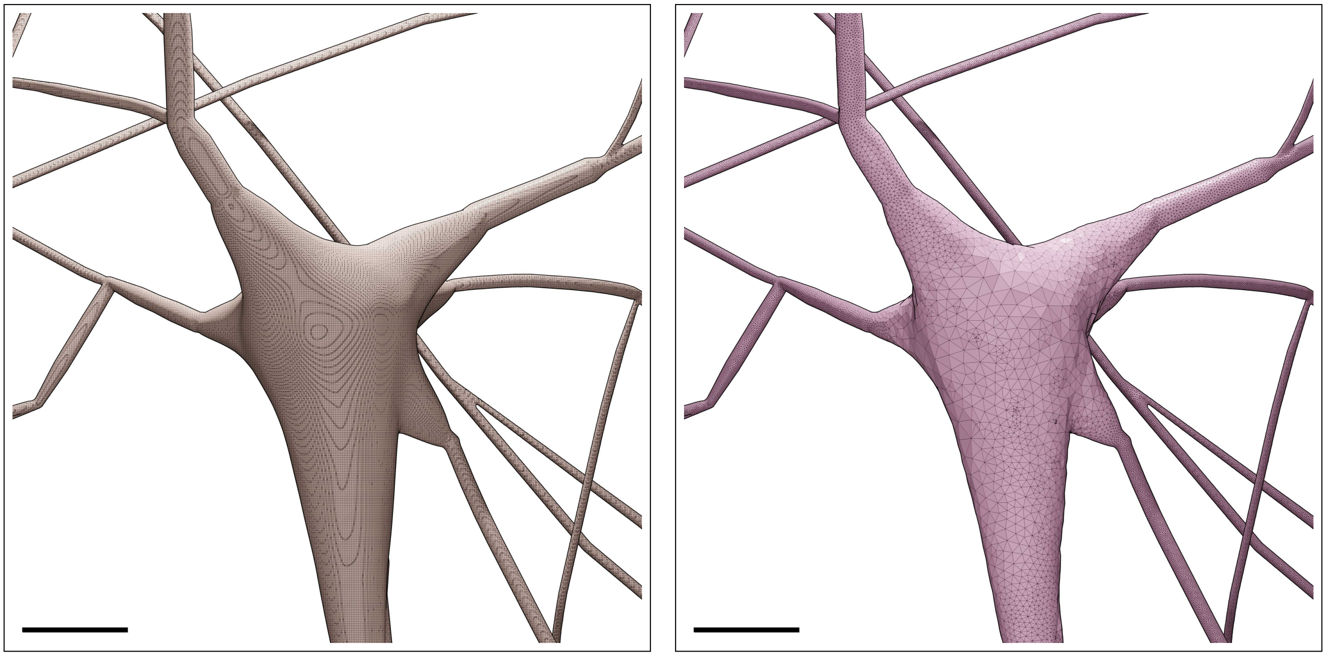

Figure S99: Wireframe visualizations of an L6_BPC neuron showing closeup comparisons between the highly tessellated surface mesh generated from the Voxel remesher (left) and the adaptively optimized surface mesh generated from the optimizer (right). Comparative quantitative and qualitative analyses of the meshes are demonstrated in Figure S100. Scale bars: 5 $\mu$m.

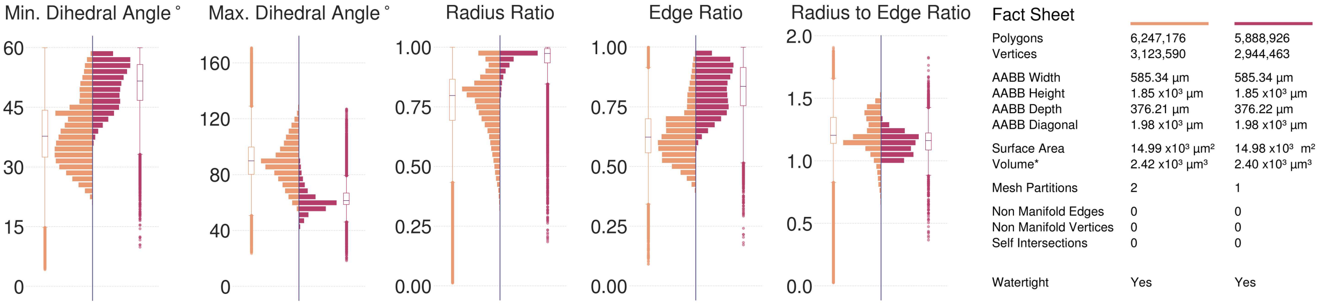

Figure S100: Comparative quantitative and qualitative analyses of the surface mesh models of the L6_BPC neuron visualized in Figure S99.

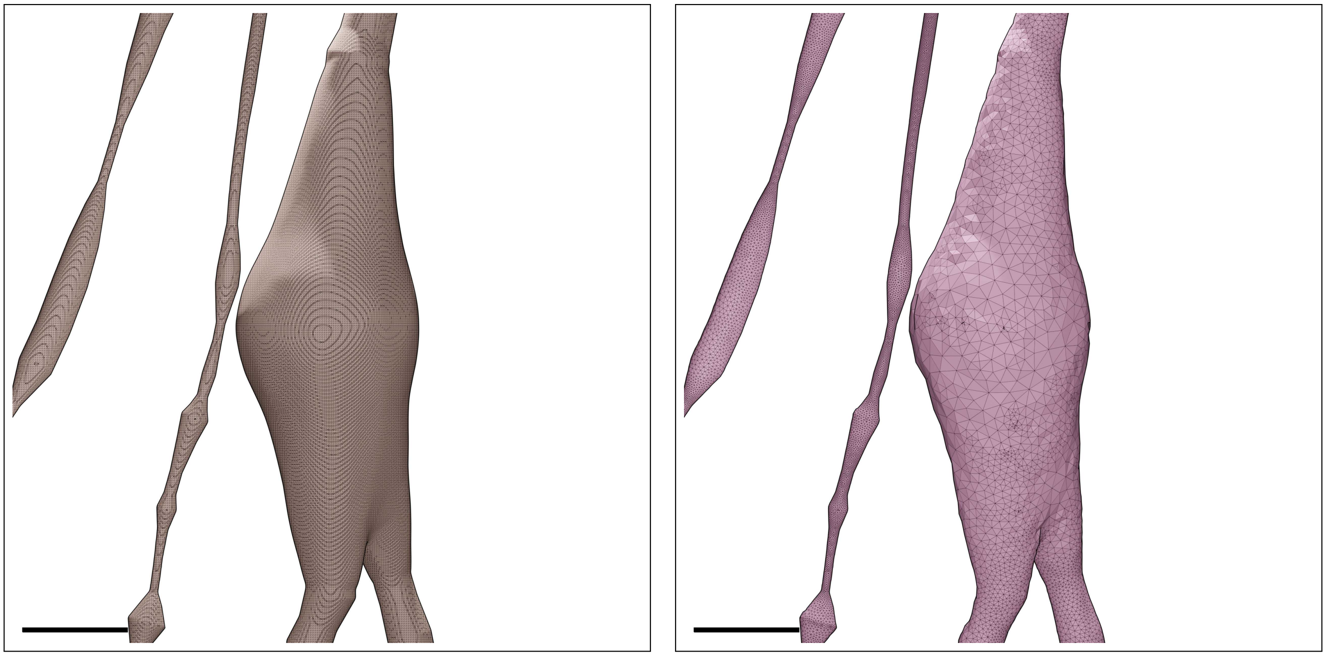

Figure S101: Wireframe visualizations of an L6_BTC neuron showing closeup comparisons between the highly tessellated surface mesh generated from the Voxel remesher (left) and the adaptively optimized surface mesh generated from the optimizer (right). Comparative quantitative and qualitative analyses of the meshes are demonstrated in Figure S102. Scale bars: 5 $\mu$m.

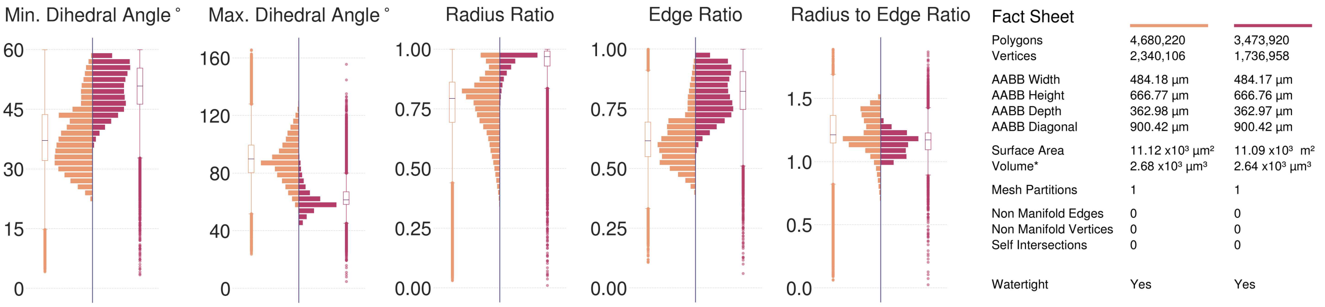

Figure S102: Comparative quantitative and qualitative analyses of the surface mesh models of the L6_BTC neuron visualized in Figure S101.

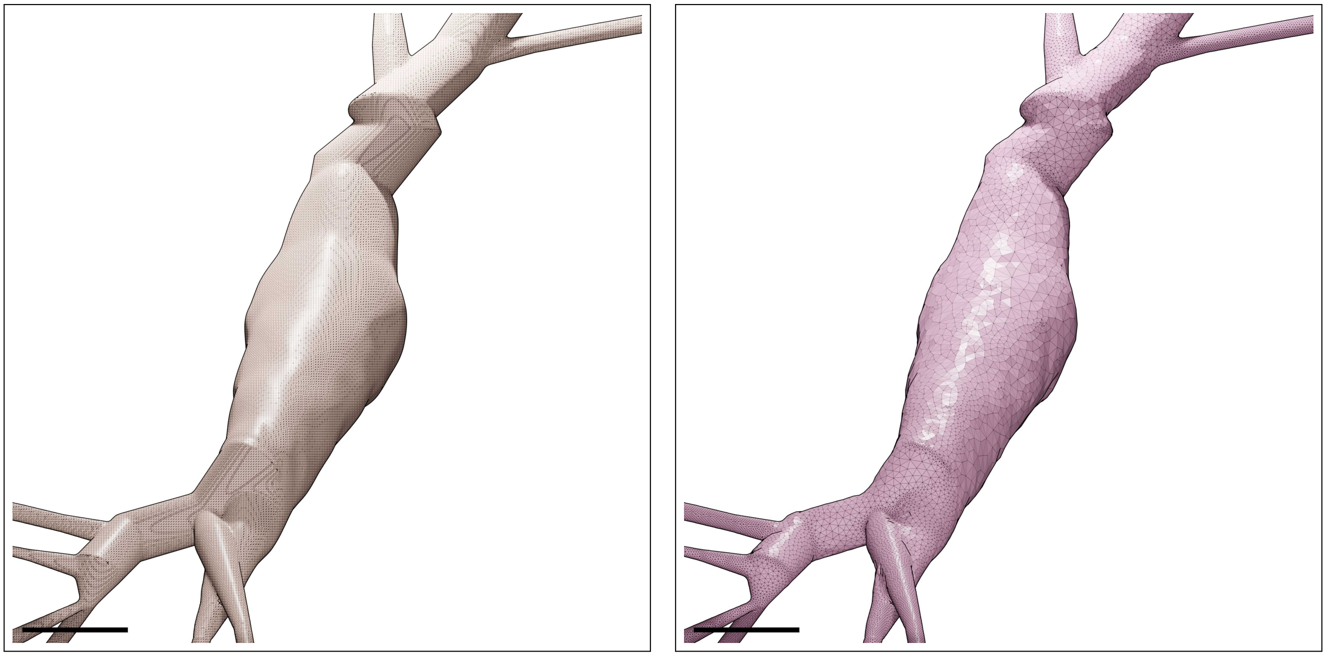

Figure S103: Wireframe visualizations of an L6_CHC neuron showing closeup comparisons between the highly tessellated surface mesh generated from the Voxel remesher (left) and the adaptively optimized surface mesh generated from the optimizer (right). Comparative quantitative and qualitative analyses of the meshes are demonstrated in Figure S104. Scale bars: 5 $\mu$m.

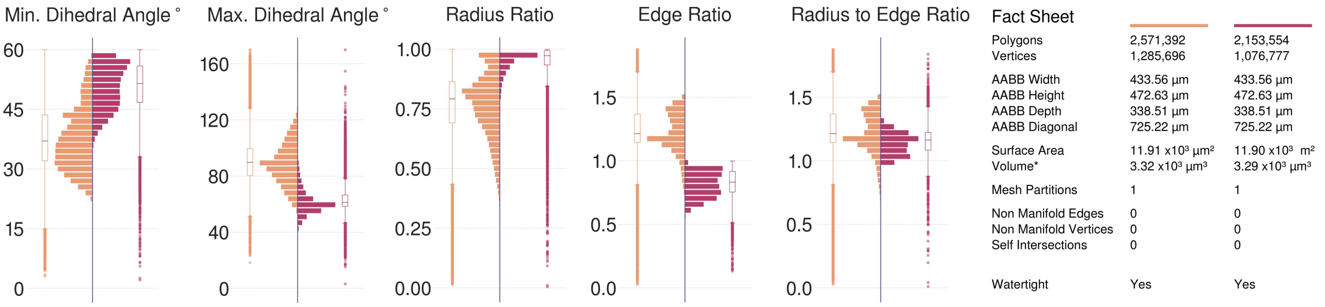

Figure S104: Comparative quantitative and qualitative analyses of the surface mesh models of the L6_CHC neuron visualized in Figure S103.

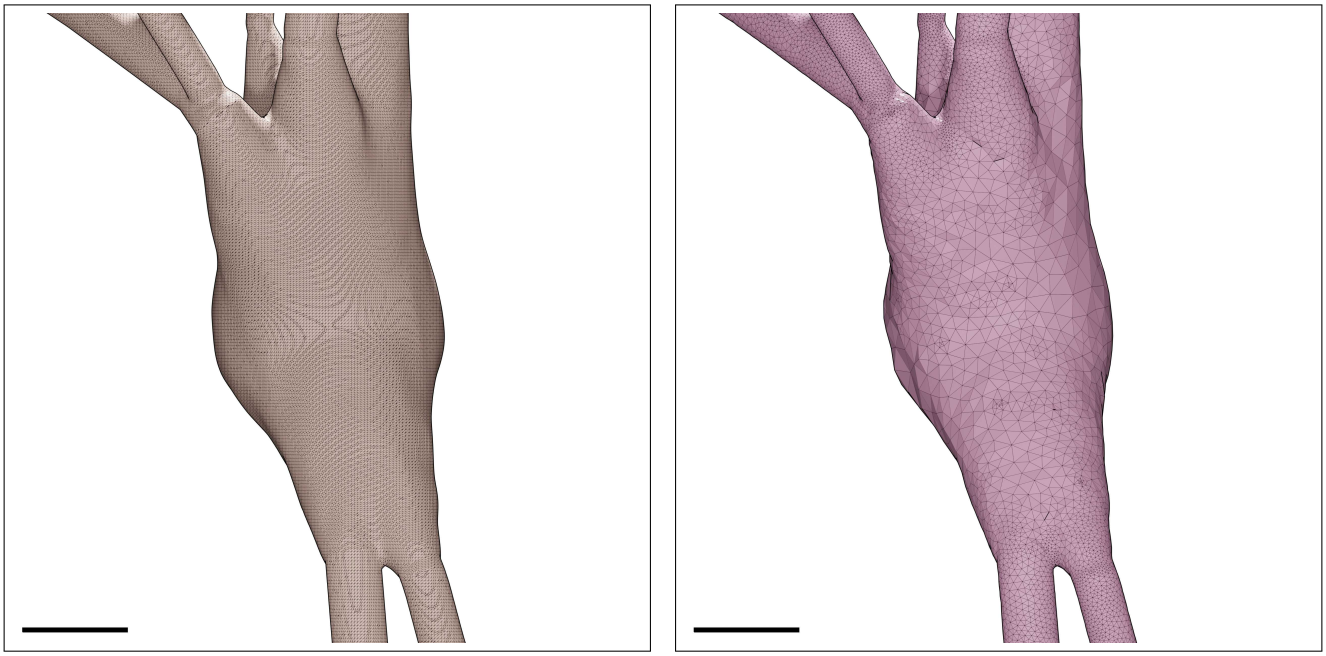

Figure S105: Wireframe visualizations of an L6_DBC neuron showing closeup comparisons between the highly tessellated surface mesh generated from the Voxel remesher (left) and the adaptively optimized surface mesh generated from the optimizer (right). Comparative quantitative and qualitative analyses of the meshes are demonstrated in Figure S106. Scale bars: 5 $\mu$m.

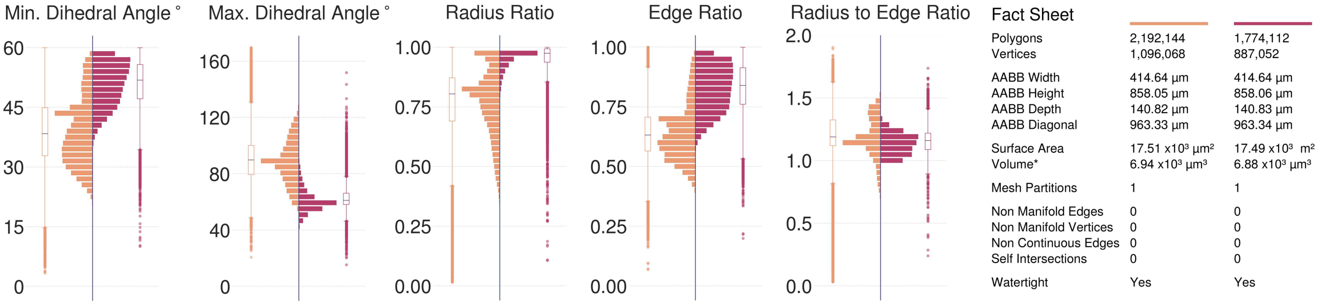

Figure S106: Comparative quantitative and qualitative analyses of the surface mesh models of the L6_DBC neuron visualized in Figure S105.

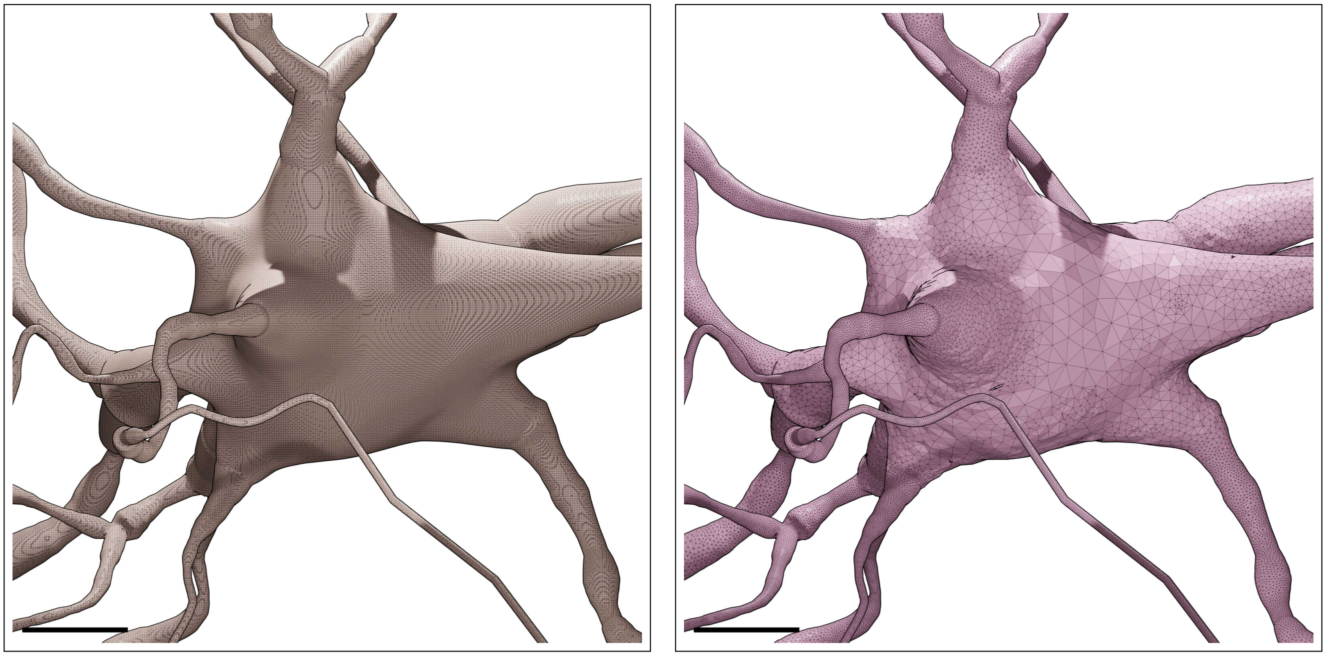

Figure S107: Wireframe visualizations of an L6_HPC neuron showing closeup comparisons between the highly tessellated surface mesh generated from the Voxel remesher (left) and the adaptively optimized surface mesh generated from the optimizer (right). Comparative quantitative and qualitative analyses of the meshes are demonstrated in Figure S108. Scale bars: 5 $\mu$m.

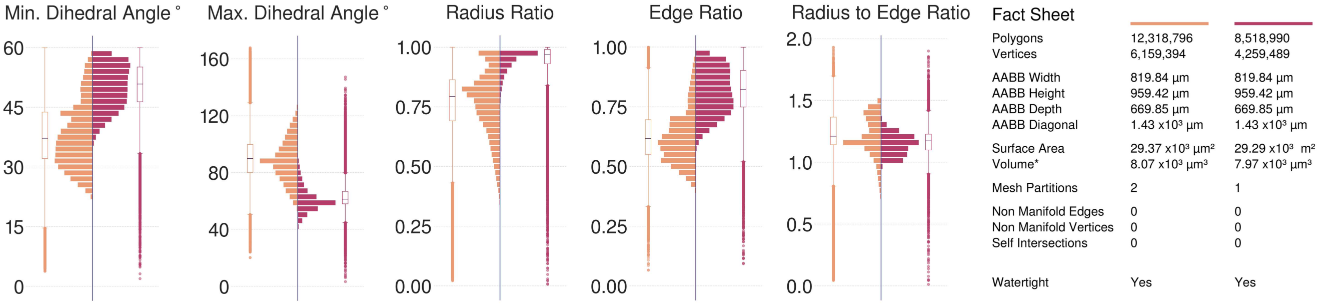

Figure S108: Comparative quantitative and qualitative analyses of the surface mesh models of the L6_HPC neuron visualized in Figure S107.

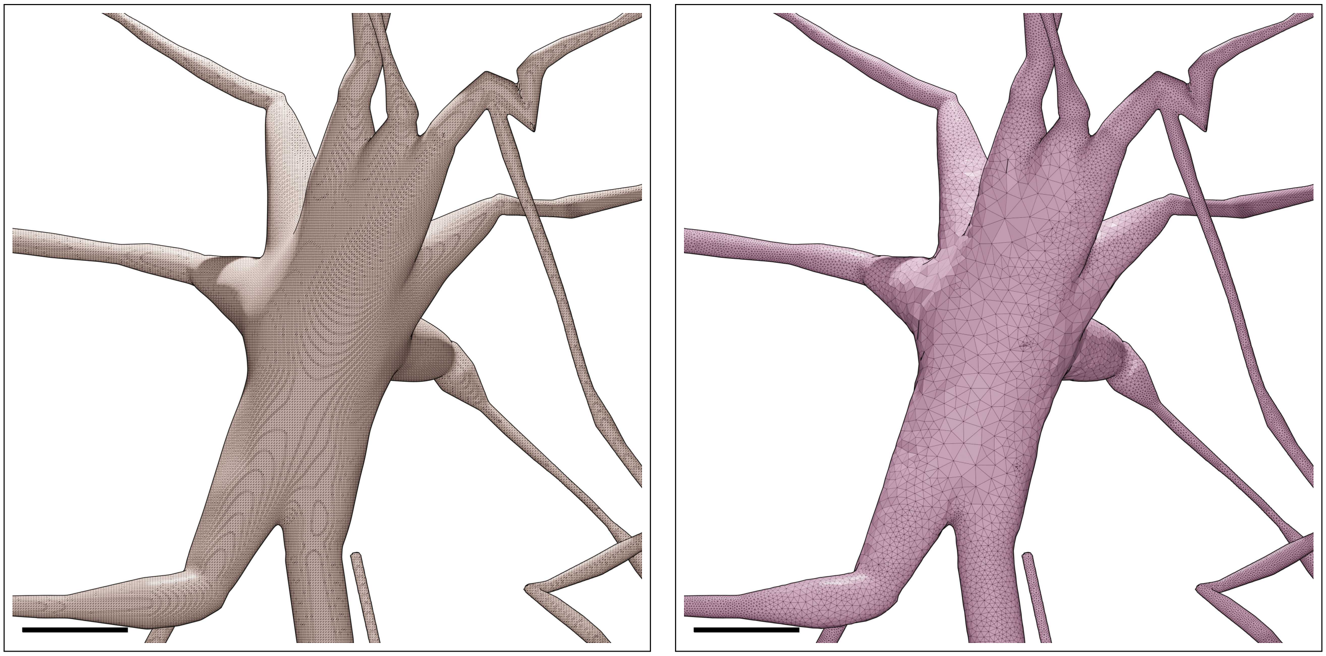

Figure S109: Wireframe visualizations of an L6_IPC neuron showing closeup comparisons between the highly tessellated surface mesh generated from the Voxel remesher (left) and the adaptively optimized surface mesh generated from the optimizer (right). Comparative quantitative and qualitative analyses of the meshes are demonstrated in Figure S110. Scale bars: 5 $\mu$m.

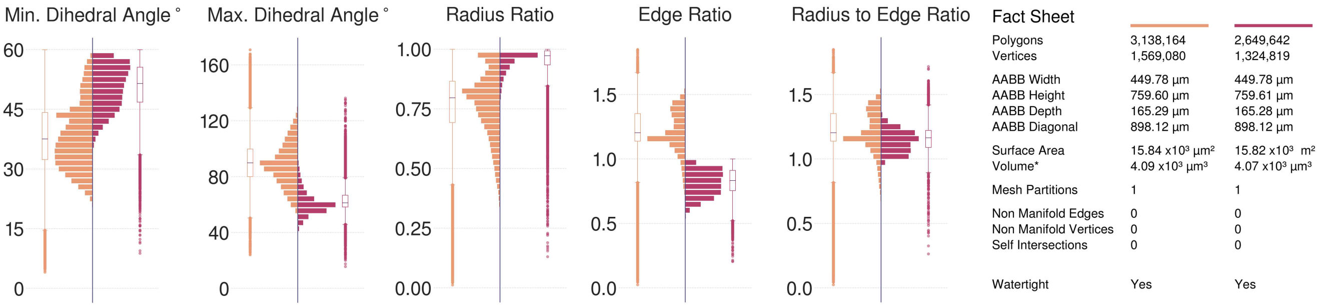

Figure S110: Comparative quantitative and qualitative analyses of the surface mesh models of the L6_IPC neuron visualized in Figure S109.

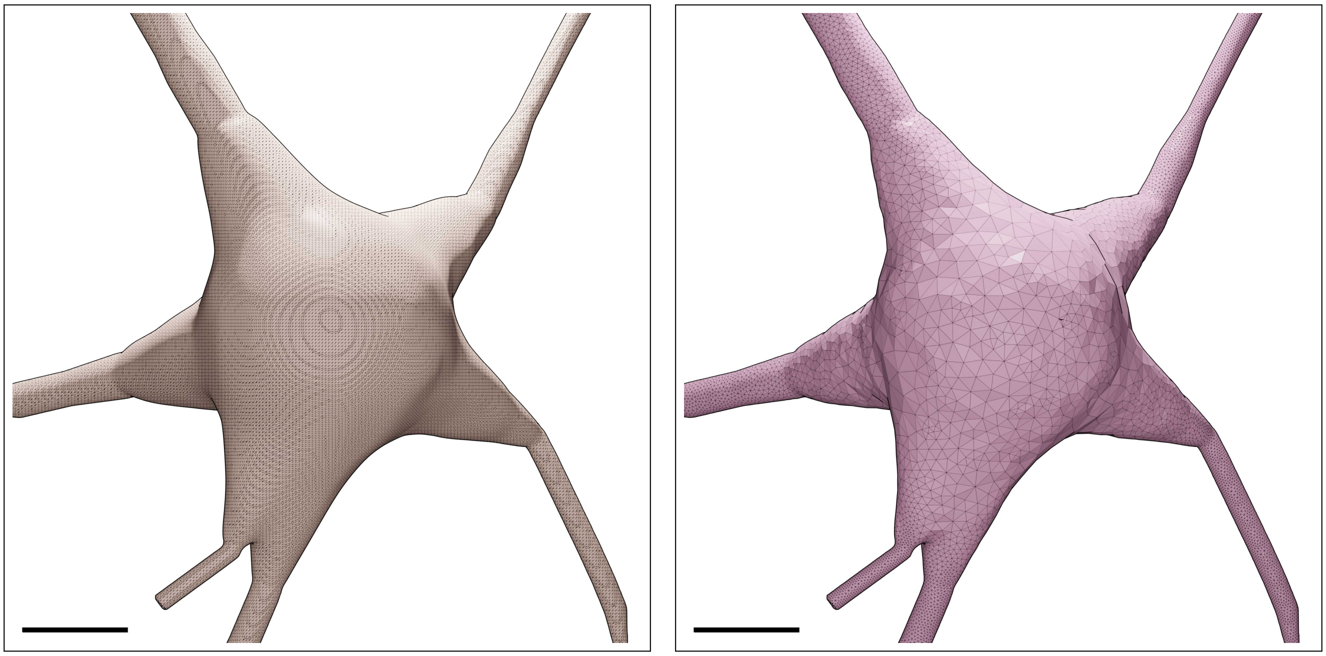

Figure S111: Wireframe visualizations of an L6_LBC neuron showing closeup comparisons between the highly tessellated surface mesh generated from the Voxel remesher (left) and the adaptively optimized surface mesh generated from the optimizer (right). Comparative quantitative and qualitative analyses of the meshes are demonstrated in Figure S112. Scale bars: 5 $\mu$m.

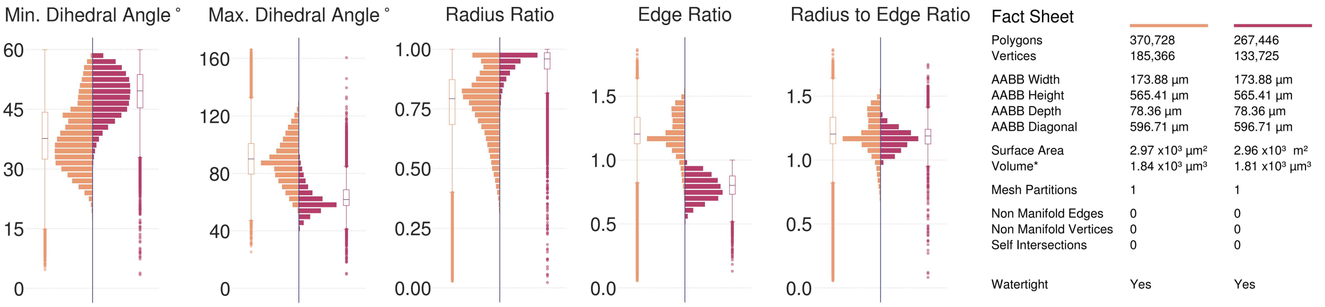

Figure S112: Comparative quantitative and qualitative analyses of the surface mesh models of the L6_LBC neuron visualized in Figure S111.

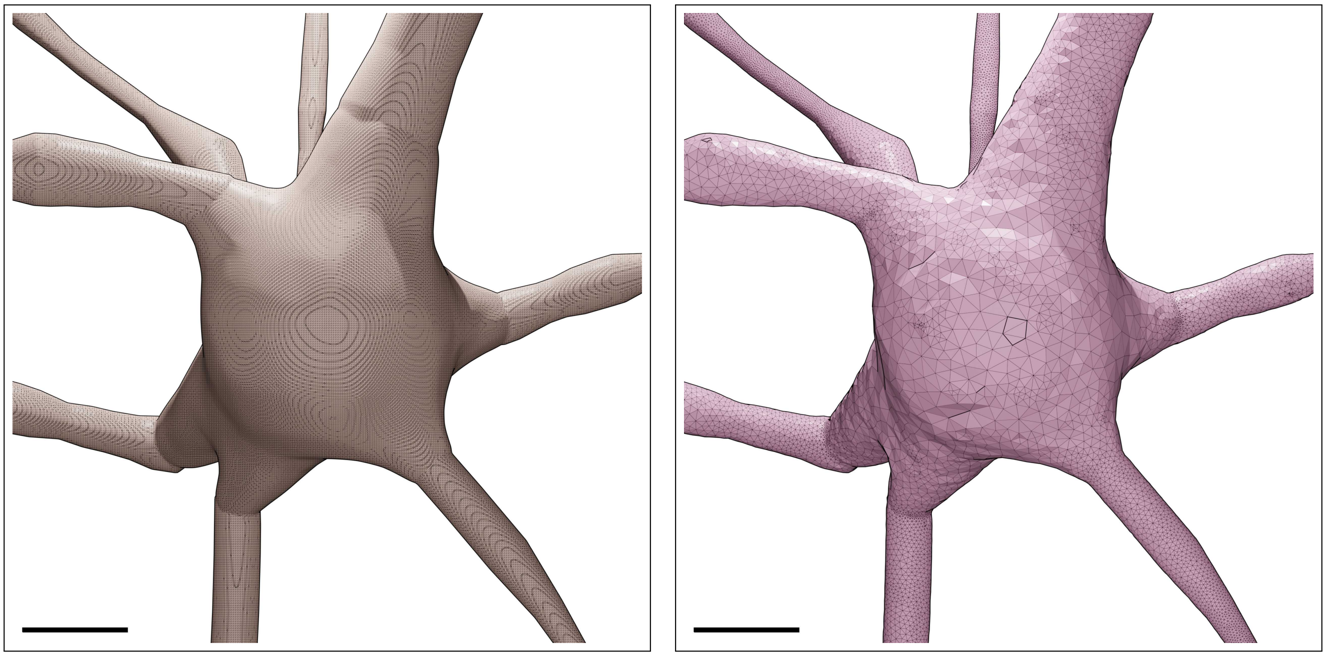

Figure S113: Wireframe visualizations of an L6_MC neuron showing closeup comparisons between the highly tessellated surface mesh generated from the Voxel remesher (left) and the adaptively optimized surface mesh generated from the optimizer (right). Comparative quantitative and qualitative analyses of the meshes are demonstrated in Figure S114. Scale bars: 5 $\mu$m.

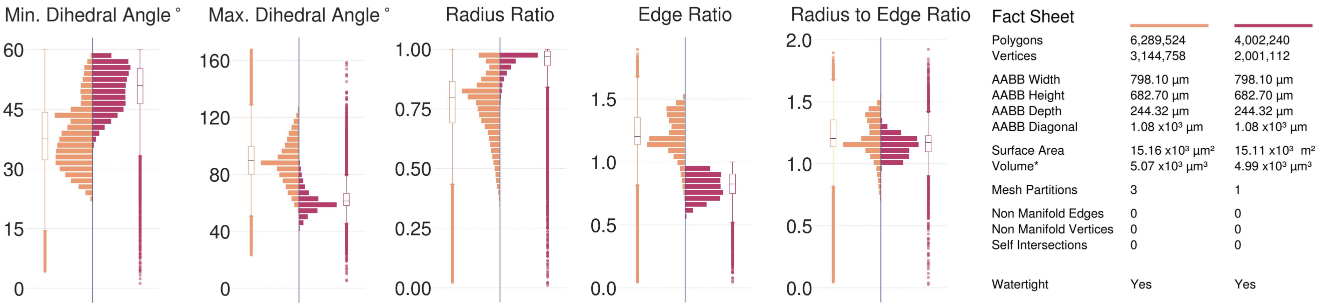

Figure S114: Comparative quantitative and qualitative analyses of the surface mesh models of the L6_MC neuron visualized in Figure S113.

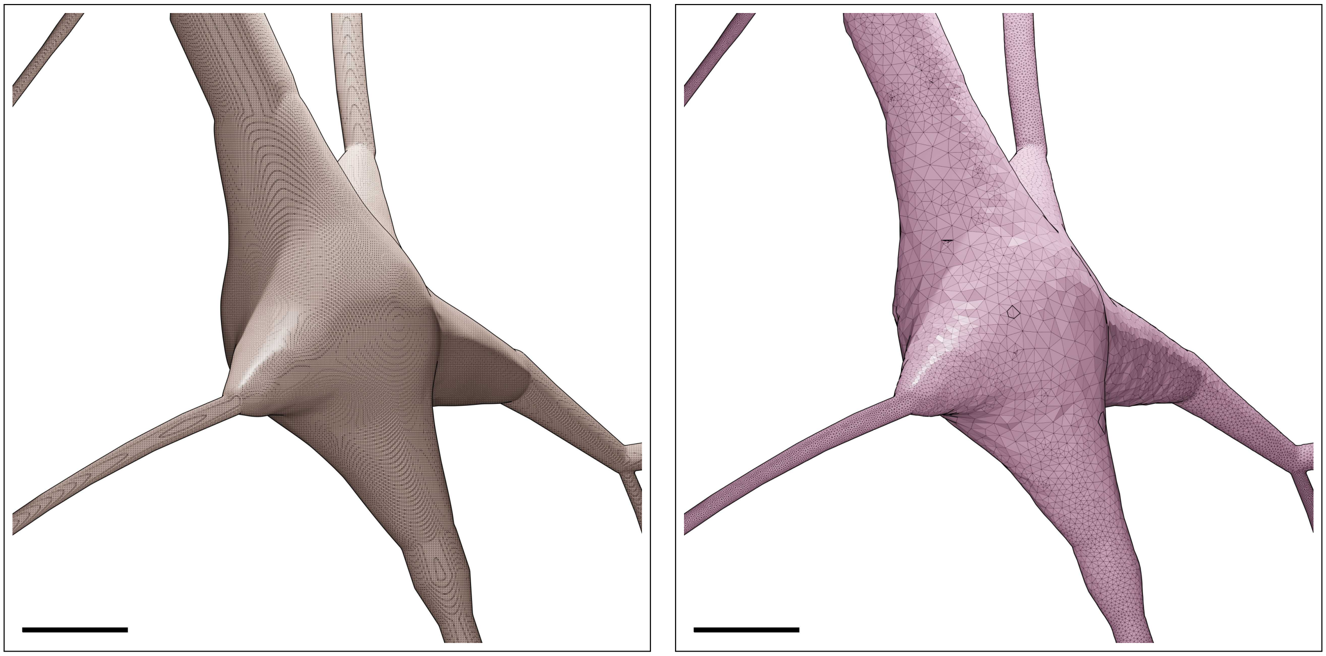

Figure S115: Wireframe visualizations of an L6_NBC neuron showing closeup comparisons between the highly tessellated surface mesh generated from the Voxel remesher (left) and the adaptively optimized surface mesh generated from the optimizer (right). Comparative quantitative and qualitative analyses of the meshes are demonstrated in Figure S116. Scale bars: 5 $\mu$m.

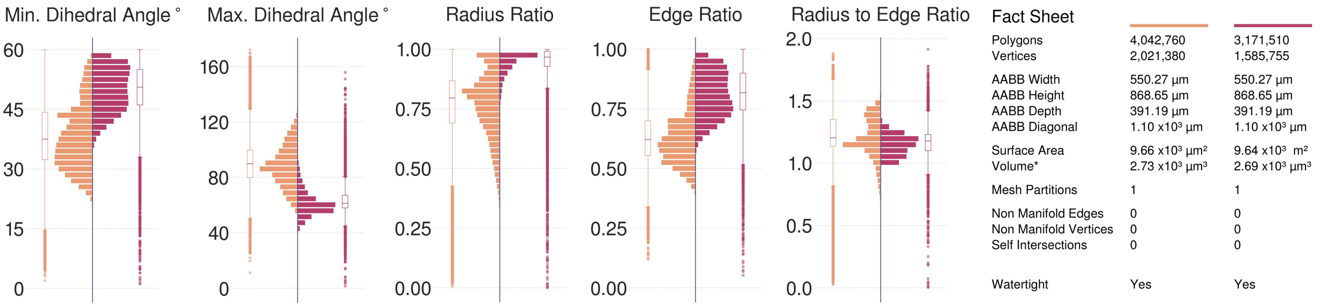

Figure S116: Comparative quantitative and qualitative analyses of the surface mesh models of the L6_NBC neuron visualized in Figure S115.

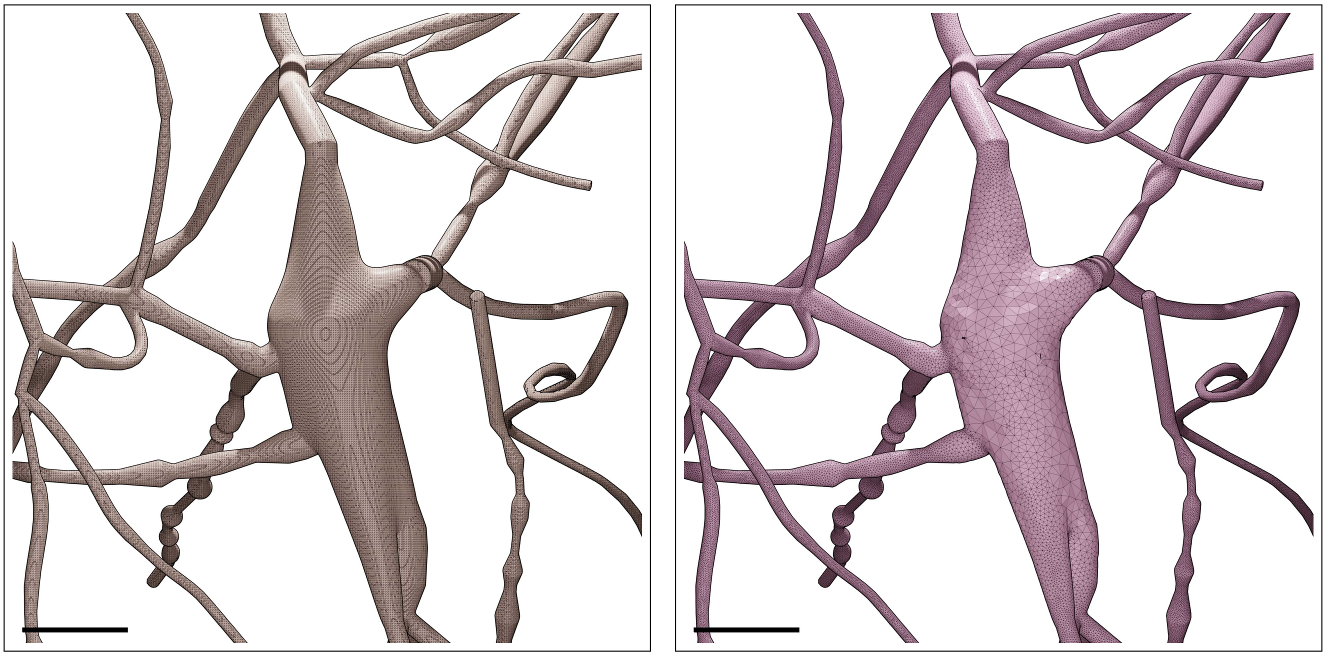

Figure S117: Wireframe visualizations of an L6_NGC neuron showing closeup comparisons between the highly tessellated surface mesh generated from the Voxel remesher (left) and the adaptively optimized surface mesh generated from the optimizer (right). Comparative quantitative and qualitative analyses of the meshes are demonstrated in Figure S118. Scale bars: 5 $\mu$m.

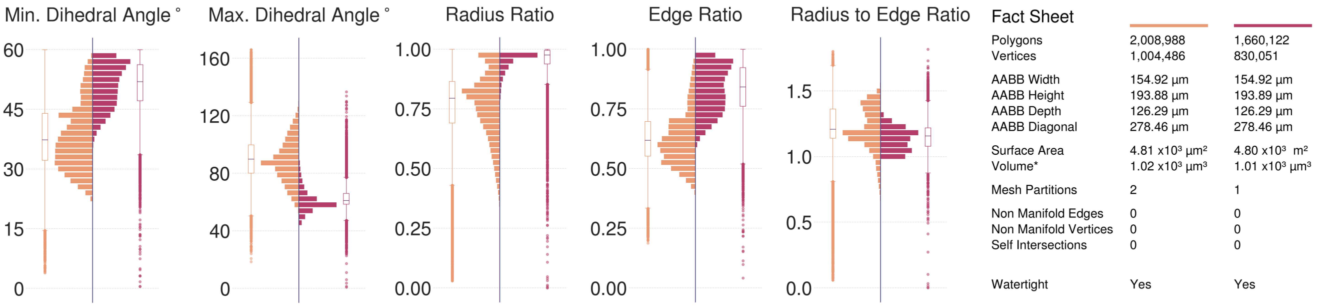

Figure S118: Comparative quantitative and qualitative analyses of the surface mesh models of the L6_NGC neuron visualized in Figure S117.

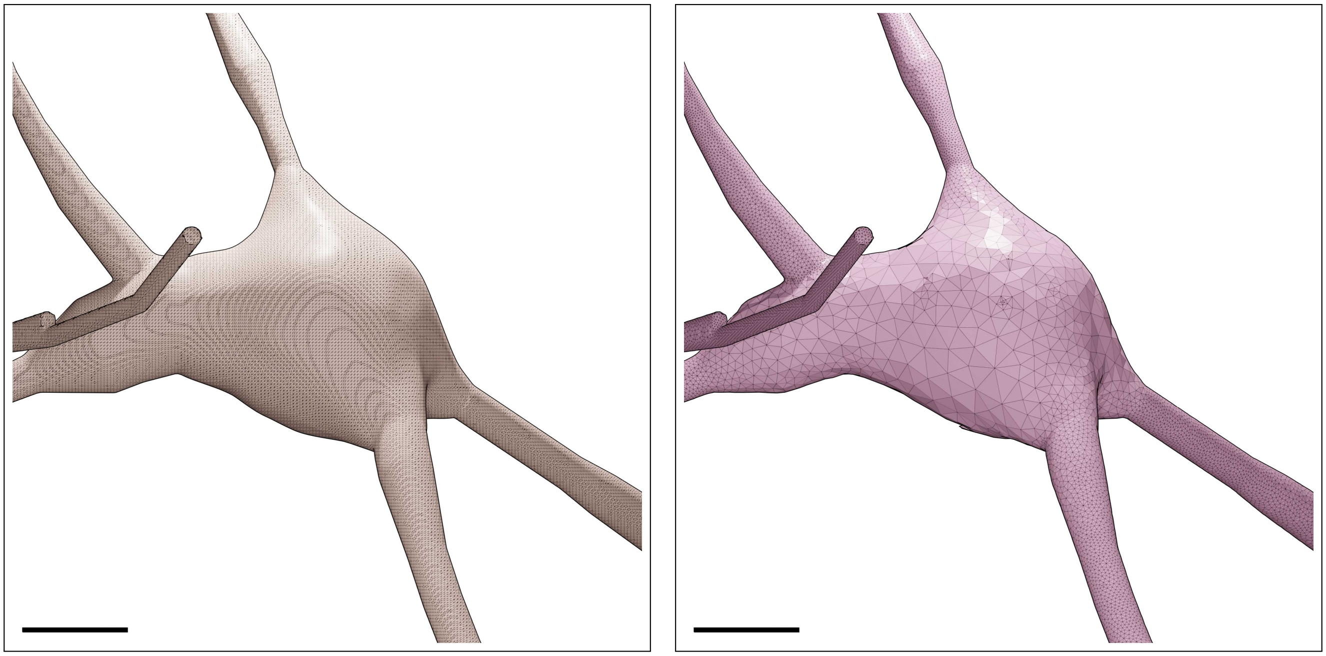

Figure S119: Wireframe visualizations of an L6_SBC neuron showing closeup comparisons between the highly tessellated surface mesh generated from the Voxel remesher (left) and the adaptively optimized surface mesh generated from the optimizer (right). Comparative quantitative and qualitative analyses of the meshes are demonstrated in Figure S120. Scale bars: 5 $\mu$m.

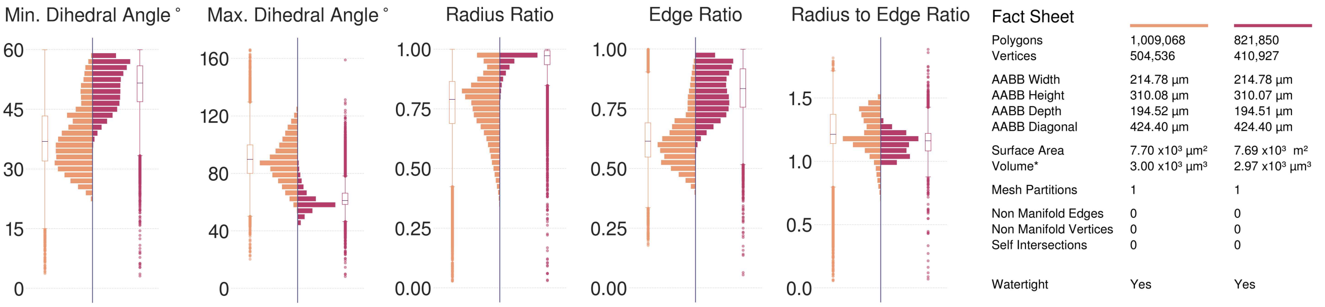

Figure S120: Comparative quantitative and qualitative analyses of the surface mesh models of the L6_SBC neuron visualized in Figure S119.

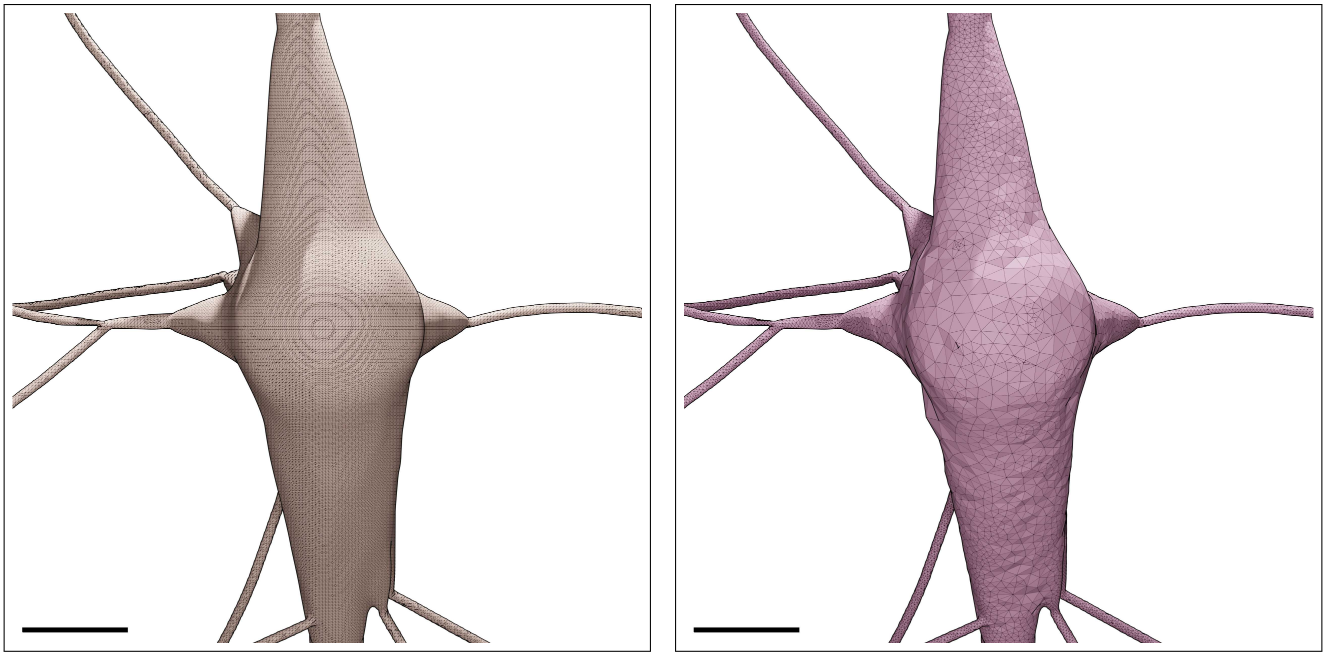

Figure S121: Wireframe visualizations of an L6_TPC:A neuron showing closeup comparisons between the highly tessellated surface mesh generated from the Voxel remesher (left) and the adaptively optimized surface mesh generated from the optimizer (right). Comparative quantitative and qualitative analyses of the meshes are demonstrated in Figure S122. Scale bars: 5 $\mu$m.

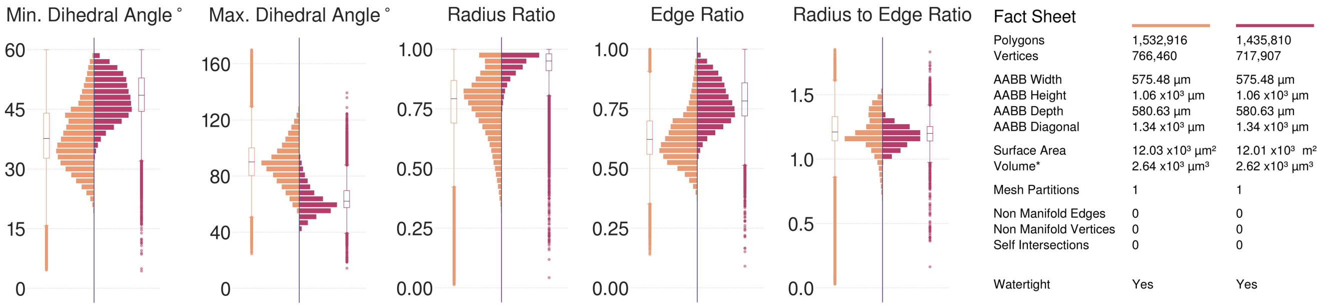

Figure S122: Comparative quantitative and qualitative analyses of the surface mesh models of the L6_TPC:A neuron visualized in Figure S121.

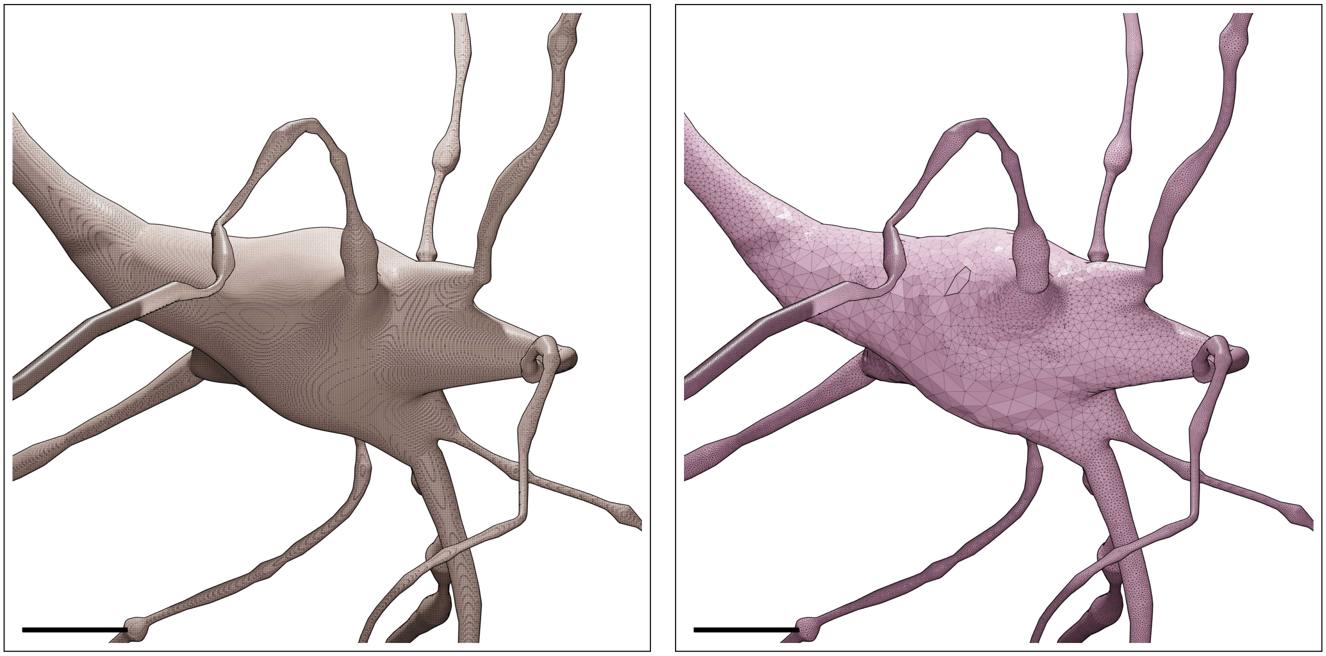

Figure S123: Wireframe visualizations of an L6_TPC:C neuron showing closeup comparisons between the highly tessellated surface mesh generated from the Voxel remesher (left) and the adaptively optimized surface mesh generated from the optimizer (right). Comparative quantitative and qualitative analyses of the meshes are demonstrated in Figure S124. Scale bars: 5 $\mu$m.

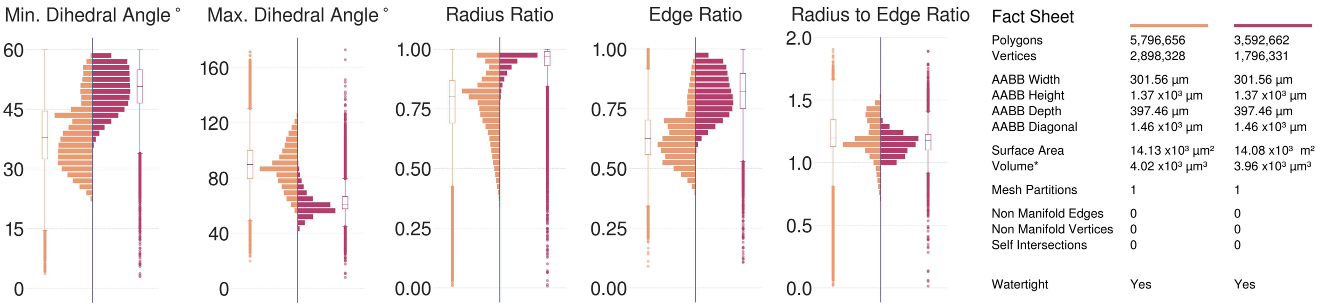

Figure S124: Comparative quantitative and qualitative analyses of the surface mesh models of the L6_TPC:C neuron visualized in Figure S123.

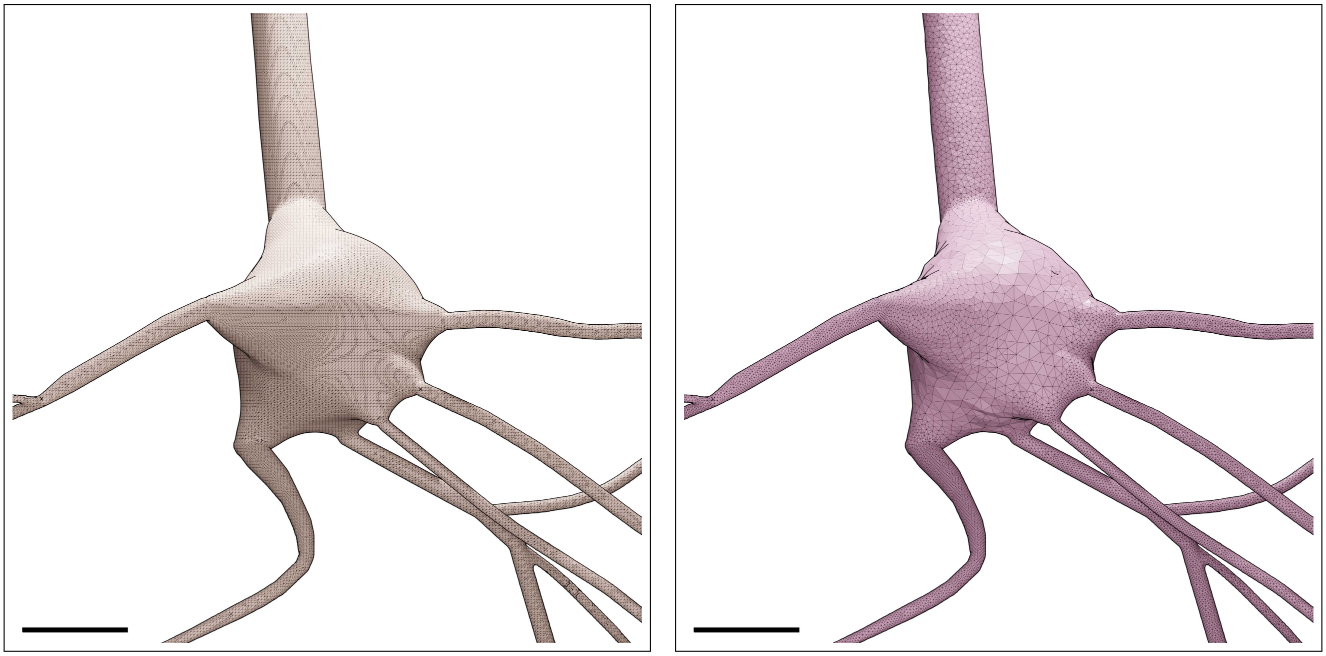

Figure S125: Wireframe visualizations of an L6_UPC neuron showing closeup comparisons between the highly tessellated surface mesh generated from the Voxel remesher (left) and the adaptively optimized surface mesh generated from the optimizer (right). Comparative quantitative and qualitative analyses of the meshes are demonstrated in Figure S126. Scale bars: 5 $\mu$m.

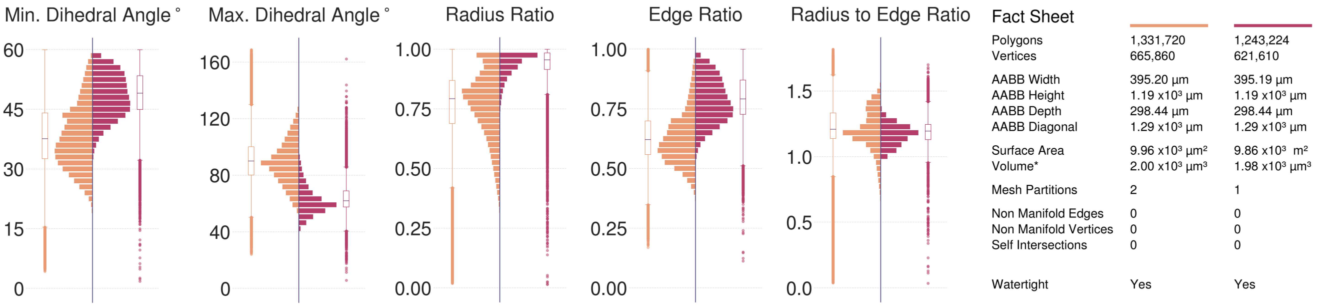

Figure S126: Comparative quantitative and qualitative analyses of the surface mesh models of the L6_UPC neuron visualized in Figure S125.

### 7 Comparative performance analysis

On average, and in comparison to previous meshing algorithms that are exclusively implemented in BLENDER, e.g.: skinning modifiers[12], implicit surface polygonization[13] and union boolean operators[14], our technique has a decent and scalable performance. This is evaluated by applying those BLENDER-based meshing techniques to a cortical pyramidal neuronal morphology[3], but with increasing branching orders (3, 4, 5 and 7). The comparative performance benchmarks are illustrated in Figure S127. It has to be noted that while the other approaches can generate models with realistic geometries as well, none of them is capable of achieving the watertightness criterion.

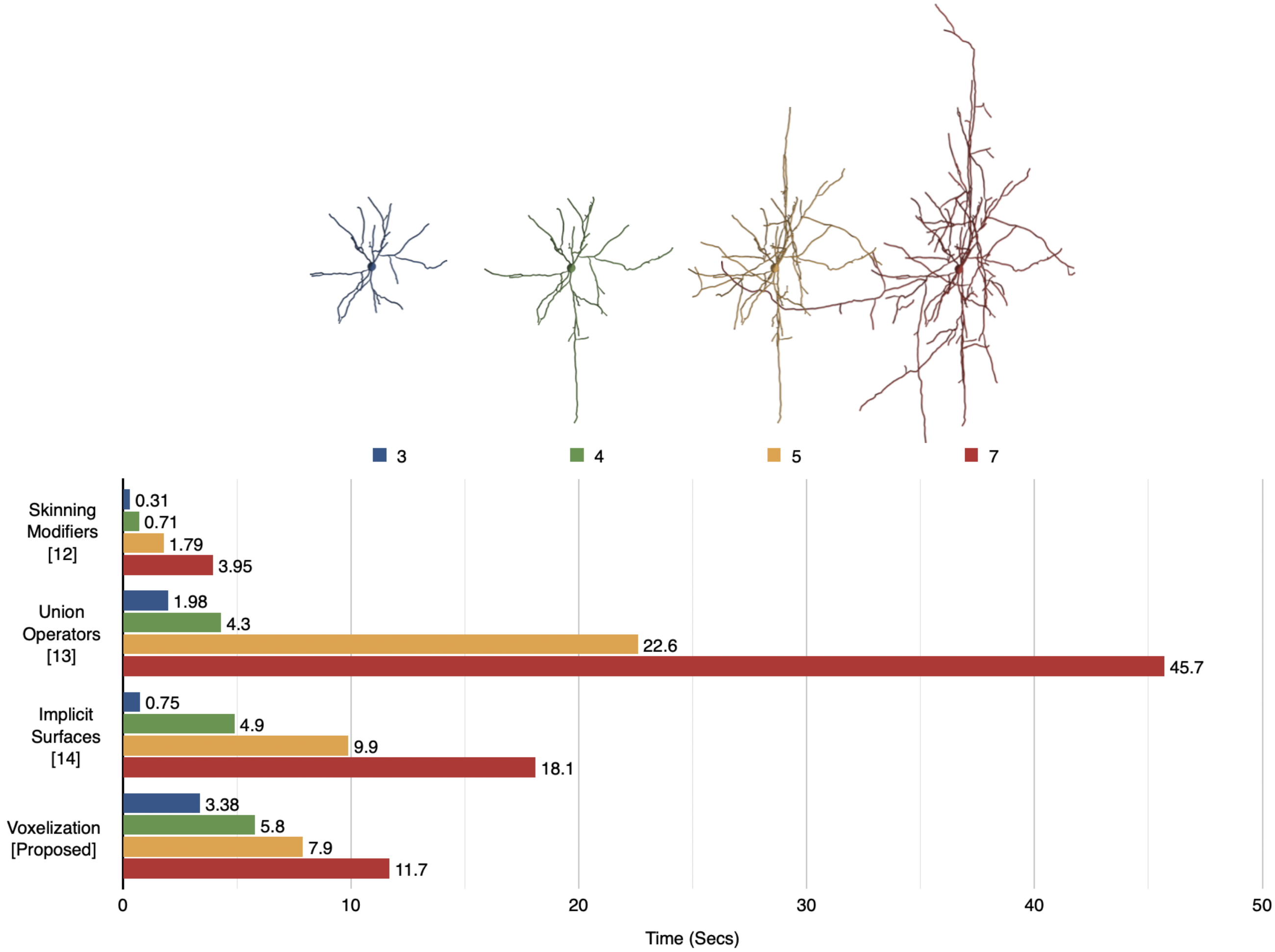

Figure S127: Comparing the performance of our proposed technique with other neuronal meshing techniques implemented exclusively in BLENDER using four morphologies of a pyramidal neuron, but with different branching orders as illustrated in the legends.

### 8 Software

#### 8.1 Code

The voxelization-based remeshing algorithm is implemented in BLENDER[9] based on its Python API. The technique is integrated within the Meshing Toolbox of the NEUROMORPHOVIS[15] add-on. The mesh optimization algorithms are implemeted in the OMESH – or OptimizationMesh– library. OMESH adapts and extends the GAMER – or Geometry-preserving Adaptive MeshER – library[10]. The optimization code is written in C++ and is integrated in NEUROMORPHOVIS[15] using Python bindings that are generated using pybind11[16].

#### 8.2 Software guide

To use our implementation to generate watertight surface manifolds of neuronal –or astrocytic– morphologies, users can install NeuroMorphoVis and select the Voxelization remesher in the Mesh Reconstruction Toolbox. If the OMESH bindings are located within the *libs* directory, resulting meshes will be automatically optimized. Otherwise, the user must compile it and copy the generated shared object to the *libs* directory. Users should use the same version of Python that is used by BLENDER. The following command should be used to install OMESH within NEUROMORPHOVIS.

```
BLENDER_PYTHON_VERSION setup.py build_ext install -prefix
PATH_TO_LIB_DIRECTORY
```

#### 8.3 Analysis code

The mesh analysis code is added to the *scripts* directory of NEUROMORPHOVIS.

#### 8.4 Complementary software

As we provide a full pipeline that takes input morphologies and creates optimized tetrahedral volumetric meshes for reaction-diffusion simulations, the following third-party software components are necessary to complement our software ecosystem. Note that our meshing implementation in NEUROMORPHOVIS requires the installation of BLENDER (at least version 3.0) to run the add-on.

1. BLENDER, which can be downloaded from https://www.blender.org.
2. TETGEN, which can be downloaded from https://wias-berlin.de/software/index.jsp?id=TetGen.
3. STEPs, which can be downloaded from https://steps.sourceforge.net/STEPS/default.php.

### 9 Supplementary data

Supplementary data including the resulting meshes of the 60 morphologies described in Table S1 and their analysis factsheets are available on Zenodo (https://doi.org/10.5281/zenodo.10558475).

### References

1. Botsch, M., Kobbelt, L., Pauly, M., Alliez, P. & Lévy, B. *Polygon mesh processing* DOI: 10.1201/b10688 (CRC press, 2010).
2. Markram, H., Muller, E., Ramaswamy, S., Reimann, M. W., Abdellah, M., *et al.* Reconstruction and simulation of neocortical microcircuitry. *Cell* **163,** 456–492. DOI: 10.1016/j.cell.2015.09.029 (2015).
3. Ramaswamy, S. *et al.* The neocortical microcircuit collaboration portal: a resource for rat somatosensory cortex. *Frontiers in neural circuits* **9.** DOI: 10.3389/fncir.2015.00044 (2015).
4. Si, H. & TetGen, A. A quality tetrahedral mesh generator and three-dimensional delaunay triangulator. *Weierstrass Institute for Applied Analysis and Stochastic, Berlin, Germany* **81** (2006).
5. Chen, W. *et al.* STEPS 4.0: Fast and memory-efficient molecular simulations of neurons at the nanoscale. *bioRxiv,* 2022–03. DOI: 10.3389/fninf.2022.883742 (2022).
6. Si, H. TetGen, a Delaunay-based quality tetrahedral mesh generator. *ACM Transactions on Mathematical Software (TOMS)* **41,** 1–36. DOI: 10.1145/2629697 (2015).
7. Wils, S. & De Schutter, E. STEPS: modeling and simulating complex reaction-diffusion systems with Python. *Frontiers in neuroinformatics* **3,** 15. DOI: 10.3389/neuro.11.015.2009 (2009).
8. Chen, W. & De Schutter, E. Parallel STEPS: large scale stochastic spatial reaction-diffusion simulation with high performance computers. *Frontiers in Neuroinformatics* **11,** 13. DOI: 10.3389/fninf.2017.00013 (2017).
9. Blender. *An open source 3D modelling and rendering package.* The Blender Foundation (Blender Institute, Amsterdam, 2024).
10. Yu, Z., Holst, M. J., Cheng, Y. & McCammon, J. A. Feature-preserving adaptive mesh generation for molecular shape modeling and simulation. *Journal of Molecular Graphics and Modelling* **26,** 1370–1380 (2008).
11. Coggan, J. S. *et al.* A process for digitizing and simulating biologically realistic oligocellular networks demonstrated for the neuro-glio-vascular ensemble. *Frontiers in neuroscience* **12.** DOI: 10.3389/fnins.2018.00664 (2018).
12. Abdellah, M., Favreau, C., Hernando, J., Lapere, S. & Schürmann, F. *Generating high fidelity surface meshes of neocortical neurons using skin modifiers* in *Computer Graphics and Visual Computing (CGVC)* (eds Vidal, F. P., Tam, G. K. L. & Roberts, J. C.) (The Eurographics Association, 2019). ISBN: 978-3-03868-096-3. DOI: 10.2312/cgvc.20191257.
13. Abdellah, M. *et al.* Metaball skinning of synthetic astroglial morphologies into realistic mesh models for in silico simulations and visual analytics. *Bioinformatics* **37,** i426–i433. DOI: 10.1093/bioinformatics/btab280 (2021).
14. Abdellah, M. *et al. Meshing of spiny neuronal morphologies using union operators* in *Computer Graphics and Visual Computing (CGVC)* (eds Vangorp, P. & Turner, M. J.) (The Eurographics Association, 2022). ISBN: 978-3-03868-188-5. DOI: 10.2312/cgvc.20221168.
15. Abdellah, M. *et al.* NeuroMorphoVis: a collaborative framework for analysis and visualization of neuronal morphology skeletons reconstructed from microscopy stacks. *Bioinformatics* **34,** i574–i582. DOI: 10.1093/bioinformatics/bty231. https://github.com/BlueBrain/NeuroMorphoVis (2018).
16. Jakob, W. *et al. pybind11 – Seamless operability between C++11 and Python (https://github.com/pybind/pybind11)* (2024).
